# BayesForge: A Bayesian Inference library for Python, R, and Julia

**DOI:** 10.64898/2026.01.19.700318

**Authors:** Sebastian Sosa, Mary Brooke McElreath, Cody Ross

## Abstract

1. Bayesian modeling is a cornerstone of modern ecological and evolutionary research, offering the flexibility to account for hierarchical structures, imperfect detection, and spatial dependencies. However, as ecological datasets grow in scale and complexity—from high-resolution telemetry to phylogenomics—researchers increasingly face a “computational ceiling” where traditional CPU-bound inference becomes prohibitively slow.
2. The current software landscape is fragmented. Researchers must often choose between high-level interfaces (e.g., *brms* in *R*) that are intuitive but sometimes rigid or slow for massive datasets, and low-level probabilistic programming languages (e.g., *Stan*, *PyMC*, *JAX*) that offer high performance but require specialized programming expertise.
3. This fragmentation is compounded by an “interoperability tax,” where code developed in one language (e.g., *R*) cannot easily leverage the hardware-accelerated backends (GPUs/TPUs) typically found in *Python*-centric machine learning frameworks.
4. To address these issues, we introduce *BayesForge(BF)*, a cross-platform software ecosystem available in *Python*, *R*, and *Julia*. *BF* provides a unified, intuitive syntax that bridges the gap between ease-of-use and high-performance computation. By leveraging *JAX*-based backends (*NumPyro* and *TensorFlow Probability*), *BF* enables seamless hardware acceleration under-the-hood.
5. We demonstrate *BF*’s utility through three ecological case studies: social network analysis (Social Relations Model), macroevolutionary uncertainty propagation across posterior tree sets, and latent-variable estimation for vocal repertoires. Benchmarks reveal that *BF* can achieve up to a 270-fold speedup over *Stan* implementations for large-scale networks, transforming weeks of computation into minutes and enabling more robust, uncertainty-aware ecological inference.

## 1 Introduction

Understanding how social bonds form and dissolve [1, 2, 3], how species assemble into complex communities while accounting for imperfect detection [4], or how animals navigate fragmented landscapes by inferring hidden behavioral states from GPS telemetry [5] requires explicit *generative models*: mechanistic representations of the micro-level processes that give rise to the patterns we observe [6]. That is, testing whether reciprocal relationships structure animal networks demands simulating the bond-formation process itself while accounting for unobserved interactions [7, 1, 8]. Similarly, estimating true species occupancy from imperfect detection records requires explicitly modeling the observation process to separate true absence from non-detection [9], and estimating individual vocal repertoire size demands accounting for the latent probability that a vocalization type exists but has yet to be recorded [10]. In each of these cases, researchers face the challenge of detection uncertainty—where the biological processes of interest are obscured by the imperfection of the observation process itself. To address such analytical problems, and many similar problems in evolution and ecology, researchers often need to build bespoke statistical models that encode domain knowledge about biological processes. The standard tools for building such models have recently been based on Bayesian inference packages. However, researchers attempting to implement these bespoke generative models confront a highly fragmented software landscape characterized by a steep accessibility-flexibility trade-off.

On the one hand, high-level interfaces such as *brms* [11] or *STRAND* [1] provide excellent accessibility for standard regression and network models in *R*, but their formula-based constraints severely limit the flexibility needed to build highly customized architectures. On the other hand, low-level Probabilistic Programming Languages (PPLs) like *BUGS* [12], *JAGS* [13], *Stan* [14], *PyMC* [15], *NumPyro* [16], and *TensorFlow Probability* [17] offer immense flexibility. Yet, they require users to master complex domain-specific languages or navigate low-level tensor mathematics, which can act as significant barriers to uptake by non-statisticians and those lacking in coding skills. Furthermore, because many of the most powerful computational engines are native to a single environment (predominantly *Python*), researchers working in *R* or *Julia* face translational friction. Accessing the capabilities of these powerful engines typically requires bridging tools or cross-language wrappers (e.g., *reticulate* [18], which further requires the manual conversion of *JAX* arrays between *Python* and *R*). This additional layer can steepen the learning curve even more, complicating cross-language error tracing, and interrupting dynamic analytical workflows.

To resolve this burden on research productivity, we introduce *BayesForge* (*BF*), a library designed specifically to maximize accessibility without sacrificing customizability. *BF* achieves this by allowing users to operate seamlessly across different levels of abstraction within the same modeling framework. Researchers can utilize highly optimized, pre-built functions for common complex structures (e.g., applying a Gaussian Process to account for spatial auto-correlation in habitat selection), while simultaneously using full mathematical notation to explicitly declare a novel biological mechanism, such as a custom likelihood function which accounts for biases in observation. This modular approach is supported by extensive, cross-disciplinary documentation detailing over 50 curated models—from social networks and phylogenetics to unsupervised clustering and Bayesian Neural Networks—all of which can be combined as needed to address complex causal inference problems. Crucially, *BF* is natively cross-platform, offering a standardized syntax across *Python*, *R*, and *Julia*. This eliminates language silos, allowing researchers to effortlessly transcribe existing models, collaborate seamlessly across disciplines, and operate entirely within their preferred programming environment.

Beyond accessibility, a structural bottleneck in modern Bayesian analysis is the often excessive computational time required. Massive datasets, such as complex social networks spanning several years of observation, or high-frequency animal tracking datasets, or simultaneous phylogenetic inferences across thousands of candidate trees, demand structured tensor computation at a scale that traditional Markov Chain Monte Carlo (MCMC) samplers cannot efficiently handle. *BayesForge* eliminates this barrier by acting as an accessible interface to modern, deep-learning-backed computation engines. By leveraging *JAX*, *NumPyro*, and *TensorFlow Probability*, *BF* inherits state-of-the-art computational efficiencies, including Just-In-Time (JIT) compilation, automatic differentiation, and massive parallelization and vectorization natively on hardware accelerators (GPUs and TPUs). This allows models that previously required days of computation to be fit in a matter of minutes.

Finally, *BayesForge* leverages its speed, minimalist syntax, and multiple layers of abstraction to facilitate a complete, end-to-end Bayesian workflow. Because ecological data is notoriously noisy and biased, principled Bayesian data analysis must extend far beyond a single posterior inference fit [19]. A robust analysis requires an iterative *Bayesian workflow* that includes: prior predictive simulations (to ensure priors do not imply biologically absurd outcomes, such as impossible animal growth rates), data simulations to facilitate parameter recovery tests (to confirm that a bespoke model is well specified and permits recovery of true parameters from empirically possible data sets, like sparse telemetry arrays), simulation-based calibration (to guarantee that credible intervals for sensitive metrics like extinction risk are statistically valid and not overconfident), and posterior predictive checking (to verify that the fitted model accurately reproduces observed structural patterns, such as the massive excess of zero-counts typical in camera trap surveys). In contemporary software landscapes, executing these steps introduces implementation overhead, often forcing users to write Data Generating Models (DGMs) in one language (e.g., *R*) and Inference Models (IMs) in another (e.g., *Stan*). Furthermore, delegating iterative simulation steps to non-optimized code creates computational bottlenecks, making the full workflow validation practically infeasible. *BF* unifies the entire analytical workflow. By treating generative simulation, inference, and calibration as element of a dedicated object-class governed by the same underlying hardware-accelerated computational engine, *BF* ensures that rigorous, computationally heavy validation pipelines are fast and routine. By lowering the barrier to comprehensive model evaluation, *BayesForge* empowers researchers to routinely execute these essential validation pipelines and fit biologically realistic models, transforming best-practice workflows from a computational challenge into an accessible standard.

## 2 Software Presentation

*BayesForge* (*BF*) is structured around a unified, object-oriented Application Programming Interface (API) that maximizes cross-disciplinary **accessibility**. The API remains structurally consistent across *Python*, *R*, and *Julia*. While some minor unavoidable syntactic differences exist to adhere to the idiomatic conventions of each language, the core model specification syntax, the procedural workflow for analysis, and the underlying computational engines remain identical.

To streamline the *Bayesian workflow*, users interact with a primary *BF* object to sequentially execute an end-to-end analysis. This object features subclasses for *handling data*, *defining a model*, *runing inference*, *analyzing posterior samples*, and *visualizing substantive results*. By standardizing this procedural pipeline natively across languages, *BF* eliminates the need to juggle disparate software tools. Crucially, it bridges the gap between methodological development and applied research: a state-of-the-art generative model designed by a quantitative methodologist in *Python* can be easily transcribed and deployed by an applied ecologist more familiar with *R* or *Julia*. This ensures that cutting-edge probabilistic architectures are immediately accessible to the broader community, without requiring ecologists to learn a new programming language or manually translate complex mathematical syntax into their preferred language—a process highly susceptible to implementation errors.

To navigate the *accessibility-flexibility trade-off*, *BF* provides a comprehensive, fully-documented library of over 50 models (see Table 1). Rather than functioning as rigid, black-box equations, these implementations serve as modular building blocks implemented directly in *JAX*. Many key modeling architectures used in evolutionary biology and ecology are provided out-of-the-box—examples of which include the *Social Relations Model (SRM)* for inferring reciprocal animal interactions [1], *Network-Based Diffusion Analysis (NBDA)* for tracing the social transmission of behaviors [7], and phylogenetic comparative methods for macroevolutionary trait mapping [20]. These are provided alongside general-purpose statistical components, including *varying effects* (e.g., for modeling repeated measurements or quantifying individual variation in animal personality studies), *Gaussian Process kernels* (e.g., to control for continuous spatial autocorrelation in habitat selection [5]), *survival models* (e.g., estimating time-to-event dynamics and mortality risk), and more novel architectures, such as *Dense Neural Network Layers* for highly nonlinear environmental regressions or bioacoustic signal classification.

**TABLE 1.** Comprehensive list of models available in *BayesForge* and their ecological applications.

| Documented Model | Example Ecological Application | Documented Model | Example Ecological Application |
| --- | --- | --- | --- |
| Linear Regression [6] | Island Species-Area Relationship [25] | Phylo. meta-analysis [20] | Relationship between species habitats and body sizes [26] |
| Multiple Regression [6] | Joint effects of temperature and rainfall on foraging [27] | Phylo. varying slopes [20] | Species' responses to climate change [28] |
| Interactions [6] | Predator presence altering resource availability effects [29] | Measurement-error [6] | Correcting for known inaccuracies in sensors [30] |
| Categorical Models [6] | Comparing clutch sizes across discrete habitat types [31] | Missing-data Models [6] | Animal population surveys [32] |
| Binomial Models [6] | Fledgling survival probability from a known clutch size [33] | Latent variable models [34] | Inferring hidden species interactions and unmeasured environmental data [35] |
| Poisson Models [6] | Observed species counts in quadrat surveys [36] | PCA [37] | Ancient human migration and language diversification [38] |
| Poisson with offset [6] | Biomass estimation [39] | Gaussian Mixtures [40] | Simultaneous tracking of changes in microalgae diversity and community structure [41] |
| Gamma-Poisson [6] | Clustered parasite loads [42] | Dirichlet Mixtures [43] | Estimating an unknown number of ecological niches [44] |
| Categorical [6] | Anthropogenic effects on biological communities [45] | Social Relations Model [1] | Grooming behavior among baboons [46] |
| Multinomial Models [6] | Estimation of population structure [47] | Stochastic Block Model [1] | Identifying latent community clustering in food webs [48] |
| Dirichlet models [49] | Proportional dietary composition from metabarcoding [50] | Network bias control [8] | Adjusting for sampling biases in network inference [51] |
| Beta-binomial [6] | Ecological richness estimation [52] | Network Edges [2] | Benefits of sociality on health [53] |
| Zero-inflated [6] | Distinguishing structural and random zeros in parasite contagion [54] | Network Metrics [55] | Influence of individual characteristics on sociality [56] |
| Varying-intercept [6] | Unobserved site-level variation in multi-site monitoring [57] | NBDA [58] | Social transmission of novel foraging behaviors [59] |
| Varying-effects [6] | Individual behavioral plasticity [60] | Weighted Attraction [61] | Kin-based or phenotype-based learning preferences [61] |
| Nested effects [6] | Genetic traits variation between multiple sites [62] | BNN [16] | Spatiotemporal prediction for air pollution [63] |
| Gaussian process [6] | Modeling animal movement [64] | BNN Classification [65] | Animal Species Recognition [66] |
| Phylogenetic analysis [20] | Quantify conservative trait correlation [67] | BNN Multiclass [65] | Cancer classification [68] |
| Phylogenetic repeated measures [20] | Species overlap and phylogenetic relatedness [69] | Survival Analysis [15] | Influence of sociality on animal fitness [70] |

Note: All models act as modular components that can be customized and combined. Full mathematical documentation and implementation snippets in R, Python, and Julia are available in the package repository.

Crucially, because all components share a common mathematical backend, users retain the flexibility to easily combine them. For example, a researcher can construct a zero-inflated Poisson model with nested varying effects to account for structural zeros in camera-trap biodiversity surveys [21], or build a joint model where principal components (derived from a Bayesian PCA of shifting climatic variables) serve as predictors in a subsequent survival analysis. This modularity allows measurement error and uncertainty to be mathematically propagated through all stages of the analysis. Each component is supported by extensive documentation detailing the underlying mathematical assumptions and providing direct code snippets in *Python*, *R*, and *Julia*.

At its computational core, *BF* delivers unprecedented *scalability* by bypassing the bottlenecks of traditional MCMC samplers. It delegates all heavy tensor operations to modern, deep-learning-backed PPLs, specifically *NumPyro* and *TensorFlow Probability*. This architecture provides automatic differentiation, Just-In-Time (JIT) compilation, and native hardware acceleration (GPU/TPU) entirely under the hood. By abstracting these low-level backend complexities into a high-level, language-agnostic API, *BF* brings state-of-the-art computational *scalability* to applied researchers. This empowers ecologists to fit models that where once computationally prohibitive—e.g., tracking disease transmission across densely connected spatial matrices [22]—without requiring deep expertise in computer science or tensor mathematics.

## 3 Example 1 Generative Model: Social Relations Model

As discussed, *BF* is useful for complex generative modeling across disciplines. We show this here regarding social network models that are commonly used in modeling animal social behaviors. More specifically, we give an example of a generative implementation of the *Social Relations Model (SRM)* (Box 1), where the *BF* package is used not only to fit observed data, but to define the features of the underlying generative process—such features include latent individual propensity to give (e.g., give food), individual propensity to receive (e.g., receive food), and dyadic reciprocity. This framework provides a “theoretical playground” where researchers can simulate how micro-level individual traits give rise to macro-scale network topologies or system-wide organizational structures. By abstracting the dense matrix algebra and hierarchical interdependence associated with the SRM into a simplified level of abstraction, *BF* allows scholars to bridge the gap between defining individual-level processes and emergent phenomena without the technical overhead typically associated with custom probabilistic programming. This transition from descriptive to generative modeling ensures that sophisticated theory-testing remains accessible to researchers regardless of their computational background or primary field of study.

## 4. Example 2 Phylogenetic analysis: Navigating Evolutionary Landscapes

We demonstrate *BF*’s capacity for *vectorized parallelization* by running a comparative phylogenetic analysis coupled with a measurement error model [6]. In the era of big-data phylogenetics, common reliance on a single consensus tree often masks the potentially substantial level of uncertainty in branch lengths and gross topology by collapsing conflicting genealogical signals into a single bifurcating topology, thereby giving an illusion of high confidence that obscures underlying gene tree discordance [23]. By loading an credible set of tree posteriors [24] as a unified 3D tensor, *BF* facilitates the seamless analysis of the entire parameter space of phylogenetic uncertainty, allowing macroevolutionary traits and diversification rates to be computed across thousands of alternative topologies.

### Box 1. SRM generative model

1. **Simulation Process:** The generative process follows these steps: (1) initializing nodal and dyadic covariates *x* and *d* (Eqs. 1a, 1b), (2) defining the sampling effort, *E*, for each tracked individual (Eq. 1c), and therefore each dyad, *E*^ (Eq. 1d), (3) incorporating nodal, *λ*, *π* (Eq. 1e) and dyadic, *δ* (Eq. 1f) varying effects, (4) defining a linear model for the log-odds of a tie, *η* (Eq. 1g), and (5) generating the final, observation-level, network links, *Y* (Eq. 1h).

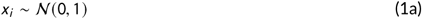

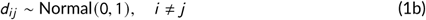

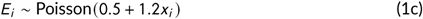

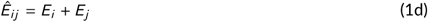

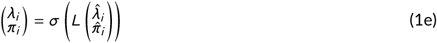

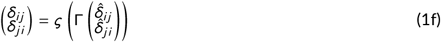

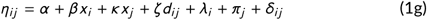

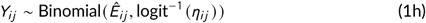

Note that in Eq. 1g, *α* is an intercept parameter, *β* and *κ* are slopes parameters on the individual covariate *x*, and *ζ* is a slope parameter on the dyadic covariate *d*. Lastly, *σ* and *ς* are the scale parameters for the varying effects, and *L* and Γ are Cholesky factors of the correlation matrices of the varying effects.
2. **Inference Model:** A complete, annotated implementation, including a decomposition of the fixed and varying effects for nodal and dyadic effetcs for the *Generative Model*, as well as the *Inference Model* and posterior validation against the generative process, is provided in *Appendix 1*.
3. **BI Implementation:**

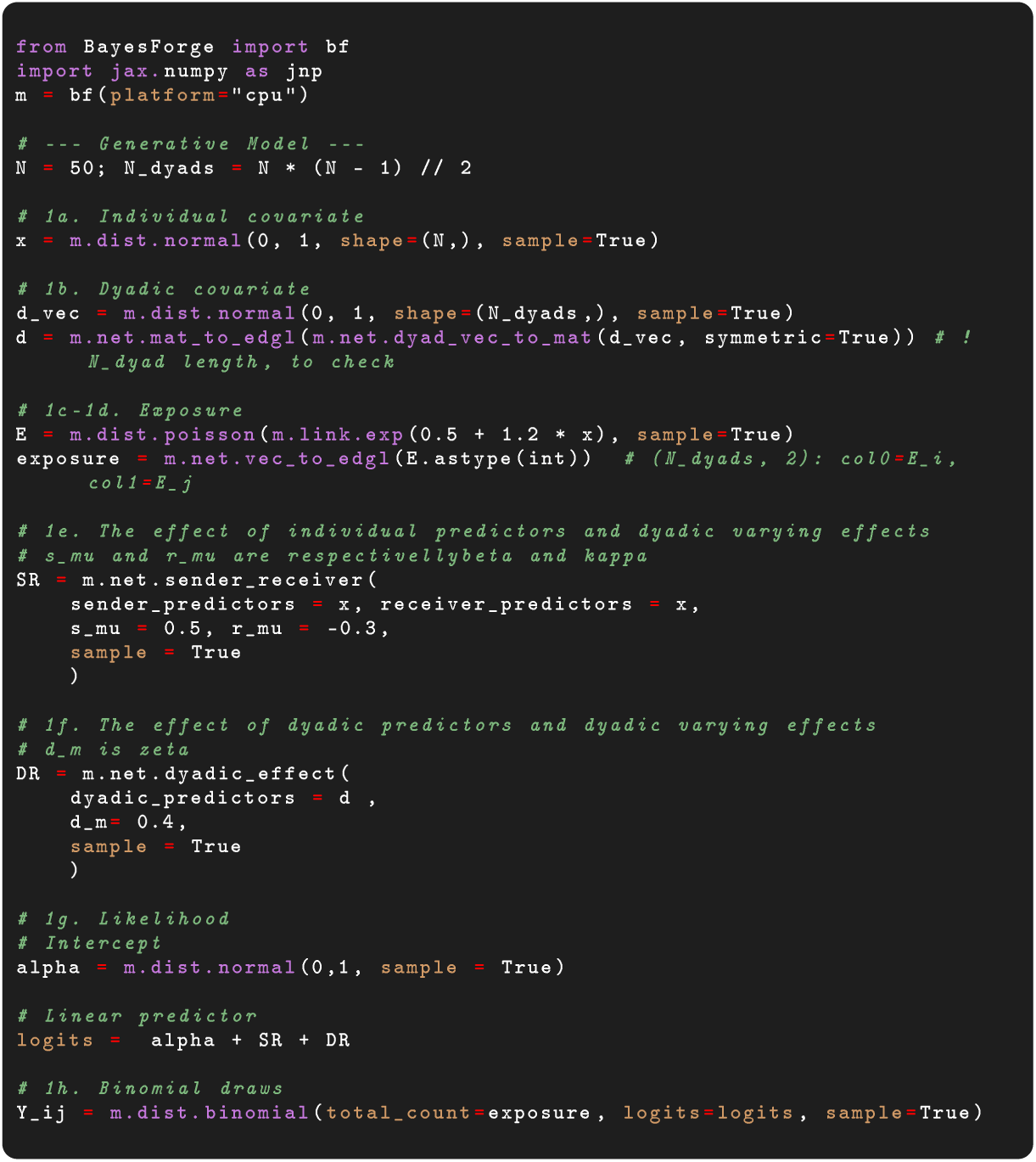

### Box 2. Phylogenetic Generalized Mixed Model with posterior tree samples

1. **Inference model:** Trait evolution across species is modelled as a linear regression that accounts for shared ancestry via a phylogenetic random effect with the phylogenetic covariance matrix derived from a posterior distribution of *T* trees to propagate phylogenetic uncertainty (Eq. 2):

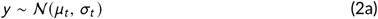

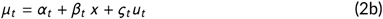

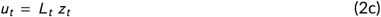

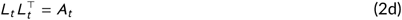

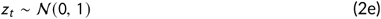

where *t* ∈ {1, …, *T* } indexes the sampled phylogenetic tree. In this example, x and y are the log-transformed, mean-centred predictor (body mass) and response (body length), respectively. The matrix A*_t_* is the *N* × *N* phylogenetic distance matrix implied by tree *t*, and L*_t_* is its lower-triangular Cholesky factor. The variable z*_t_* ∈ R*^N^* is a vector of standard normal auxiliaries, which generates u*_t_*, the phylogenetic random effects. The parameters αt, βt ~ N(0, 2) are the intercept and slope, respectively, for tree *t*. Finally, *ς_t_*, *σ_t_* ~ Half Student-*t* (0, 20) are the phylogenetic-signal standard deviation and the residual standard deviation, respectively.
2. **Posterior tree samples:** Rather than collapsing the tree distribution to a single consensus tree, the model is vectorised over the full tree tensor A ∈ R^*T*×*N*×*N*^ so that each tree yields an independent posterior, and the concatenated vector of posteriors samples of *α_t_*, *β_t_*, *ς_t_*, and *σ_t_* automatically reflect uncertainty caused by variation in topology and branch lengths. As for any *PGMM*, multiple covariates and interaction terms can be included [71].
3. **3. BI Implementation:**

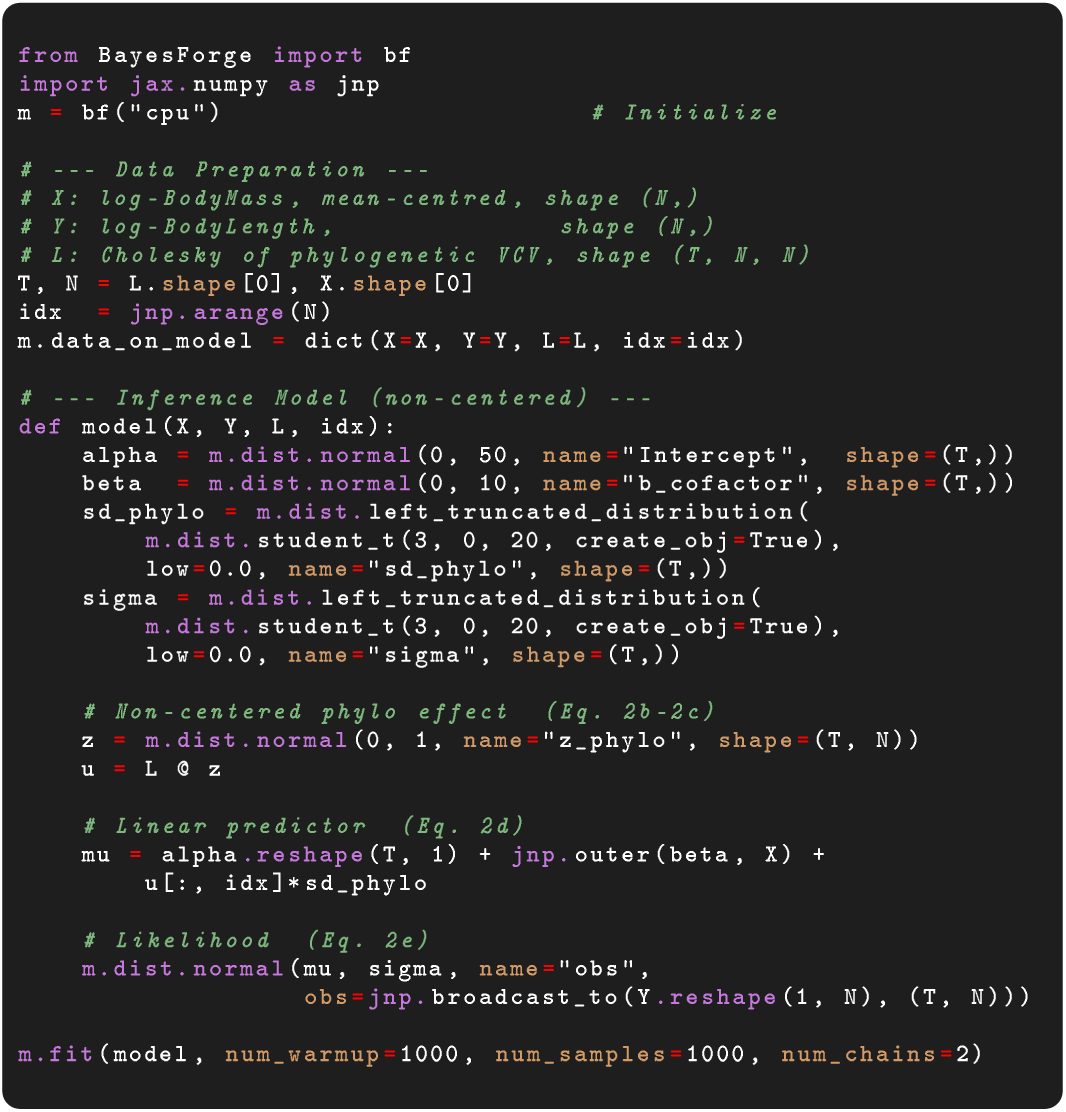
Full code and comparative estimation between BI and equivalent brms models can be found in Appendix 2. Appendix 2 also show that with matched priors and parameterisation, BI and brms recover the same posterior on all parameters to within 0.5% across 100 posterior trees.

## 5 Example 3 Vocal Repertoire Size Estimation: Custom undersampling model and latent-variable marginalization

Naturalistic observations are inherently prone to incomplete recording, which in turn requires specialized modeling approaches to account for non-detection rates while simultaneously estimating presence/absence and trait diversity. For example, because research on vocal complexity—such as variation in repertoire size—relies on finite recording sessions, it is highly susceptible to undersampling. This introduces a strong downward bias, underestimating the true vocal repertoire size by missing rare, low-probability syllables or sequences [72]. Consequently, finite observation windows not only miss latent vocal diversity, but can also violate the underlying model assumptions of equal observability across the repertoire. To model such data, a custom-made model tailored to account for heterogeneous production probabilities across vocalization types and unequal sampling effort across individuals needs to be developed [10]. While software like *Stan* requires complex, manual marginalization to bypass the inability to sample discrete parameters (e.g., whether an item exists in a repertoire), *BF* handles this natively through automated JAX-based enumeration on hardware-accelerated kernels. Following the framework of Sigmundson et al. [10], we show how this custom architecture enables the recovery of true latent diversity from noisy empirical data, providing a specialized modeling capability that is currently missing in the trove of ecological software available to researchers.

## 6 Scalability

We evaluate computational performance by comparing the execution time of the *Social Relations Model* (SRM) across networks of increasing size (50, 100, 200, and 400 nodes) using both *Stan* and *BF* (Figure 1a). For *BF*, computations were performed using both CPU and GPU implementations. CPU-based executions were carried out on a 120-core processor, while GPU-based executions were conducted on an NVIDIA A40 accelerator. Because the *SRM* in *BF* is implemented in a vectorized manner, and such an approach is not currently available in other software packages (e.g. STRAND [1], AMEN [73]), we implemented the same vectorized approach in *Stan* (*Appendix 4*). Additionally, before performing the benchmark, we compared the posterior distributions between the original non-vectorized model [1] and our new vectorized version in *Stan* and *BF*, and found them to be virtually identical (*Appendix 4*).

**FIGURE 1.**
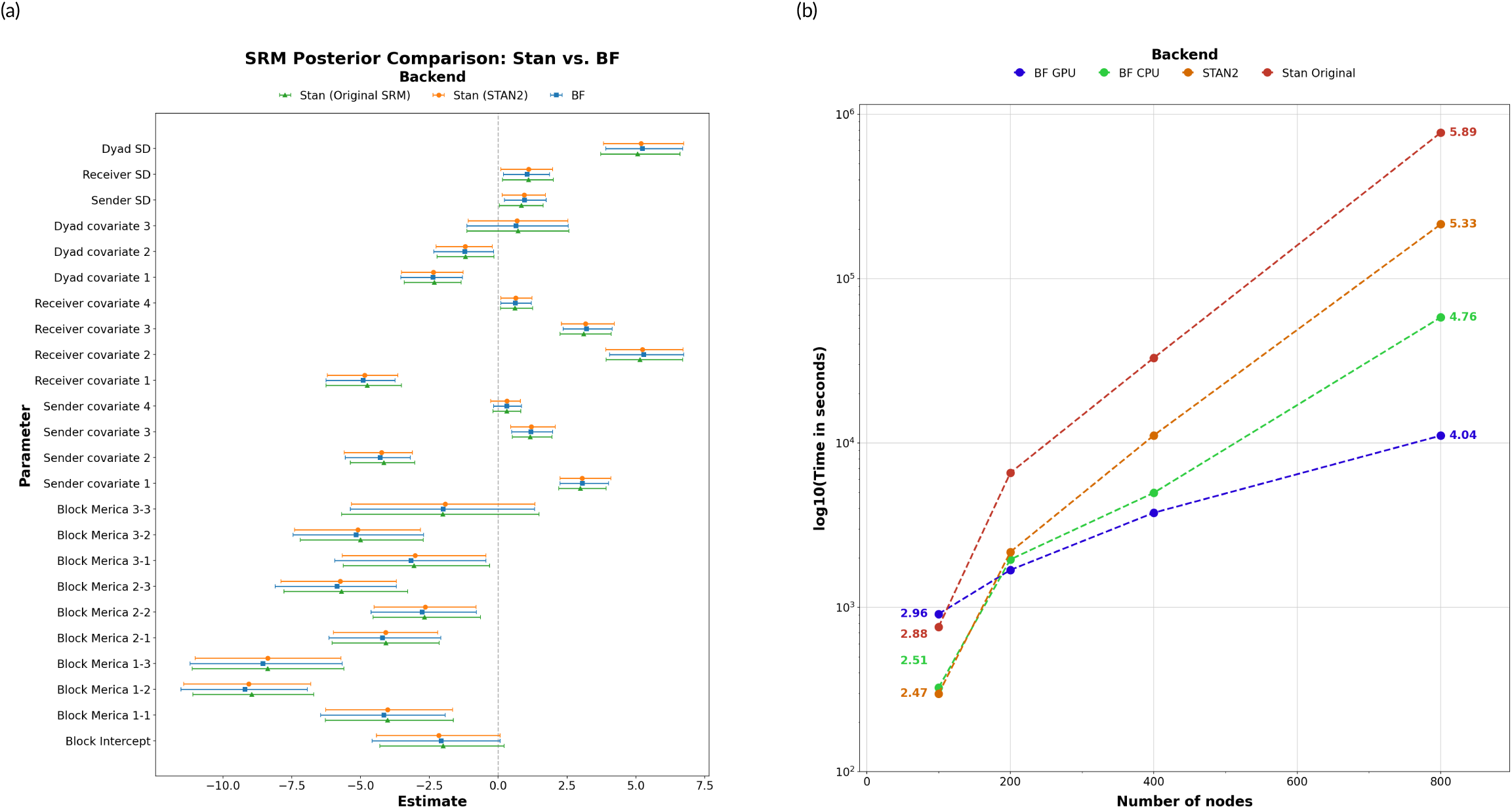
(A) Benchmarking runtime performance of Stan and BI (CPU and GPU implementations) for the Social Relations Model (SRM) across networks of varying sizes (50, 100, 200, 400 and 800 nodes). **(B)** Posterior distributions for model parameters.

### Box 3. Custom Latent-Variable Marginalization – Vocal Repertoire Size Estimation

1. **Measurement-error formulation:** Let *N* individuals be observed over *J* recording sessions of duration *d_j_* ∈ {1, …, *J* }. For a population-level lexicon of *M* unique vocalisation types, each individual *i* produces type *m* at latent rate *λ_m_* only when it possesses that unit (i.e., *Z_i,m_* = 1). In this example, we assume that possession is only ever partially observable. The observed count *Y_j,m_* of vocalisations of type *m* in session *j* is therefore a Zero-Inflated Poisson mixture. This model allows us to separate *structural zeros* (i.e., vocalisation types absent from repertoire) from *sampling zeros* (i.e., vocalisation types present in the repertoire, but not produced). We first aggregate the recording-specific data here by writing:

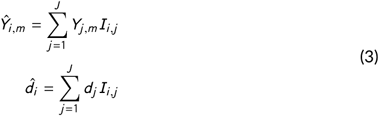

where *I_i,j_* equals 1 if recording *j* was produced by individual *i*, and is zero otherwise. Then, the model is as follows:

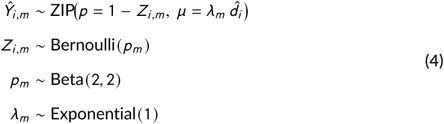 After fitting, the repertoire size for individual *i*, *R_i,m_*, is recovered via:

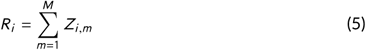
2. **Latent-variable marginalization:** Standard pipelines in Stan instead require the user to derive and code the probability of each token for each individual manually, and then use Bayes’ rule to determine the posterior repertoire size (see *Appendix 3*), while *BF* performs such computations automatically, provided the model’s dependence structure is declared via `plate`.
3. BI Implementation:

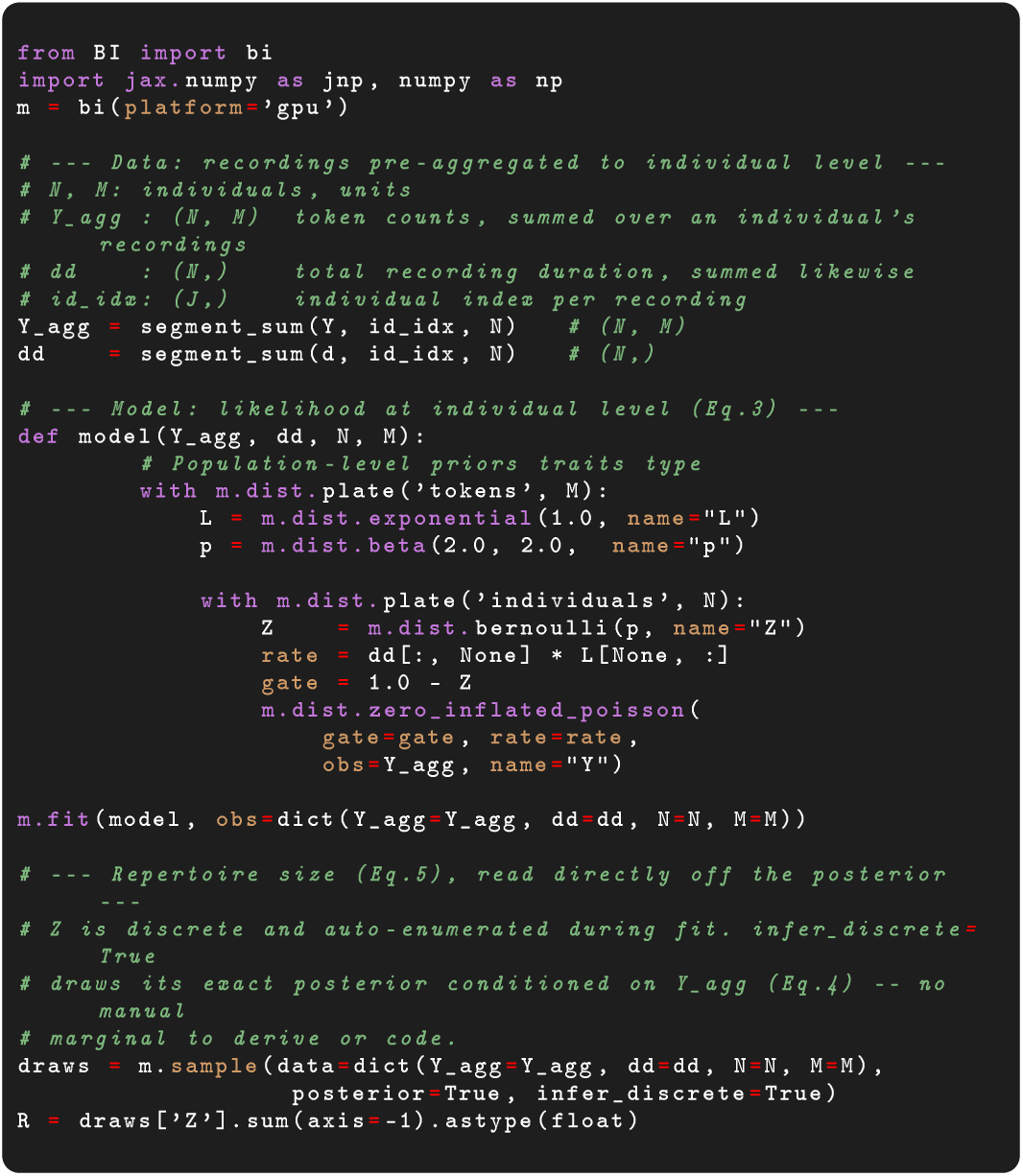
Full code, Stan benchmark, and recovery tests on synthetic data following Sigmundson et al. [10] are provided in *Appendix 3*.

Overall, *BF* exhibits substantially faster computation times than *Stan*, with the performance gap widening markedly as network size increases. In particular, for networks with 400 nodes, *BF* achieves up to a 19.32-fold reduction in execution time relative to *Stan* without substantial differences in *Effective Sample Size*. An additional noteworthy observation concerns GPU-based computation in *BF*: although GPU execution does not outperform CPU execution for smaller networks (e.g., 50 nodes), its rate of increase in computation time is considerably lower as network size grows, ultimately yielding superior performance for larger networks. Collectively, these results highlight the strong scalability of *BF* for complex models and large datasets, on both CPU and GPU architectures.

Parameter estimates obtained using *Stan* and *BF* for the SRM are highly consistent and accurately recover parameter values used in the simulation (Figure 1b). This validation across varying network sizes ensures that the computational enhancements provided by *BF* do not come at the cost of inferential accuracy. The observed posterior distributions overlap significantly with the true parameter values (represented by the dashed lines in Figure 1b), confirming that both the *JAX*-based sampler and the *Stan* benchmark converge on the same estimated quantities. Code to reproduce these simulations, the benchmark tests, and the posterior comparisons are available in *Appendix 4*.

## 7 Discussion

As the field of ecology embraces a growing number of large, mulit-layered datasets, traditional Markov chain Monte Carlo samplers, such as *Stan*, struggle to scale. Fitting a hierarchical network model using traditional CPU-bound software can require days of computation, effectively rendering a full Bayesian workflow impracticle. *BayesForge* alleviates this bottleneck by delegating mathematical operations to computational engines (*JAX*, *NumPyro* and *TensorFlow Probability*) designed for vectorization and parallelisation. By bringing *JAX*-based Just-In-Time (JIT) compilation and native GPU/TPU acceleration to the R and Julia ecosystems, *BF* facilitates computations that were previously out of reach.

The three applications presented in this study highlight the practical consequences of removing these computational and structural barriers.

First, the Social Relations Model (SRM) benchmark test demonstrates that GPU-accelerated tensor operations, through their increased efficiency, allow for the analysis of larger networks. This in turn, can fundamentally expand the scope of feasible data analysis problems, allowing researchers to apply complexe generative models to study systems characterized by large population sizes. Second, the phylogenetic application illustrates that eliminating the computational “uncertainty tax” can facilite the application of more appropirate models: ecologists no longer have to collapse speculative evolutionary histories into a single consensus tree, but can seamlessly vectorize their models across an entire distribution of potential trees. Finally, the vocal repertoire application demonstrates that automated, hardware-accelerated discrete enumeration facilitates coding complex latent-variable models. By removing the need for manual mathematical derivations (e.g., custom `log_sum_exp` loops), *BF* significantly lowers the barriers to entry for implementing architectures that handle structural unobserved states, such as Hidden Markov Models (HMMs) and occupancy models.

To properly contextualize *BayesForge*, it is necessary to map its position within the broader software ecosystem. *BF* is not intended to replace existing, highly optimized tools within their specific domains. *Stan* [14] remains the gold standard for highly expressive modeling where the user requires ultimate low-level algorithmic control. Regarding interfaces, *brms* [20] provides an unmatched, highly idiomatic formula interface for standard multilevel modeling natively within *R*. Concurrently, *PyMC* [15] serves as the premier environment for strictly Python-native users. *BayesForge* complements these tools by occupying a unique, synergistic niche. It acts as an interdisciplinary bridge, providing a curated library of over 50 pre-built models relevent for ecological applications (e.g., NBDA, SRM, and spatial Gaussian Processes), while integrating the efficiency of Python-native tensor mathematics (vectorisation, parallelisation and automatic sharding). This allows users in the *R* and *Julia* communities to build custom models without needing to use cross-language wrappers that were not design to facilitate statistical model building—e.g., *reticulate* can wrap *Python* code into *R*, but users must essentially hand-build *Python* versions of their models.

In designing this cross-platform framework, we purposefully navigated the steep trade-off between language-specific readability and cross-environment portability. Adhering simultaneously to the idiomatic conventions of *R*, *Python*, and *Julia* is impossible. Consequently, *BF* enforces a unified, slightly more explicit syntax across all three environments, requiring R users to forego some familiar formula interfaces in favor of explicitly defined probability distributions. Furthermore, we acknowledge the inherent technical friction of bridging software. While *BF* successfully abstracts away the notoriously difficult manual type-conversions of *JAX* device arrays that plague standard *reticulate* workflows, *R* and *Julia* users inherently rely on under-the-hood Python environments, which can occasionally introduce local installation friction. Finally, while *BF* drastically reduces computation time, it still require users to understand their data analysis problems, define appropriate models, and conduct suficient posterior predictive checks and model diagnostics.

Ultimately, *BayesForge* transforms best-practice Bayesian validation from a computational luxury into an accessible, everyday standard for applied ecologists. Historically, computational and software constraints have functioned as invisible methodological barriers, inadvertently forcing researchers to narrow their hypotheses, drop validation steps, or simplify their models to accommodate the limitations of their MCMC samplers. By unifying generative simulation, calibration, and hardware-accelerated inference into a streamlined API, *BF* ensures that rigorous, end-to-end model validation is as computationally accessible in *R* and *Julia* as it is in modern deep-learning platforms.

The computational ecology of the coming decade will be defined by the breadth of researchers who can feasibly build and validate mechanistic models at scale. As monitoring technologies produce increasingly massive and complex datasets, the overarching goal of statistical ecology must be to align our analytical complexity with the true complexity of natural systems. Moving forward, the development of *BayesForge* will focus on expanding its library of pre-built architectures through open-source community contributions, particularly targeting emerging, data-dense fields like movement ecology and spatial transcriptomics.

## 8 Data and Code Availability

The exact version of the code, appendices, and simulation scripts supporting this manuscript is permanently archived on [Zenodo](https://zenodo.org/records/21936604). The active codebase and development versions are maintained on [GitHub](https://github.com/BGN-for-ASNA/BF_MS). Stable releases of *BayesForge* can be installed directly from the official package repositories: [pip](https://pypi.org/project/BayesForge/) for Python, [CRAN](https://cran.r-project.org/web/packages/BayesForge/) for R, and the General Registry for [Julia](https://github.com/JuliaRegistries/General/tree/master/P/PyBayesForge). Active development versions and source code are hosted on their respective GitHub repositories (https://github.com/BGN-for-ASNA). Comprehensive documentation, including installation instructions, tutorials, and extended examples, are available on the official project [website](https://s-sosa.com/BF).

