## Appendix 1, 2, 3, and 4 for "BayesForge: A Bayesian Inference library for Python, R, and Julia": benchmark_gpu_only.py

```
import os, sys

# Must clear XLA_FLAGS and set PREALLOCATE=false BEFORE any JAX import
os.environ.pop("XLA_FLAGS", None)
os.environ["XLA_PYTHON_CLIENT_PREALLOCATE"] = "false"

_ld_paths = [
    "/home/sebastian_sosa/BI/nvidia_driver/extracted",
    "/home/sebastian_sosa/.cmdstan/cmdstan-2.38.0/stan/lib/stan_math/lib/tbb/",
]
_existing = os.environ.get("LD_LIBRARY_PATH", "")
os.environ["LD_LIBRARY_PATH"] = ":".join(_ld_paths) + (":" + _existing if _existing else "")

import jax
import jax.numpy as jnp
print("JAX devices:", jax.devices())

import numpy as np
import time
import pandas as pd
from BI import bi
from numpyro.diagnostics import summary as numpyro_summary


def simulate_data(N_nodes):
    N_focal_vars, N_target_vars, N_dyad_vars = 4, 4, 3

    m_utils = bi("gpu", rand_seed=False, print_devices_found=False)

    wide_focal_np  = np.random.normal(0, 1, size=(N_nodes, N_focal_vars))
    wide_target_np = np.random.normal(0, 1, size=(N_nodes, N_target_vars))

    dyadic_predictors_mat = np.random.binomial(1, 0.3, size=(N_nodes, N_nodes, N_dyad_vars))
    for k in range(N_dyad_vars):
        np.fill_diagonal(dyadic_predictors_mat[:, :, k], 0)
    dyadic_predictors_mat[:, :, 2] = (
        dyadic_predictors_mat[:, :, 0] * dyadic_predictors_mat[:, :, 1]
    )

    dyadic_predictors_edgl = jnp.stack(
        [m_utils.net.mat_to_edgl(dyadic_predictors_mat[:, :, k]) for k in range(N_dyad_vars)],
        axis=2,
    )

    Any_np    = np.zeros(N_nodes, dtype=int)
    Merica_np = np.random.choice([0, 1, 2], size=(N_nodes,), p=[0.25, 0.5625, 0.1875])

    N_by_grp_Any    = np.array([N_nodes])
    N_by_grp_Merica = np.array([np.sum(Merica_np == i) for i in range(3)])

    m_sim = bi("gpu", rand_seed=False, print_devices_found=False)
    B_intercept = m_sim.net.block_model(Any_np,    1, jnp.array(N_by_grp_Any),    sample=True, name="intercept")
    B_category  = m_sim.net.block_model(Merica_np, 3, jnp.array(N_by_grp_Merica), sample=True, name="category")
    sr = m_sim.net.sender_receiver(
        jnp.array(wide_focal_np), jnp.array(wide_target_np),
        s_mu=0.4, r_mu=-0.4, sample=True,
    )
    dr = m_sim.net.dyadic_effect(dyadic_predictors_edgl, d_sd=2.5, sample=True)
    network_edgl = m_sim.dist.bernoulli(logits=B_intercept + B_category + sr + dr, sample=True)

    return {
        "N_nodes":               N_nodes,
        "N_dyads":               network_edgl.shape[0],
        "network_edgl":          np.array(network_edgl),
        "dyadic_predictors_edgl": np.array(dyadic_predictors_edgl),
        "wide_focal_np":         np.array(wide_focal_np),
        "wide_target_np":        np.array(wide_target_np),
        "Any_np":                Any_np,
        "Merica_np":             Merica_np,
    }


def model_srm_wide(network_edgl, dyadic_predictors, sender_predictors,
                   receiver_predictors, Any, Merica, N_by_grp_Any, N_by_grp_Merica):
    try:
        from model_effects import Neteffect
    except ImportError:
        from BI.Network.model_effects import Neteffect

    m_inner  = bi("gpu", print_devices_found=False)
    B_any    = Neteffect.block_model(Any,    1, jnp.array(N_by_grp_Any),    name="intercept")
    B_Merica = Neteffect.block_model(Merica, 3, jnp.array(N_by_grp_Merica), name="Merica")
    sr = m_inner.net.sender_receiver(sender_predictors, receiver_predictors)
    dr = m_inner.net.dyadic_effect(dyadic_predictors)
    m_inner.dist.bernoulli(
        logits=B_any + B_Merica + sr + dr, obs=network_edgl, name="network_edgl"
    )


if __name__ == "__main__":
    script_dir  = os.path.dirname(os.path.abspath(__file__))
    csv_path    = os.path.join(script_dir, "benchmark_results_gpu_n800.csv")
    ITER_WARMUP = ITER_SAMPLING = 1000
    network_sizes = [800]
    results = []

    print("-" * 100)
    print(f"{'Model':<16} | {'N':<5} | {'Dyads':<6} | {'Time (s)':<10} | {'Min ESS':<10} | {'Max ESS':<10}")
    print("-" * 100)

    for N in network_sizes:
        np.random.seed(42)
        sim_data = simulate_data(N)

        N_by_grp_Any    = np.array([N])
        N_by_grp_Merica = np.array([np.sum(sim_data["Merica_np"] == i) for i in range(3)])

        m = bi("gpu", print_devices_found=False)
        m.data_on_model = dict(
            network_edgl          = sim_data["network_edgl"],
            dyadic_predictors     = sim_data["dyadic_predictors_edgl"],
            sender_predictors     = jnp.array(sim_data["wide_focal_np"]),
            receiver_predictors   = jnp.array(sim_data["wide_target_np"]),
            Any                   = jnp.array(sim_data["Any_np"],    dtype=jnp.int32),
            Merica                = jnp.array(sim_data["Merica_np"], dtype=jnp.int32),
            N_by_grp_Any          = N_by_grp_Any,
            N_by_grp_Merica       = N_by_grp_Merica,
        )

        t0 = time.time()
        m.fit(model_srm_wide, num_samples=ITER_SAMPLING, num_warmup=ITER_WARMUP, num_chains=1)
        elapsed = time.time() - t0

        try:
            posteriors_wc = {k: np.expand_dims(v, 0) for k, v in m.posteriors.items()}
            bi_sum = numpyro_summary(posteriors_wc)
            all_n_effs = np.array([
                val
                for d in bi_sum.values()
                for val in np.array(d["n_eff"]).flatten()
            ])
            all_n_effs = all_n_effs[np.isfinite(all_n_effs)]
            min_ess, max_ess = (
                (np.min(all_n_effs), np.max(all_n_effs)) if len(all_n_effs) else (0.0, 0.0)
            )
        except Exception:
            min_ess, max_ess = 0.0, 0.0

        results.append({
            "Model": "BI Wide (GPU)", "N_nodes": N, "N_dyads": sim_data["N_dyads"],
            "Time_s": elapsed, "Min_ESS": min_ess, "Max_ESS": max_ess,
            "Iter_Warmup": ITER_WARMUP, "Iter_Sampling": ITER_SAMPLING,
        })
        print(
            f"{'BI Wide (GPU)':<16} | {N:<5} | {sim_data['N_dyads']:<6} | "
            f"{elapsed:<10.2f} | {min_ess:<10.2f} | {max_ess:<10.2f}"
        )
        sys.stdout.flush()
        pd.DataFrame(results).to_csv(csv_path, index=False)

    print("-" * 100)
    print(f"GPU benchmark complete. Results saved to {csv_path}")
```
