## Appendix 1, 2, 3, and 4 for "BayesForge: A Bayesian Inference library for Python, R, and Julia": benchmark_stan2.py

```
import os
import sys

# Set LD_LIBRARY_PATH programmatically so JAX and CmdStan find their libraries automatically
ld_paths = [
    "/home/sebastian_sosa/BI/nvidia_driver/extracted",
    "/home/sebastian_sosa/.cmdstan/cmdstan-2.38.0/stan/lib/stan_math/lib/tbb/"
]
existing_ld = os.environ.get('LD_LIBRARY_PATH', '')
if existing_ld:
    os.environ['LD_LIBRARY_PATH'] = existing_ld + ":" + ":".join(ld_paths)
else:
    os.environ['LD_LIBRARY_PATH'] = ":".join(ld_paths)

import numpy as np
import time
import pandas as pd
from cmdstanpy import CmdStanModel
import multiprocessing as mp

# Setup device and BI model
from BI import bi
import jax.numpy as jnp
import jax
from numpyro.diagnostics import summary as numpyro_summary

# Prevent JAX from hogging memory completely so Stan can run
os.environ['XLA_PYTHON_CLIENT_PREALLOCATE'] = 'false'

def simulate_data(N_nodes):
    N_focal_vars = 4
    N_target_vars = 4
    N_dyad_vars = 3
    
    # Instantiate bi for network utilities
    m_utils = bi('cpu', rand_seed=False)
    
    # Generate individual (focal/target) predictors
    wide_focal_np = np.random.normal(0, 1, size=(N_nodes, N_focal_vars))
    wide_target_np = np.random.normal(0, 1, size=(N_nodes, N_target_vars))
    
    # Generate dyadic predictors (N x N matrices)
    dyadic_predictors_mat = np.random.binomial(1, 0.3, size=(N_nodes, N_nodes, N_dyad_vars))
    for k in range(N_dyad_vars):
        np.fill_diagonal(dyadic_predictors_mat[:, :, k], 0)
    # Make the 3rd dyadic predictor the interaction of the first two
    dyadic_predictors_mat[:, :, 2] = dyadic_predictors_mat[:, :, 0] * dyadic_predictors_mat[:, :, 1]
    
    # Convert dyadic predictors to edgelist (N_dyads, 2, N_dyad_vars)
    dyadic_predictors_edgl = jnp.stack([m_utils.net.mat_to_edgl(dyadic_predictors_mat[:, :, k]) for k in range(N_dyad_vars)], axis=2)
    
    # Generate block groups
    Any_np = np.zeros(N_nodes, dtype=int)
    Merica_np = np.random.choice([0, 1, 2], size=(N_nodes,), p=[0.25, 0.5625, 0.1875])
    
    N_grp_Any = 1
    N_grp_Merica = 3
    N_by_grp_Any = np.array([N_nodes])
    N_by_grp_Merica = np.array([np.sum(Merica_np == i) for i in range(N_grp_Merica)])
    
    # Simulate network outcomes using bi's generative APIs
    m_sim = bi('cpu', rand_seed=False)
    B_intercept = m_sim.net.block_model(Any_np, 1, jnp.array(N_by_grp_Any), sample=True, name="intercept")
    B_category = m_sim.net.block_model(Merica_np, 3, jnp.array(N_by_grp_Merica), sample=True, name="category")
    sr = m_sim.net.sender_receiver(
        jnp.array(wide_focal_np), 
        jnp.array(wide_target_np), 
        s_mu=0.4, r_mu=-0.4, sample=True)
    dr = m_sim.net.dyadic_effect(dyadic_predictors_edgl, d_sd=2.5, sample=True)
    logits = B_intercept + B_category + sr + dr
    network_edgl = m_sim.dist.bernoulli(logits=logits, sample=True)
    
    N_dyads = network_edgl.shape[0]
    N_obs = N_dyads * 2
    
    return {
        'N_nodes': N_nodes,
        'N_dyads': N_dyads,
        'N_obs': N_obs,
        'network_edgl': np.array(network_edgl),
        'dyadic_predictors_edgl': np.array(dyadic_predictors_edgl),
        'dyadic_predictors_mat': np.array(dyadic_predictors_mat),
        'wide_focal_np': np.array(wide_focal_np),
        'wide_target_np': np.array(wide_target_np),
        'Any_np': np.array(Any_np),
        'Merica_np': np.array(Merica_np),
        'N_dyad_vars': N_dyad_vars
    }

def get_stan_priors():
    priors = np.zeros((23, 2))
    p_data = [
        [-3.00, 1.5], [3.00, 1.5], [-1.50, 1.0], [1.00, 0.0], [1.00, 0.0], [1.00, 0.0],
        [0.00, 2.5], [0.00, 2.5], [0.00, 2.5], [0.10, 2.5], [0.01, 2.5], [0.00, 2.5],
        [0.00, 2.5], [0.00, 2.5], [0.00, 2.5], [0.00, 2.5], [2.50, 0.0], [2.50, 0.0],
        [1.50, 0.0], [3.00, 1.0], [2.00, 0.0], [3.00, 12.0], [0.00, 2.5]
    ]
    for i, row in enumerate(p_data):
        priors[i] = row
    return priors

def prepare_stan2_data(sim_data):
    N_nodes = sim_data['N_nodes']
    N_dyads = sim_data['N_dyads']
    N_obs = sim_data['N_obs']
    network_edgl = sim_data['network_edgl']
    dyadic_predictors_edgl = sim_data['dyadic_predictors_edgl']
    wide_focal_np = sim_data['wide_focal_np']
    wide_target_np = sim_data['wide_target_np']
    Any_np = sim_data['Any_np']
    Merica_np = sim_data['Merica_np']
    
    urows, ucols = np.triu_indices(N_nodes, k=1)
    long_ids_int = np.stack([urows, ucols], axis=1)
    
    sender = np.concatenate([long_ids_int[:, 0], long_ids_int[:, 1]]) + 1
    receiver = np.concatenate([long_ids_int[:, 1], long_ids_int[:, 0]]) + 1
    dyad_id = np.concatenate([np.arange(1, N_dyads + 1), np.arange(1, N_dyads + 1)])
    dyad_dir = np.concatenate([np.ones(N_dyads, dtype=int), np.full(N_dyads, 2, dtype=int)])
    outcomes_srm2 = np.concatenate([network_edgl[:, 0], network_edgl[:, 1]])
    
    focal_set_stan = np.column_stack([np.ones(N_nodes), wide_focal_np])
    target_set_stan = np.column_stack([np.ones(N_nodes), wide_target_np])
    
    flat_dyad_preds = np.concatenate([dyadic_predictors_edgl[:, 0, :], dyadic_predictors_edgl[:, 1, :]], axis=0)
    dyad_set_stan = np.column_stack([np.ones(N_obs), flat_dyad_preds])
    
    block_set_stan = np.column_stack([Any_np + 1, Merica_np + 1])
    priors = get_stan_priors()
        
    stan_data = {
        'N_networktypes': 1,
        'N_id': N_nodes,
        'N_dyads': N_dyads,
        'N_obs': N_obs,
        'N_responses': 1,
        'N_params': [wide_focal_np.shape[1] + 1, wide_target_np.shape[1] + 1, flat_dyad_preds.shape[1] + 1],
        'sender': sender.tolist(),
        'receiver': receiver.tolist(),
        'dyad_id': dyad_id.tolist(),
        'dyad_dir': dyad_dir.tolist(),
        'outcomes': outcomes_srm2.astype(int).tolist(),
        'outcomes_real': outcomes_srm2.astype(float).tolist(),
        'exposure': np.ones(N_obs, dtype=int).tolist(),
        'N_group_vars': 2,
        'max_N_groups': 3,
        'N_groups_per_var': [1, 3],
        'block_set': block_set_stan.tolist(),
        'focal_set': focal_set_stan.tolist(),
        'target_set': target_set_stan.tolist(),
        'dyad_set': dyad_set_stan.tolist(),
        'priors': priors.tolist(),
        'export_network': 0,
        'outcome_mode': 1,
        'link_mode': 1
    }
    return stan_data

def prepare_orig_stan_data(sim_data):
    N_nodes = sim_data['N_nodes']
    N_dyads = sim_data['N_dyads']
    N_obs = sim_data['N_obs']
    network_edgl = sim_data['network_edgl']
    dyadic_predictors_mat = sim_data['dyadic_predictors_mat']
    wide_focal_np = sim_data['wide_focal_np']
    wide_target_np = sim_data['wide_target_np']
    Any_np = sim_data['Any_np']
    Merica_np = sim_data['Merica_np']
    N_dyad_vars = sim_data['N_dyad_vars']
    
    urows, ucols = np.triu_indices(N_nodes, k=1)
    
    Y = np.zeros((N_nodes, N_nodes), dtype=int)
    for d in range(N_dyads):
        u = urows[d]
        v = ucols[d]
        Y[u, v] = int(network_edgl[d, 0])
        Y[v, u] = int(network_edgl[d, 1])

    outcomes_3d = Y.reshape(N_nodes, N_nodes, 1).tolist()
    outcomes_real_3d = Y.reshape(N_nodes, N_nodes, 1).astype(float).tolist()
    exposure_3d = np.ones((N_nodes, N_nodes, 1), dtype=int).tolist()
    mask_3d = np.zeros((N_nodes, N_nodes, 1), dtype=int).tolist()

    dyad_set_original = np.zeros((N_nodes, N_nodes, N_dyad_vars + 1))
    dyad_set_original[:, :, 0] = 1.0  
    dyad_set_original[:, :, 1:] = dyadic_predictors_mat
    
    focal_set_stan = np.column_stack([np.ones(N_nodes), wide_focal_np])
    target_set_stan = np.column_stack([np.ones(N_nodes), wide_target_np])
    block_set_stan = np.column_stack([Any_np + 1, Merica_np + 1])
    priors = get_stan_priors()

    original_stan_data = {
        'N_networktypes': 1,
        'N_id': N_nodes,
        'N_responses': 1,
        'N_params': [wide_focal_np.shape[1] + 1, wide_target_np.shape[1] + 1, N_dyad_vars + 1],
        'outcomes': outcomes_3d,
        'outcomes_real': outcomes_real_3d,
        'exposure': exposure_3d,
        'mask': mask_3d,
        'focal_set': focal_set_stan.tolist(),
        'target_set': target_set_stan.tolist(),
        'dyad_set': dyad_set_original.tolist(),
        'priors': priors.tolist(),
        'export_network': 0,
        'outcome_mode': 1,
        'link_mode': 1,
        'N_group_vars': 2,
        'max_N_groups': 3,
        'N_groups_per_var': [1, 3],
        'block_set': block_set_stan.tolist()
    }
    return original_stan_data

def get_stan_ess(fit):
    try:
        summ = fit.summary()
        col = None
        for c in ['ESS_bulk', 'N_Eff', 'ess_bulk']:
            if c in summ.columns:
                col = c
                break
        if col is None:
            for c in summ.columns:
                if 'eff' in c.lower() or 'ess' in c.lower():
                    col = c
                    break
        if col is None:
            return 0.0, 0.0
        n_effs = summ[col].dropna().values
        n_effs = n_effs[np.isfinite(n_effs)]
        if len(n_effs) == 0:
            return 0.0, 0.0
        return np.min(n_effs), np.max(n_effs)
    except Exception as e:
        print("Error calculating Stan ESS:", e)
        return 0.0, 0.0

def run_bi_isolated(device, sim_data, N, ITER_WARMUP, ITER_SAMPLING, q):
    """
    Runs BI inference in a strictly isolated process.
    This guarantees JAX initializes freshly with the correct backend device.
    """
    import time
    import jax
    import jax.numpy as jnp
    import numpy as np
    from BI import bi
    import os
    os.environ['XLA_PYTHON_CLIENT_PREALLOCATE'] = 'false'
    # When running CPU-only, hide the GPU so JAX cannot accidentally allocate VRAM
    if device == 'cpu':
        os.environ['CUDA_VISIBLE_DEVICES'] = ''
    
    def model_srm_wide(network_edgl, dyadic_predictors, sender_predictors, receiver_predictors, Any, Merica, N_by_grp_Any, N_by_grp_Merica):
        try:
            from model_effects import Neteffect
        except ImportError:
            from BI.Network.model_effects import Neteffect
            
        m2_inner = bi(device)
        B_any    = Neteffect.block_model(Any,    1,    jnp.array(N_by_grp_Any),    name='intercept')
        B_Merica = Neteffect.block_model(Merica, 3,    jnp.array(N_by_grp_Merica), name='Merica')
        sr = m2_inner.net.sender_receiver(sender_predictors, receiver_predictors)
        dr = m2_inner.net.dyadic_effect(dyadic_predictors)
        m2_inner.dist.bernoulli(logits=B_any + B_Merica + sr + dr, obs=network_edgl, name='network_edgl')

    m2 = bi(device)
    
    N_grp_Any = 1
    N_grp_Merica = 3
    N_by_grp_Any = np.array([N])
    N_by_grp_Merica = np.array([np.sum(sim_data['Merica_np'] == i) for i in range(N_grp_Merica)])
    
    m2.data_on_model = dict(
        network_edgl=sim_data['network_edgl'], 
        dyadic_predictors=sim_data['dyadic_predictors_edgl'], 
        sender_predictors=jnp.array(sim_data['wide_focal_np']), 
        receiver_predictors=jnp.array(sim_data['wide_target_np']), 
        Any=jnp.array(sim_data['Any_np'], dtype=jnp.int32), 
        Merica=jnp.array(sim_data['Merica_np'], dtype=jnp.int32),
        N_by_grp_Any=N_by_grp_Any,
        N_by_grp_Merica=N_by_grp_Merica
    )
    
    start_time = time.time()
    m2.fit(model_srm_wide, num_samples=ITER_SAMPLING, num_warmup=ITER_WARMUP, num_chains=1)
    end_time = time.time()
    elapsed_bi = end_time - start_time
    
    from numpyro.diagnostics import summary as numpyro_summary
    try:
        posteriors_with_chains = {k: np.expand_dims(v, axis=0) for k, v in m2.posteriors.items()}
        bi_sum = numpyro_summary(posteriors_with_chains)
        all_n_effs = []
        for k, v in bi_sum.items():
            if 'n_eff' in v:
                all_n_effs.extend(np.array(v['n_eff']).flatten().tolist())
        all_n_effs = np.array(all_n_effs)
        all_n_effs = all_n_effs[np.isfinite(all_n_effs)]
        if len(all_n_effs) == 0:
            min_ess, max_ess = 0.0, 0.0
        else:
            min_ess, max_ess = np.min(all_n_effs), np.max(all_n_effs)
    except Exception as e:
        min_ess, max_ess = 0.0, 0.0

    q.put((elapsed_bi, min_ess, max_ess))

def run_benchmark():
    import argparse
    parser = argparse.ArgumentParser(description="Run SRM Benchmarks")
    parser.add_argument('-n', '--nodes', nargs='+', type=int, default=[50], help="List of network sizes (N_nodes) to test")
    parser.add_argument('--skip-stan', action='store_true', help="Skip Stan models and only run BI Wide")
    
    group = parser.add_mutually_exclusive_group(required=True)
    group.add_argument('--cpu', action='store_true', help="Run BI Wide model on CPU (includes Stan models unless skipped)")
    group.add_argument('--gpu', action='store_true', help="Run BI Wide model on GPU (skips Stan models)")
    group.add_argument('--both', action='store_true', help="Run both CPU and GPU benchmarks sequentially")
    
    args = parser.parse_args()
    network_sizes = args.nodes

    script_dir = os.path.dirname(os.path.abspath(__file__))
    stan2_file = os.path.join(script_dir, 'STAN2.stan')
    orig_stan_file = os.path.join(script_dir, 'original_srm.stan')
    
    if args.both:
        csv_filename = 'benchmark_results_both.csv'
        devices_to_test = ['cpu', 'gpu']
        run_stan = not args.skip_stan
    elif args.gpu:
        csv_filename = 'benchmark_results_gpu.csv'
        devices_to_test = ['gpu']
        run_stan = False
    else:
        csv_filename = 'benchmark_results_cpu.csv'
        devices_to_test = ['cpu']
        run_stan = not args.skip_stan
        
    csv_path = os.path.join(script_dir, csv_filename)
    
    # Suppress cmdstanpy logging for clean output
    import logging
    logging.getLogger("cmdstanpy").setLevel(logging.WARNING)
    if run_stan:
        print("Compiling STAN2 model before benchmark...")
        sm_stan2 = CmdStanModel(stan_file=stan2_file)
        print("Compiling Stan Original model before benchmark...")
        sm_orig = CmdStanModel(stan_file=orig_stan_file)
        print("Compilation complete.\n")
    
    results = []
    
    print("-" * 110)
    print(f"{'Model':<15} | {'Size':<6} | {'Dyads':<6} | {'Time (s)':<10} | {'Min ESS':<10} | {'Max ESS':<10} | {'Iter Warmup':<12} | {'Iter Sampling':<13}")
    print("-" * 110)
    
    ITER_WARMUP = 1000
    ITER_SAMPLING = 1000
    
    for N in network_sizes:
        np.random.seed(42)
        sim_data = simulate_data(N)
        
        # --- 1. Run STAN2 ---
        if run_stan:
            stan2_data = prepare_stan2_data(sim_data)
            start_time = time.time()
            fit_stan2 = sm_stan2.sample(data=stan2_data, iter_sampling=ITER_SAMPLING, iter_warmup=ITER_WARMUP, chains=1, show_progress=False)
            end_time = time.time()
            elapsed_stan2 = end_time - start_time
            min_ess_stan2, max_ess_stan2 = get_stan_ess(fit_stan2)
            
            results.append({
                'Model': 'STAN2', 'N_nodes': N, 'N_dyads': sim_data['N_dyads'], 
                'Time_s': elapsed_stan2, 'Min_ESS': min_ess_stan2, 'Max_ESS': max_ess_stan2,
                'Iter_Warmup': ITER_WARMUP, 'Iter_Sampling': ITER_SAMPLING
            })
            print(f"{'STAN2':<15} | {N:<6} | {sim_data['N_dyads']:<6} | {elapsed_stan2:<10.2f} | {min_ess_stan2:<10.2f} | {max_ess_stan2:<10.2f} | {ITER_WARMUP:<12} | {ITER_SAMPLING:<13}")
            sys.stdout.flush()
            pd.DataFrame(results).to_csv(csv_path, index=False)
        
        # --- 2. Run Stan Original ---
        if run_stan:
            orig_data = prepare_orig_stan_data(sim_data)
            start_time = time.time()
            fit_orig = sm_orig.sample(data=orig_data, iter_sampling=ITER_SAMPLING, iter_warmup=ITER_WARMUP, chains=1, show_progress=False)
            end_time = time.time()
            elapsed_orig = end_time - start_time
            min_ess_orig, max_ess_orig = get_stan_ess(fit_orig)
            
            results.append({
                'Model': 'Stan Original', 'N_nodes': N, 'N_dyads': sim_data['N_dyads'], 
                'Time_s': elapsed_orig, 'Min_ESS': min_ess_orig, 'Max_ESS': max_ess_orig,
                'Iter_Warmup': ITER_WARMUP, 'Iter_Sampling': ITER_SAMPLING
            })
            print(f"{'Stan Original':<15} | {N:<6} | {sim_data['N_dyads']:<6} | {elapsed_orig:<10.2f} | {min_ess_orig:<10.2f} | {max_ess_orig:<10.2f} | {ITER_WARMUP:<12} | {ITER_SAMPLING:<13}")
            sys.stdout.flush()
            pd.DataFrame(results).to_csv(csv_path, index=False)

        # --- 3. Run BI Wide (CPU and potentially GPU via subprocess) ---
        for device in devices_to_test:
            bi_model_name = f"BI Wide ({device.upper()})"
            
            ctx = mp.get_context('spawn')
            q = ctx.Queue()
            p = ctx.Process(target=run_bi_isolated, args=(device, sim_data, N, ITER_WARMUP, ITER_SAMPLING, q))
            p.start()
            p.join()
            
            # If process crashed or failed, q might be empty
            if not q.empty():
                elapsed_bi, min_ess_bi, max_ess_bi = q.get()
            else:
                elapsed_bi, min_ess_bi, max_ess_bi = 0.0, 0.0, 0.0
                print(f"[ERROR] {bi_model_name} execution failed in subprocess.")
            
            results.append({
                'Model': bi_model_name, 'N_nodes': N, 'N_dyads': sim_data['N_dyads'], 
                'Time_s': elapsed_bi, 'Min_ESS': min_ess_bi, 'Max_ESS': max_ess_bi,
                'Iter_Warmup': ITER_WARMUP, 'Iter_Sampling': ITER_SAMPLING
            })
            print(f"{bi_model_name:<15} | {N:<6} | {sim_data['N_dyads']:<6} | {elapsed_bi:<10.2f} | {min_ess_bi:<10.2f} | {max_ess_bi:<10.2f} | {ITER_WARMUP:<12} | {ITER_SAMPLING:<13}")
            sys.stdout.flush()
            pd.DataFrame(results).to_csv(csv_path, index=False)

    print("-" * 110)
    print(f"Benchmark complete. Results saved to {csv_path}")
    
    # Generate plot
    try:
        plot_benchmark(csv_path)
    except Exception as e:
        print(f"Could not generate plot: {e}")

def plot_benchmark(csv_path):
    import pandas as pd
    import matplotlib.pyplot as plt
    import numpy as np
    import os

    df = pd.read_csv(csv_path)

    plt.figure(figsize=(8.27, 11.69))

    colors = {
        'STAN2': '#D55E00',
        'Stan Original': '#FFA500', 
        'BI Wide (GPU)': '#2500D6',
        'BI Wide (CPU)': '#02D600'
    }

    # Group by model
    for model in df['Model'].unique():
        subset = df[df['Model'] == model].sort_values('N_nodes')
        if subset.empty:
            continue
            
        x = subset['N_nodes'].values
        y = subset['Time_s'].values
        
        c = colors.get(model, 'black')
        plt.plot(x, y, label=model, color=c, linestyle='--', linewidth=2, marker='o', markersize=8)
        
        # Add text labels at first and last points
        first_x, first_y = x[0], y[0]
        last_x, last_y = x[-1], y[-1]
        
        # Add offsets to prevent overlap with markers
        offset_x = (x.max() - x.min()) * 0.03 if len(x) > 1 else 5
        
        # Log10 labels
        plt.text(first_x - offset_x, first_y, f"{np.log10(first_y):.2f}", color=c, ha='right', va='center', fontsize=12, 
                 bbox=dict(facecolor='white', edgecolor='none', alpha=0.7, pad=0))
        plt.text(last_x + offset_x, last_y, f"{np.log10(last_y):.2f}", color=c, ha='left', va='center', fontsize=12,
                 bbox=dict(facecolor='white', edgecolor='none', alpha=0.7, pad=0))

    plt.yscale('log')
    plt.xlabel('Number of nodes', fontsize=18, fontweight='bold')
    plt.ylabel('Time in seconds', fontsize=18, fontweight='bold')
    
    plt.xticks(fontsize=16)
    plt.yticks(fontsize=16)
    
    plt.legend(title='Backend', title_fontsize=16, fontsize=16, loc='upper center', bbox_to_anchor=(0.5, 1.1), ncol=2, frameon=False)
    
    # Minimal theme styling
    plt.grid(True, which='major', linestyle='-', alpha=0.2)
    plt.grid(True, which='minor', linestyle=':', alpha=0.2)
    plt.gca().spines['top'].set_visible(False)
    plt.gca().spines['right'].set_visible(False)
    
    plt.tight_layout()
    
    base_csv = os.path.basename(csv_path)
    if 'gpu' in base_csv:
        plot_name = 'Benchmark_plot_gpu.png'
    elif 'both' in base_csv:
        plot_name = 'Benchmark_plot_both.png'
    else:
        plot_name = 'Benchmark_plot_cpu.png'
    out_path = os.path.join(os.path.dirname(csv_path), plot_name)
    plt.savefig(out_path, dpi=300, bbox_inches='tight')
    print(f"Benchmark plot saved to {out_path}")

if __name__ == '__main__':
    run_benchmark()
```
