## Appendix 1, 2, 3, and 4 for "BayesForge: A Bayesian Inference library for Python, R, and Julia": plot_benchmark.py

```
import pandas as pd
import numpy as np
import matplotlib.pyplot as plt
import matplotlib.ticker as ticker
import os

cpu_df = pd.read_csv("benchmark_results_cpu.csv")
gpu_csv = "benchmark_results_gpu.csv"
if os.path.exists(gpu_csv):
    gpu_df = pd.read_csv(gpu_csv)
    df = pd.concat([cpu_df, gpu_df], ignore_index=True)
else:
    df = cpu_df

# Map models to display labels, colors, and first-point vertical offsets (log-space multipliers)
model_map = {
    "BI Wide (GPU)":  ("BF GPU",        "#2500D6", 1.00),   # blue,   label centered
    "BI Wide (CPU)":  ("BF CPU",        "#2ecc40", 1.45),   # green,  label up
    "STAN2":          ("STAN2",          "#d46a00", 1.00),   # orange, label center
    "Stan Original":  ("Stan Original",  "#c0392b", 0.90),   # red,    label down slightly
}

fig, ax = plt.subplots(figsize=(14, 15.2))

x_max = df["N_nodes"].max()
x_margin = x_max * 0.12  # right margin for end labels

for model_key, (label, color, y_mult) in model_map.items():
    sub = df[df["Model"] == model_key].sort_values("N_nodes")
    if sub.empty:
        continue
    x = sub["N_nodes"].values
    y = sub["Time_s"].values

    ax.plot(x, y, color=color, linewidth=2.5, linestyle="--",
            marker="o", markersize=11, markerfacecolor=color,
            markeredgecolor=color, label=label, zorder=3)

    # Label at first point (left side, staggered vertically)
    ax.annotate(
        f"{np.log10(y[0]):.2f}",
        xy=(x[0], y[0]),
        xytext=(x[0] - 15, y[0] * y_mult),
        va="center", ha="right",
        color=color, fontsize=16, fontweight="bold",
    )

    # Label at last point (right side)
    ax.annotate(
        f"{np.log10(y[-1]):.2f}",
        xy=(x[-1], y[-1]),
        xytext=(x[-1] + 12, y[-1]),
        va="center", ha="left",
        color=color, fontsize=16, fontweight="bold",
    )

ax.set_yscale("log")
ax.yaxis.set_major_formatter(ticker.FuncFormatter(
    lambda val, _: f"$10^{{{int(np.log10(val))}}}$" if val > 0 else ""
))
ax.set_yticks([10**i for i in range(2, 7)])

ax.set_xlabel("Number of nodes", fontsize=20, fontweight="bold", labelpad=10)
ax.set_ylabel("log10(Time in seconds)", fontsize=20, fontweight="bold", labelpad=10)

ax.set_xlim(left=-10, right=x_max + x_margin)
ax.xaxis.set_major_locator(ticker.MultipleLocator(200))
ax.xaxis.set_minor_locator(ticker.MultipleLocator(100))

ax.grid(True, which="major", color="#cccccc", linewidth=0.8, zorder=0)
ax.set_facecolor("white")
fig.patch.set_facecolor("white")
ax.tick_params(axis="both", labelsize=16)

# Legend at top center, outside axes
legend = ax.legend(
    title="Backend", title_fontsize=20,
    loc="upper center", bbox_to_anchor=(0.5, 1.10),
    ncol=4, frameon=False, fontsize=16,
    handlelength=1.0, handleheight=1.0,
    markerscale=1.0,
)
legend.get_title().set_fontweight("bold")

plt.tight_layout()
plt.savefig("benchmark_plot_cpu.png", dpi=150)
print("Saved benchmark_plot_cpu.png")
```
