## Appendix 1, 2, 3, and 4 for "BayesForge: A Bayesian Inference library for Python, R, and Julia": plot_efficiency.py

```
import pandas as pd
import numpy as np
import matplotlib.pyplot as plt
import matplotlib.ticker as ticker
import os

cpu_df = pd.read_csv("benchmark_results_cpu.csv")
gpu_csv = "benchmark_results_gpu.csv"
if os.path.exists(gpu_csv):
    gpu_df = pd.read_csv(gpu_csv)
    df = pd.concat([cpu_df, gpu_df], ignore_index=True)
else:
    df = cpu_df

df["Min_ESS_per_s"] = df["Min_ESS"] / df["Time_s"]
df["Max_ESS_per_s"] = df["Max_ESS"] / df["Time_s"]

model_map = {
    "BI Wide (GPU)": ("BI GPU",        "#2500D6"),
    "BI Wide (CPU)": ("BI CPU",        "#2ecc40"),
    "STAN2":         ("STAN2",         "#d46a00"),
    "Stan Original": ("Stan Original", "#c0392b"),
}

# vertical offsets for first-point labels (log-space multipliers)
first_offsets = {
    "BI Wide (GPU)": 3.5,
    "BI Wide (CPU)": 2.0,
    "STAN2":         1.0,
    "Stan Original": 0.45,
}

def fmt(v):
    """Format ESS/s value: use scientific for very small, 3 sig figs otherwise."""
    if v < 0.001:
        return f"{v:.2e}"
    if v < 0.1:
        return f"{v:.4f}"
    return f"{v:.3f}"

fig, axes = plt.subplots(1, 2, figsize=(16, 7), sharey=False)
panel_info = [
    ("Min_ESS_per_s", "Min ESS per second"),
    ("Max_ESS_per_s", "Max ESS per second"),
]

for ax, (col, title) in zip(axes, panel_info):
    x_max = df["N_nodes"].max()
    x_margin = x_max * 0.14

    for model_key, (label, color) in model_map.items():
        sub = df[df["Model"] == model_key].sort_values("N_nodes")
        if sub.empty:
            continue
        x = sub["N_nodes"].values
        y = sub[col].values

        ax.plot(x, y, color=color, linewidth=2.5, linestyle="--",
                marker="o", markersize=11, markerfacecolor=color,
                markeredgecolor=color, label=label, zorder=3)

        y_mult = first_offsets[model_key]
        # first-point label (left side)
        ax.annotate(
            fmt(y[0]),
            xy=(x[0], y[0]),
            xytext=(x[0] - 15, y[0] * y_mult),
            va="center", ha="right",
            color=color, fontsize=11, fontweight="bold",
        )
        # last-point label (right side)
        ax.annotate(
            fmt(y[-1]),
            xy=(x[-1], y[-1]),
            xytext=(x[-1] + 12, y[-1]),
            va="center", ha="left",
            color=color, fontsize=11, fontweight="bold",
        )

    ax.set_yscale("log")
    ax.set_title(title, fontsize=15, fontweight="bold", pad=12)
    ax.set_xlabel("Number of nodes", fontsize=14, fontweight="bold", labelpad=8)
    ax.set_ylabel("ESS / second  (higher = faster)", fontsize=13, fontweight="bold", labelpad=8)
    ax.set_xlim(left=-10, right=x_max + x_margin)
    ax.xaxis.set_major_locator(ticker.MultipleLocator(200))
    ax.grid(True, which="major", color="#cccccc", linewidth=0.8, zorder=0)
    ax.set_facecolor("white")
    ax.tick_params(axis="both", labelsize=11)

    legend = ax.legend(
        title="Backend", title_fontsize=11,
        loc="upper right", frameon=False, fontsize=10,
    )
    legend.get_title().set_fontweight("bold")

fig.patch.set_facecolor("white")
plt.tight_layout()
plt.savefig("benchmark_efficiency.png", dpi=150, bbox_inches="tight")
print("Saved benchmark_efficiency.png")
```
