## Appendix 1, 2, 3, and 4 for "BayesForge: A Bayesian Inference library for Python, R, and Julia": README.md

# Benchmark Suite

Benchmarks comparing BI (CPU/GPU), STAN2, and Stan Original on the SRM network model
across network sizes N = 100, 200, 400, 800.

## Reproducing the plots

Requires the `myenv` virtualenv (has `pandas`/`matplotlib`; the system `python3` does not).

```bash
cd /home/sebastian_sosa/BI/Test/Network/SRM/benchmark_suite
/home/sebastian_sosa/myenv/bin/python3 plot_efficiency.py   # -> benchmark_efficiency.png (Min/Max ESS per second)
/home/sebastian_sosa/myenv/bin/python3 plot_benchmark.py    # -> benchmark_plot_cpu.png (wall-clock time)
```

Both scripts read two CSVs from the current directory and concatenate them:

- `benchmark_results_cpu.csv` — STAN2, Stan Original, BI Wide (CPU)
- `benchmark_results_gpu.csv` — BI Wide (GPU)

If `benchmark_results_gpu.csv` is missing, the plots fall back to CPU-only backends.

## Regenerating the underlying results

- CPU backends (STAN2 / Stan Original / BI CPU): `benchmark_stan2.py`
- GPU backend (BI Wide, JAX): `benchmark_gpu_only.py`
  ```bash
  cd /home/sebastian_sosa/BI/Test/Network/SRM/benchmark_suite
  CUDA_VISIBLE_DEVICES=<gpu_id> XLA_PYTHON_CLIENT_ALLOCATOR=platform \
    /home/sebastian_sosa/myenv/bin/python3 benchmark_gpu_only.py
  ```
  `XLA_PYTHON_CLIENT_ALLOCATOR=platform` avoids an XLA constant-allocation failure seen at N=800.
  Edit `network_sizes` and `csv_path` at the bottom of the script to control which sizes are run
  and where results are written, then merge into `benchmark_results_gpu.csv` before plotting.

Each script appends/writes rows with columns:
`Model, N_nodes, N_dyads, Time_s, Min_ESS, Max_ESS, Iter_Warmup, Iter_Sampling`.
