## Appendix 1, 2, 3, and 4 for "BayesForge: A Bayesian Inference library for Python, R, and Julia": STAN2.stan

```
data {
    int N_networktypes;                                               
    int N_id;           // Number of individuals                                                                                                 
    int N_dyads;        // Number of unique dyads
    int N_obs;          // Number of observations (edges/rows in the edgelist)
    int N_responses;        

    array[3] int N_params;                                          
    
    // --- Long format (edgelist) inputs ---
    array[N_obs] int sender;      // Index of sender (1 to N_id)
    array[N_obs] int receiver;    // Index of receiver (1 to N_id)
    array[N_obs] int dyad_id;     // Index of dyad (1 to N_dyads)
    array[N_obs] int dyad_dir;    // Direction identifier (1 or 2)

    array[N_obs] int outcomes;  
    vector[N_obs] outcomes_real; 
    array[N_obs] int exposure; 

    // --- Block structure ---
    int N_group_vars;                          
    int max_N_groups;                          
    array[N_group_vars] int N_groups_per_var;  
    array[N_id, N_group_vars] int block_set;   

    // --- Predictors ---
    matrix[N_id, N_params[1]] focal_set;
    matrix[N_id, N_params[2]] target_set;
    matrix[N_obs, N_params[3]] dyad_set; 

    matrix[23, 2] priors;
    
    int export_network;
    int outcome_mode;
    int link_mode;                       
}

transformed data {
    matrix[N_id, N_params[1]-1] focal_individual_predictors; 
    matrix[N_id, N_params[2]-1] target_individual_predictors; 
    matrix[N_obs, N_params[3]-1] flat_dyad_preds; 

    array[max_N_groups, N_group_vars] int N_per_group;      
    array[N_group_vars+1] int block_indexes;                
    int block_param_size;                                   

    block_param_size = 0;                             
    block_indexes[1] = 0;                             
    
    for(q in 1: N_group_vars){
      block_param_size += N_groups_per_var[q]*N_groups_per_var[q];                      
      block_indexes[1+q] = N_groups_per_var[q]*N_groups_per_var[q] + block_indexes[q];  
    }

    for(q in 1: N_group_vars){
      for(k in 1: max_N_groups){
        N_per_group[k, q] = 0;   
      }
    }

    for(q in 1: N_group_vars){
      for(i in 1:N_id){
        N_per_group[block_set[i,q],q] += 1;
      }
    }

    if (N_params[1] > 1) {
        for(i in 2:N_params[1]) focal_individual_predictors[ , i-1] = focal_set[,i];  
    }
    if (N_params[2] > 1) {
        for(i in 2:N_params[2]) target_individual_predictors[ , i-1] = target_set[,i];  
    }
    if (N_params[3] > 1) {
        for(i in 2:N_params[3]) flat_dyad_preds[ , i-1] = dyad_set[,i];  
    }
}

parameters {
    vector[block_param_size] block_effects;

    // Sender-receiver effects
    vector<lower=0>[2] sr_sigma;  
    cholesky_factor_corr[2] sr_L;
    matrix[2, N_id] z_sr;

    // Dyadic effects
    real<lower=0> dr_sigma;     
    cholesky_factor_corr[2] dr_L;
    matrix[2, N_dyads] z_dr;

    // Effects of covariate
    vector[N_params[1]-1] focal_effects;
    vector[N_params[2]-1] target_effects;
    vector[N_params[3]-1] dyad_effects;

    real<lower=0> error_sigma;       
}

model {
    array[N_group_vars] matrix[max_N_groups, max_N_groups] B;
    
    // Transform vector of block effects into matrices
    for(q in 1:N_group_vars){
      B[q, 1:N_groups_per_var[q], 1:N_groups_per_var[q]] = to_matrix(block_effects[(block_indexes[q]+1):(block_indexes[q+1])], N_groups_per_var[q], N_groups_per_var[q]);
    }

    // Priors on B
    for (q in 1:N_group_vars) {
        for (i in 1:N_groups_per_var[q]) {
            for (j in 1:N_groups_per_var[q]) {
                if (i == j) {
                    B[q, i, j] ~ normal(logit(priors[10, 1] / sqrt(N_per_group[i, q])), priors[10, 2]);
                } else {
                    B[q, i, j] ~ normal(logit(priors[11, 1] / sqrt(N_per_group[i, q] * 0.5 + N_per_group[j, q] * 0.5)), priors[11, 2]);
                }
            }
        }
    }

    focal_effects ~ normal(priors[12,1], priors[12,2]);
    target_effects ~ normal(priors[13,1], priors[13,2]);
    dyad_effects ~ normal(priors[14,1], priors[14,2]);
    error_sigma ~ normal(priors[23,1], priors[23,2]);

    to_vector(z_sr) ~ std_normal();
    sr_sigma ~ normal(priors[15,1], priors[15,2]);    
    sr_L ~ lkj_corr_cholesky(priors[17,1]);

    to_vector(z_dr) ~ std_normal();
    dr_sigma ~ normal(priors[16,1], priors[16,2]);
    dr_L ~ lkj_corr_cholesky(priors[18,1]);

    matrix[N_id, 2] sr_eff = (diag_pre_multiply(sr_sigma, sr_L) * z_sr)';
    matrix[N_dyads, 2] dr_eff = (diag_pre_multiply(rep_vector(dr_sigma, 2), dr_L) * z_dr)';

    vector[N_id] f_eff = rep_vector(0.0, N_id);
    if (N_params[1] > 1) f_eff = focal_individual_predictors * focal_effects;
    
    vector[N_id] t_eff = rep_vector(0.0, N_id);
    if (N_params[2] > 1) t_eff = target_individual_predictors * target_effects;
    
    vector[N_obs] d_eff = rep_vector(0.0, N_obs);
    if (N_params[3] > 1) d_eff = flat_dyad_preds * dyad_effects;

    vector[N_obs] linear_model = f_eff[sender] + t_eff[receiver] + sr_eff[sender, 1] + sr_eff[receiver, 2] + d_eff;

    for (k in 1:N_obs) {
        linear_model[k] += dr_eff[dyad_id[k], dyad_dir[k]];
        for (q in 1:N_group_vars) {
            linear_model[k] += B[q, block_set[sender[k], q], block_set[receiver[k], q]];
        }
    }

    if (outcome_mode == 1) {
        if (link_mode == 1) outcomes ~ bernoulli_logit(linear_model);
        if (link_mode == 2) outcomes ~ bernoulli(Phi(linear_model));
    } else if (outcome_mode == 2) {
        if (link_mode == 1) outcomes ~ binomial_logit(exposure, linear_model);
        if (link_mode == 2) outcomes ~ binomial(exposure, Phi(linear_model));
    } else if (outcome_mode == 3) {
        outcomes ~ poisson_log(linear_model);
    } else if (outcome_mode == 4) {
        outcomes_real ~ normal(linear_model, error_sigma);
    }
}

generated quantities {
    vector[export_network == 1 ? N_obs : 0] p;

    if (export_network == 1) {
        matrix[2, N_id] sr_base = diag_pre_multiply(sr_sigma, sr_L) * z_sr;
        matrix[2, N_dyads] dr_base = diag_pre_multiply(rep_vector(dr_sigma, 2), dr_L) * z_dr;
        
        array[N_group_vars] matrix[max_N_groups, max_N_groups] B;
        for(q in 1:N_group_vars){
          B[q, 1:N_groups_per_var[q], 1:N_groups_per_var[q]] = to_matrix(block_effects[(block_indexes[q]+1):(block_indexes[q+1])], N_groups_per_var[q], N_groups_per_var[q]);
        }

        for (k in 1:N_obs) {
            real f_val = 0.0;
            real t_val = 0.0;
            real d_val = 0.0;
            real b_val = 0.0;
            
            if (N_params[1] > 1) f_val = dot_product(focal_effects, to_vector(focal_individual_predictors[sender[k]]));
            if (N_params[2] > 1) t_val = dot_product(target_effects, to_vector(target_individual_predictors[receiver[k]]));
            if (N_params[3] > 1) d_val = dot_product(dyad_effects, to_vector(flat_dyad_preds[k]));
            for (q in 1:N_group_vars) {
                b_val += B[q, block_set[sender[k], q], block_set[receiver[k], q]];
            }

            real lp = b_val + f_val + t_val 
                      + sr_base[1, sender[k]] + sr_base[2, receiver[k]] 
                      + d_val + dr_base[dyad_dir[k], dyad_id[k]];

            if (outcome_mode == 1 || outcome_mode == 2) {
                p[k] = (link_mode == 1) ? inv_logit(lp) : Phi(lp);
            } else if (outcome_mode == 3) {
                p[k] = exp(lp);
            } else if (outcome_mode == 4) {
                p[k] = lp;
            }
        }
    }
}
```
