## Appendix 1, 2, 3, and 4 for "BayesForge: A Bayesian Inference library for Python, R, and Julia": fit_comparison.py

```
import numpy as np
import pandas as pd
import matplotlib.pyplot as plt
import os
import sys
from cmdstanpy import CmdStanModel

# Force non-interactive matplotlib backend
plt.switch_backend('Agg')

# Setup device and BI model
from BayesForge import bf
import jax.numpy as jnp
m = bf('cpu', rand_seed=False)

print("Step 1: Simulating data 100% in Python (matching SIM.py structure)...")
sys.stdout.flush()

N_nodes = 50 
N_focal_vars = 4
N_target_vars = 4
N_dyad_vars = 3

np.random.seed(1)

# Generate individual (focal/target) predictors
wide_focal_np = np.random.normal(0, 1, size=(N_nodes, N_focal_vars))
wide_target_np = np.random.normal(0, 1, size=(N_nodes, N_target_vars))

# Generate dyadic predictors (N x N matrices)
dyadic_predictors_mat = np.random.binomial(1, 0.3, size=(N_nodes, N_nodes, N_dyad_vars))
for k in range(N_dyad_vars):
    np.fill_diagonal(dyadic_predictors_mat[:, :, k], 0)
# Make the 3rd dyadic predictor the interaction of the first two
dyadic_predictors_mat[:, :, 2] = dyadic_predictors_mat[:, :, 0] * dyadic_predictors_mat[:, :, 1]

# Convert dyadic predictors to edgelist (N_dyads, 2, N_dyad_vars)
dyadic_predictors_edgl = jnp.stack([m.net.mat_to_edgl(dyadic_predictors_mat[:, :, k]) for k in range(N_dyad_vars)], axis=2)

# Generate block groups
Any_np = np.zeros(N_nodes, dtype=int)
Merica_np = np.random.choice([0, 1, 2], size=(N_nodes,), p=[0.25, 0.5625, 0.1875])

N_grp_Any = 1
N_grp_Merica = 3
N_by_grp_Any = np.array([N_nodes])
N_by_grp_Merica = np.array([np.sum(Merica_np == i) for i in range(N_grp_Merica)])

# Simulate network outcomes using bi's generative APIs
def sim_network(dyadic_predictors, focal_individual_predictors, target_individual_predictors, Any, Merica):
    from BayesForge import bf
    m_sim = bf()
    B_intercept = m_sim.net.block_model(Any, 1, jnp.array(N_by_grp_Any), sample=True, name="intercept")
    B_category = m_sim.net.block_model(Merica, 3, jnp.array(N_by_grp_Merica), sample=True, name="category")
    sr = m_sim.net.sender_receiver(
        focal_individual_predictors, 
        target_individual_predictors, 
        s_mu=0.4, r_mu=-0.4, sample=True)
    dr = m_sim.net.dyadic_effect(dyadic_predictors, d_sd=2.5, sample=True)
    logits = B_intercept + B_category + sr + dr
    return m_sim.dist.bernoulli(logits=logits, sample=True)

# Generate simulated network (N_dyads, 2)
Any_jnp = jnp.array(Any_np, dtype=jnp.int32)
Merica_jnp = jnp.array(Merica_np, dtype=jnp.int32)
network_edgl = sim_network(
    dyadic_predictors_edgl, 
    jnp.array(wide_focal_np), 
    jnp.array(wide_target_np), 
    Any_jnp, 
    Merica_jnp
)

N_dyads = network_edgl.shape[0]
N_obs = N_dyads * 2
print(f"Data successfully simulated. Dyads: {N_dyads}, Observations: {N_obs}")
sys.stdout.flush()

# Prepare variables for BI wide model fitting
wide_network_edgl = network_edgl
wide_dyad_edgl = dyadic_predictors_edgl
wide_sender_preds = jnp.array(wide_focal_np)
wide_receiver_preds = jnp.array(wide_target_np)

print("Step 2: Fitting model_srm_wide in BF...")
sys.stdout.flush()

from BayesForge.Network.model_effects import Neteffect

def model_srm_wide(network_edgl, dyadic_predictors, sender_predictors, receiver_predictors, Any, Merica, N_by_grp_Any, N_by_grp_Merica):
    m2_inner = bf()
    B_any    = Neteffect.block_model(Any,    1,    jnp.array(N_by_grp_Any),    name='intercept')
    B_Merica = Neteffect.block_model(Merica, 3,    jnp.array(N_by_grp_Merica), name='Merica')
    sr = m2_inner.net.sender_receiver(sender_predictors, receiver_predictors)
    dr = m2_inner.net.dyadic_effect(dyadic_predictors)
    m2_inner.dist.bernoulli(logits=B_any + B_Merica + sr + dr, obs=network_edgl, name='network_edgl')

m2 = bf('cpu')
m2.data_on_model = dict(
    network_edgl=wide_network_edgl, 
    dyadic_predictors=wide_dyad_edgl, 
    sender_predictors=wide_sender_preds, 
    receiver_predictors=wide_receiver_preds, 
    Any=Any_jnp, 
    Merica=Merica_jnp,
    N_by_grp_Any=N_by_grp_Any,
    N_by_grp_Merica=N_by_grp_Merica
)
m2.fit(model_srm_wide, num_samples=1000, num_warmup=1000, num_chains=1)
bi_samples = m2.posteriors

print("Step 3: Preparing data and fitting STAN2.stan...")
sys.stdout.flush()

# Construct dyad index representation
# Standard row-major indexing off-diagonal:
urows, ucols = np.triu_indices(N_nodes, k=1)
long_ids_int = np.stack([urows, ucols], axis=1) # Corrected to urows, ucols to match BI's mat_to_edgl

sender = np.concatenate([long_ids_int[:, 0], long_ids_int[:, 1]]) + 1
receiver = np.concatenate([long_ids_int[:, 1], long_ids_int[:, 0]]) + 1
dyad_id = np.concatenate([np.arange(1, N_dyads + 1), np.arange(1, N_dyads + 1)])
dyad_dir = np.concatenate([np.ones(N_dyads, dtype=int), np.full(N_dyads, 2, dtype=int)])
outcomes_srm2 = np.concatenate([network_edgl[:, 0], network_edgl[:, 1]])

# Format predictor matrices with intercepts for Stan
focal_set_stan = np.column_stack([np.ones(N_nodes), wide_focal_np])
target_set_stan = np.column_stack([np.ones(N_nodes), wide_target_np])

# Flatten dyadic_predictors_edgl along dyads & directions, adding intercept column 0
flat_dyad_preds = np.concatenate([dyadic_predictors_edgl[:, 0, :], dyadic_predictors_edgl[:, 1, :]], axis=0)
dyad_set_stan = np.column_stack([np.ones(N_obs), flat_dyad_preds])

# Block set (1-based category indices)
block_set_stan = np.column_stack([Any_np + 1, Merica_np + 1])

# Priors matching STRAND defaults
priors = np.zeros((23, 2))
p_data = [
    [-3.00, 1.5], [3.00, 1.5], [-1.50, 1.0], [1.00, 0.0], [1.00, 0.0], [1.00, 0.0],
    [0.00, 2.5], [0.00, 2.5], [0.00, 2.5], [0.10, 2.5], [0.01, 2.5], [0.00, 2.5],
    [0.00, 2.5], [0.00, 2.5], [0.00, 2.5], [0.00, 2.5], [2.50, 0.0], [2.50, 0.0],
    [1.50, 0.0], [3.00, 1.0], [2.00, 0.0], [3.00, 12.0], [0.00, 2.5]
]
for i, row in enumerate(p_data):
    priors[i] = row

stan_data = {
    'N_networktypes': 1,
    'N_id': N_nodes,
    'N_dyads': N_dyads,
    'N_obs': N_obs,
    'N_responses': 1,
    'N_params': [wide_focal_np.shape[1] + 1, wide_target_np.shape[1] + 1, flat_dyad_preds.shape[1] + 1],
    'sender': sender.tolist(),
    'receiver': receiver.tolist(),
    'dyad_id': dyad_id.tolist(),
    'dyad_dir': dyad_dir.tolist(),
    'outcomes': outcomes_srm2.astype(int).tolist(),
    'outcomes_real': outcomes_srm2.astype(float).tolist(),
    'exposure': np.ones(N_obs, dtype=int).tolist(),
    'N_group_vars': 2,
    'max_N_groups': 3,
    'N_groups_per_var': [1, 3],
    'block_set': block_set_stan.tolist(),
    'focal_set': focal_set_stan.tolist(),
    'target_set': target_set_stan.tolist(),
    'dyad_set': dyad_set_stan.tolist(),
    'priors': priors.tolist(),
    'export_network': 0,
    'outcome_mode': 1,
    'link_mode': 1
}

# Compile and run STAN2 model
script_dir = os.path.dirname(os.path.abspath(__file__))
stan_file = os.path.join(script_dir, 'benchmark_suite', 'STAN2.stan')
exe_file = os.path.join(script_dir, 'benchmark_suite', 'STAN2')

print("Loading/compiling STAN2.stan...")
sm = CmdStanModel(stan_file=stan_file)

fit = sm.sample(data=stan_data, iter_sampling=1000, iter_warmup=1000, chains=1, show_progress=False)
stan_samples = fit.stan_variables()

print("\nStep 3.5: Preparing data and fitting original_srm.stan...")
sys.stdout.flush()

# Format outcomes matrix for original_srm.stan
Y = np.zeros((N_nodes, N_nodes), dtype=int)
for d in range(N_dyads):
    u = urows[d]
    v = ucols[d]
    Y[u, v] = int(network_edgl[d, 0])
    Y[v, u] = int(network_edgl[d, 1])

outcomes_3d = Y.reshape(N_nodes, N_nodes, 1).tolist()
outcomes_real_3d = Y.reshape(N_nodes, N_nodes, 1).astype(float).tolist()
exposure_3d = np.ones((N_nodes, N_nodes, 1), dtype=int).tolist()
mask_3d = np.zeros((N_nodes, N_nodes, 1), dtype=int).tolist()

dyad_set_original = np.zeros((N_nodes, N_nodes, N_dyad_vars + 1))
dyad_set_original[:, :, 0] = 1.0  # ignored layer
dyad_set_original[:, :, 1:] = dyadic_predictors_mat

original_stan_data = {
    'N_networktypes': 1,
    'N_id': N_nodes,
    'N_responses': 1,
    'N_params': [wide_focal_np.shape[1] + 1, wide_target_np.shape[1] + 1, N_dyad_vars + 1],
    'outcomes': outcomes_3d,
    'outcomes_real': outcomes_real_3d,
    'exposure': exposure_3d,
    'mask': mask_3d,
    'focal_set': focal_set_stan.tolist(),
    'target_set': target_set_stan.tolist(),
    'dyad_set': dyad_set_original.tolist(),
    'priors': priors.tolist(),
    'export_network': 0,
    'outcome_mode': 1,
    'link_mode': 1,
    # Block structure variables for updated original_srm.stan
    'N_group_vars': 2,
    'max_N_groups': 3,
    'N_groups_per_var': [1, 3],
    'block_set': block_set_stan.tolist()
}

original_stan_file = os.path.join(script_dir, 'original_srm.stan')
original_exe_file = os.path.join(script_dir, 'original_srm')

print("Loading/compiling original_srm.stan...")
sm_orig = CmdStanModel(stan_file=original_stan_file)

fit_orig = sm_orig.sample(data=original_stan_data, iter_sampling=1000, iter_warmup=1000, chains=1, show_progress=False)
orig_samples = fit_orig.stan_variables()

print("Step 3.7: Skipped STRAND NumPyro")
sys.stdout.flush()

RUN_STRAND = False

if RUN_STRAND:
    import os
    os.environ['RETICULATE_PYTHON'] = '/home/sosa/.virtualenvs/BayesInference/bin/python'
    
    import rpy2.robjects as ro
    from rpy2.robjects import numpy2ri, pandas2ri
    from rpy2.robjects.conversion import localconverter
    
    # Build outcome network matrix
    outcome_mat = np.zeros((N_nodes, N_nodes), dtype=int)
    for d in range(N_dyads):
        u = urows[d]
        v = ucols[d]
        outcome_mat[u, v] = int(network_edgl[d, 0])
        outcome_mat[v, u] = int(network_edgl[d, 1])
    
    # Exposure matrix
    exposure_mat = np.ones((N_nodes, N_nodes), dtype=int)
    
    # Dyadic covariates
    Kinship = dyadic_predictors_mat[:, :, 0]
    Dominant = dyadic_predictors_mat[:, :, 1]
    
    # Individual covariates (separate focal and target for STRAND)
    Mass_f = wide_focal_np[:, 0]
    Age_f = wide_focal_np[:, 1]
    Love_f = wide_focal_np[:, 2]
    Fire_f = wide_focal_np[:, 3]
    
    Mass_t = wide_target_np[:, 0]
    Age_t = wide_target_np[:, 1]
    Love_t = wide_target_np[:, 2]
    Fire_t = wide_target_np[:, 3]
    
    with localconverter(ro.default_converter + numpy2ri.converter + pandas2ri.converter):
        ro.globalenv['outcome_mat'] = outcome_mat
        ro.globalenv['exposure_mat'] = exposure_mat
        ro.globalenv['Kinship'] = Kinship
        ro.globalenv['Dominant'] = Dominant
        ro.globalenv['Mass_f'] = Mass_f
        ro.globalenv['Age_f'] = Age_f
        ro.globalenv['Love_f'] = Love_f
        ro.globalenv['Fire_f'] = Fire_f
        ro.globalenv['Mass_t'] = Mass_t
        ro.globalenv['Age_t'] = Age_t
        ro.globalenv['Love_t'] = Love_t
        ro.globalenv['Fire_t'] = Fire_t
        ro.globalenv['Merica_np'] = Merica_np
    
        ro.r('library(STRAND)')
        ro.r("""
        name_vec = paste0("Ind_", 1:50)
        rownames(outcome_mat) = colnames(outcome_mat) = name_vec
        rownames(exposure_mat) = colnames(exposure_mat) = name_vec
        rownames(Kinship) = colnames(Kinship) = name_vec
        rownames(Dominant) = colnames(Dominant) = name_vec
    
        individual = data.frame(
          Mass_f=Mass_f, Age_f=Age_f, Love_f=Love_f, Fire_f=Fire_f,
          Mass_t=Mass_t, Age_t=Age_t, Love_t=Love_t, Fire_t=Fire_t
        )
        rownames(individual) = name_vec
    
        groups_f = data.frame(Merica=as.factor(Merica_np + 1))
        rownames(groups_f) = name_vec
    
        model_dat = make_strand_data(
          outcome=list(Association = outcome_mat),  
          block_covariates=groups_f, 
          individual_covariates=individual, 
          dyadic_covariates=list(Kinship=Kinship, Dominant=Dominant),  
          outcome_mode="bernoulli", 
          link_mode="logit",
          exposure=list(Association = exposure_mat)
        )
    
        fit_numpyro = fit_block_plus_social_relations_model(
          data=model_dat,
          block_regression = ~ Merica,
          focal_regression = ~ Mass_f + Age_f + Love_f + Fire_f,
          target_regression = ~ Mass_t + Age_t + Love_t + Fire_t,
          dyad_regression = ~ Kinship*Dominant,
          mode="numpyro",
          mcmc_parameters = list(seed = 1, chains = 1, iter_warmup = 1000, iter_sampling = 1000)
        )
        res_stan = summarize_strand_results(fit_numpyro)
        samps = res_stan$samples$srm_model_samples
        """)
    
    # Extract STRAND samples in Python
    strand_samps = ro.globalenv['samps']
    strand_block_params = strand_samps.rx2('block_parameters')
    strand_focal_target_sd = np.array(strand_samps.rx2('focal_target_sd'))
    strand_focal_coeffs = np.array(strand_samps.rx2('focal_coeffs'))
    strand_target_coeffs = np.array(strand_samps.rx2('target_coeffs'))
    strand_dyadic_coeffs = np.array(strand_samps.rx2('dyadic_coeffs'))
    strand_dyadic_sd = np.array(strand_samps.rx2('dyadic_sd')).flatten()
    
    # Retrieve block matrices
    strand_b_intercept = np.array(strand_block_params.rx2(1)).reshape(-1)
    strand_b_Merica = np.array(strand_block_params.rx2(2))
else:
    strand_mapped = {}


print("\nStep 4: Printing Posterior Comparison Table...")
print("BI posteriors keys:", list(bi_samples.keys()))
print("Stan (Block) posteriors keys:", list(stan_samples.keys()))
print("Stan (Original) posteriors keys:", list(orig_samples.keys()))
sys.stdout.flush()

# Map BI and Stan parameters for exact side-by-side comparison
bi_mapped = {
    'Block Intercept': bi_samples['b_intercept'].reshape(bi_samples['b_intercept'].shape[0], -1)[:, 0],
    **{f'Block Merica {i+1}-{j+1}': np.array([m[i, j] for m in bi_samples['b_Merica']]) for i in range(3) for j in range(3)},
    **{f'Sender covariate {i+1}': bi_samples['sender_effects'][:, i] for i in range(4)},
    **{f'Receiver covariate {i+1}': bi_samples['receiver_effects'][:, i] for i in range(4)},
    **{f'Dyad covariate {i+1}': bi_samples['dyad_effects'][:, i] for i in range(3)},
    'Sender SD': bi_samples['sr_sigma'][:, 0],
    'Receiver SD': bi_samples['sr_sigma'][:, 1],
    'Dyad SD': bi_samples['dr_sigma']
}

stan_mapped = {
    'Block Intercept': stan_samples['block_effects'][:, 0],
    **{f'Block Merica {i+1}-{j+1}': stan_samples['block_effects'][:, 1 + (j * 3 + i)] for i in range(3) for j in range(3)},
    **{f'Sender covariate {i+1}': stan_samples['focal_effects'][:, i] for i in range(4)},
    **{f'Receiver covariate {i+1}': stan_samples['target_effects'][:, i] for i in range(4)},
    **{f'Dyad covariate {i+1}': stan_samples['dyad_effects'][:, i] for i in range(3)},
    'Sender SD': stan_samples['sr_sigma'][:, 0],
    'Receiver SD': stan_samples['sr_sigma'][:, 1],
    'Dyad SD': stan_samples['dr_sigma']
}

orig_mapped = {
    'Block Intercept': orig_samples['block_effects'][:, 0],
    **{f'Block Merica {i+1}-{j+1}': orig_samples['block_effects'][:, 1 + (j * 3 + i)] for i in range(3) for j in range(3)},
    **{f'Sender covariate {i+1}': orig_samples['focal_effects'][:, i] for i in range(4)},
    **{f'Receiver covariate {i+1}': orig_samples['target_effects'][:, i] for i in range(4)},
    **{f'Dyad covariate {i+1}': orig_samples['dyad_effects'][:, i] for i in range(3)},
    'Sender SD': orig_samples['sr_sigma'][:, 0],
    'Receiver SD': orig_samples['sr_sigma'][:, 1],
    'Dyad SD': orig_samples['dr_sigma']
}

strand_mapped = {}

records = []
for key in bi_mapped:
    bi_mean, bi_std = np.mean(bi_mapped[key]), np.std(bi_mapped[key])
    stan_mean, stan_std = np.mean(stan_mapped[key]), np.std(stan_mapped[key])
    orig_mean, orig_std = np.mean(orig_mapped[key]), np.std(orig_mapped[key])
        
    records.append({
        'Parameter': key,
        'BI Mean': f"{bi_mean: .4f}",
        'BI Std': f"{bi_std: .4f}",
        'Stan Blk Mean': f"{stan_mean: .4f}",
        'Stan Blk Std': f"{stan_std: .4f}",
        'Stan Orig Mean': f"{orig_mean: .4f}",
        'Stan Orig Std': f"{orig_std: .4f}"
    })

df_comparison = pd.DataFrame(records)
print(df_comparison.to_string(index=False))
sys.stdout.flush()

print("\nStep 5: Generating Forest Plot 'stan V BF.png'...")
sys.stdout.flush()

def get_stats(data):
    m = np.mean(data)
    hpd = np.percentile(data, [2.5, 97.5])
    return m, np.array([[m - hpd[0]], [hpd[1] - m]])

params = list(bi_mapped.keys())
n = len(params)
y = np.arange(n)

# Extract stats for all models
bi_stats = [get_stats(bi_mapped[key]) for key in params]
stan_stats = [get_stats(stan_mapped[key]) for key in params]
orig_stats = [get_stats(orig_mapped[key]) for key in params]

bi_means = [s[0] for s in bi_stats]
bi_errs = np.hstack([s[1] for s in bi_stats])

stan_means = [s[0] for s in stan_stats]
stan_errs = np.hstack([s[1] for s in stan_stats])

orig_means = [s[0] for s in orig_stats]
orig_errs = np.hstack([s[1] for s in orig_stats])

colors = {'stan': '#ff7f0e', 'bi': '#1f77b4', 'orig': '#2ca02c', 'strand': '#d62728'}

plt.figure(figsize=(14, n * 0.55 + 2))
plt.axvline(0, color='k', ls='--', alpha=0.3)

plt.errorbar(orig_means, y - 0.2, xerr=orig_errs, fmt='^', label='Stan (Original SRM)', color=colors['orig'], capsize=3)
plt.errorbar(stan_means, y + 0.2, xerr=stan_errs, fmt='o', label='Stan (STAN2)', color=colors['stan'], capsize=3)
plt.errorbar(bi_means, y, xerr=bi_errs, fmt='s', label='BF', color=colors['bi'], capsize=3)

plt.yticks(y, params, fontsize=16)
plt.xticks(fontsize=16)
plt.xlabel("Estimate", fontsize=20, fontweight='bold')
plt.ylabel("Parameter", fontsize=20, fontweight='bold')
legend = plt.legend(
    title="Backend", title_fontsize=20,
    loc="upper center", bbox_to_anchor=(0.5, 1.08),
    ncol=3, frameon=False, fontsize=16,
    handlelength=1.0, handleheight=1.0,
    markerscale=1.0,
)
legend.get_title().set_fontweight("bold")
plt.title('SRM Posterior Comparison: Stan vs. BF', fontsize=24, fontweight='bold', pad=70)
plt.tight_layout()

# Save plot relative to script directory
plot_path = os.path.join(script_dir, 'stan V BF.png')
plt.savefig(plot_path, dpi=150)
plt.close()
print(f"Forest plot saved successfully to {plot_path}")
sys.stdout.flush()
```
