## Appendix 1, 2, 3, and 4 for "BayesForge: A Bayesian Inference library for Python, R, and Julia": original_srm.stan

```
data{   
  //# Array dimension variables                                   
    int N_id;                                  //# Number of people                                                                                                   
    int N_responses;                           //# Number of outcome networks
    array[3] int N_params;                     //# Number of focal, target, and dyadic predictors

  //# Block predictor variables 
    int N_group_vars;                          //# Number of block structure variables
    int max_N_groups;                          //# Max number of group labels in any variable
    array[N_group_vars] int N_groups_per_var;  //# Number of group labels, per variable type
    array[N_id, N_group_vars] int block_set;   //# Dataframe holding the group ID codes for each person (rows) for each variable type (cols)

  //# Focal, target, and dyadic predictor variables                                                                                                      
    matrix[N_id, N_params[1]] focal_set;             //# Focal slash decider predictor variables    
    matrix[N_id, N_params[2]] target_set;            //# Target slash alter predictor variables
    array[N_id, N_id, N_params[3]] real dyad_set;    //# Dyadic predictor variables

  //# Outcome and exposure data
    array[N_id,N_id,N_responses] int outcomes;       //# Outcome network of binary ties
    array[N_id,N_id,N_responses] real outcomes_real; //# Outcome network if real
    array[N_id,N_id,N_responses] int exposure;       //# Exposure for each outcome
    array[N_id,N_id,N_responses] int mask;           //# Mask for each outcome

  //# Accessory paramters 
    matrix[23, 2] priors;                       //# Priors in a matrix, see details in the make_priors() function
    int export_network;                         //# Controls export of predictions
    int outcome_mode;                           //# Are outcomes binomial
    int link_mode;
}

transformed data{
  //# Refactor to the first predictor slot, becuase it is unity
    matrix[N_id, N_params[1]-1] focal_predictors;           //# Same as focal_set without first column
    matrix[N_id, N_params[2]-1] target_predictors;          //# Same as target_set without first column
    array[N_id, N_id, N_params[3]-1] real dyad_predictors;  //# Same as dyad_set without first shelf

  //# Store some key indexes
    array[max_N_groups, N_group_vars] int N_per_group;      //# Number of people in each block-type for each group variable
    array[N_group_vars+1] int block_indexes;                //# The indexes of each block parameter when stored as a vector instead of ragged array
    int block_param_size;                                   //# Total number of block-level parameters

  //# Get size of parameters for block model
    block_param_size = 0;                             //# Start at zero
    block_indexes[1] = 0;                             //# Start at zero for first index 
    
    for(q in 1: N_group_vars){
     block_param_size += N_groups_per_var[q]*N_groups_per_var[q];                      //# Count up number of parameters in each K by K block matrix and add to total
     block_indexes[1+q] = N_groups_per_var[q]*N_groups_per_var[q] + block_indexes[q];  //# Create cummulative sum of block indices, by adding new sum to old sum
     }

  //# First fill with scrap
    for(q in 1: N_group_vars){
    for(k in 1: max_N_groups){
     N_per_group[k, q] = 0;   
     }}

  //# Now fill in real values
    for(q in 1: N_group_vars){
    for(i in 1:N_id){
     N_per_group[block_set[i,q],q] += 1;
     }}

  //# Make pruned data for predictor variables, by dropping first column
    if(N_params[1]>1){
     for(i in 2:N_params[1]){
     focal_predictors[ , i-1] = focal_set[,i];  
     }}

    if(N_params[2]>1){
     for(i in 2:N_params[2]){
     target_predictors[ , i-1] = target_set[,i];  
     }}

    if(N_params[3]>1){
     for(i in 2:N_params[3]){
     dyad_predictors[ , , i-1] = dyad_set[,,i];  
     }}
}

parameters{
    //# Block effects, stored as a vector to save space
    vector[block_param_size] block_effects;

    //# Variation of sender-receiver effects
    vector<lower=0>[2] sr_sigma;  
    cholesky_factor_corr[2] sr_L;
    array[N_id] vector[2] sr_raw;

    //# Variation of dyadic effects
    real<lower=0> dr_sigma;       
    cholesky_factor_corr[2] dr_L;
    matrix[N_id, N_id] dr_raw;

    //# Effects of covariate
    vector[N_params[1]-1] focal_effects;
    vector[N_params[2]-1] target_effects;
    vector[N_params[3]-1] dyad_effects; 

    //# Error in Gaussian model
    real<lower=0> error_sigma;      
}

transformed parameters{
    matrix[2*N_responses, 2*N_responses] G_corr; 
    matrix[2*N_responses, 2*N_responses] D_corr; 

    G_corr = tcrossprod(sr_L); 
    D_corr = tcrossprod(dr_L);  
}

model{
  //# Local storage to make code more readable
    array[N_id] vector[2] sr;                                  //# Sender and receiver effects
    matrix[N_id, N_id] dr;                                     //# Dyadic effects
    array[N_group_vars] matrix[max_N_groups, max_N_groups] B;  //# Block effects, in array form
    vector[N_group_vars] br;                                   //# Sum of block effects per dyad    
    vector[2] scrap;                                           //# Local storage                    
    
    //# The first step, is to transform the vector of block effects into a list of matrices
    for(q in 1:N_group_vars){
      B[q,1:N_groups_per_var[q], 1:N_groups_per_var[q]] = to_matrix(block_effects[(block_indexes[q]+1):(block_indexes[q+1])], N_groups_per_var[q], N_groups_per_var[q]);
    }

    //# Then put priors on B, which scale loosely with the block size
    for ( q in 1:N_group_vars ){
    for ( i in 1:N_groups_per_var[q] ){
        for ( j in 1:N_groups_per_var[q] ) {
            if ( i==j ) {
                B[q,i,j] ~ normal(logit(priors[10,1]/sqrt(N_per_group[i,q])), priors[10,2]);   //# transfers more likely within groups
            } else {
                B[q,i,j] ~ normal(logit(priors[11,1]/sqrt(N_per_group[i,q]*0.5 + N_per_group[j,q]*0.5)), priors[11,2]); //# transfers less likely between groups
            }
        }}
    }

    //# Priors on effects of covariates
     focal_effects ~ normal(priors[12,1], priors[12,2]);
     target_effects ~ normal(priors[13,1], priors[13,2]);
     dyad_effects ~ normal(priors[14,1], priors[14,2]);

     error_sigma ~ normal(priors[23,1], priors[23,2]);

    //# Sender-receiver priors for social relations model
    for(i in 1:N_id)
    sr_raw[i] ~ normal(0,1);
    sr_sigma ~ normal(priors[15,1], priors[15,2]);
    sr_L ~ lkj_corr_cholesky(priors[17,1]);

    for(i in 1:N_id){
     vector[2] sr_terms;

     sr_terms[1] = dot_product(focal_effects,  to_vector(focal_predictors[i]));
     sr_terms[2] = dot_product(target_effects,  to_vector(target_predictors[i]));  

     sr[i] = diag_pre_multiply(sr_sigma, sr_L) * sr_raw[i] + sr_terms;
     }

    //# Dyadic priors for social relations model
    to_vector(dr_raw) ~ normal(0,1);
    dr_sigma ~ normal(priors[16,1], priors[16,2]);
    dr_L ~ lkj_corr_cholesky(priors[18,1]);

    for(i in 1:(N_id-1)){
    for(j in (i+1):N_id){
     scrap[1] = dr_raw[i,j];
     scrap[2] = dr_raw[j,i];
     scrap = rep_vector(dr_sigma, 2) .* (dr_L*scrap);
     dr[i,j] = scrap[1] + dot_product(dyad_effects,  to_vector(dyad_predictors[i, j, ]));
     dr[j,i] = scrap[2] + dot_product(dyad_effects,  to_vector(dyad_predictors[j, i, ]));
     }}

    for(i in 1:N_id){
     dr[i,i] = -99; //# ignore this :)
    }

    //# likelihood
    for(i in 1:N_id){
     for(j in 1:N_id){
       if(i != j){
       if(mask[i,j,1]==0){
        for(q in 1:N_group_vars){
          br[q] = B[q,block_set[i,q], block_set[j,q]]; //# Extract all of the block components for this dyad
         }

      if(outcome_mode==1){
        if(link_mode==1){
         outcomes[i,j,1] ~ bernoulli_logit(sum(br) + sr[i,1] + sr[j,2] + dr[i,j]);  //# Then model the outcomes
        }
        if(link_mode==2){
         outcomes[i,j,1] ~ bernoulli(Phi(sum(br) + sr[i,1] + sr[j,2] + dr[i,j]));  //# Then model the outcomes
        }
       }

      if(outcome_mode==2){
        if(link_mode==1){
         outcomes[i,j,1] ~ binomial_logit(exposure[i,j,1], sum(br) + sr[i,1] + sr[j,2] + dr[i,j]);  //# Then model the outcomes
        }
        if(link_mode==2){
         outcomes[i,j,1] ~ binomial(exposure[i,j,1], Phi(sum(br) + sr[i,1] + sr[j,2] + dr[i,j]));  //# Then model the outcomes
        }
       }

      if(outcome_mode==3){
      outcomes[i,j,1] ~ poisson_log(sum(br) + sr[i,1] + sr[j,2] + dr[i,j]);  //# Then model the outcomes
       }

      if(outcome_mode==4){
      outcomes_real[i,j,1] ~ normal(sum(br) + sr[i,1] + sr[j,2] + dr[i,j], error_sigma);  //# Then model the outcomes
       }

       }
      }
     }}


 }


generated quantities{
    //# compute posterior prob of each network tie
    matrix[N_id*export_network, N_id*export_network] p;
    array[N_id*export_network] vector[2*export_network] sr;
    matrix[N_id*export_network, N_id*export_network] dr;
 
    if(export_network==1){                
     vector[2] terms;
     int tie;
     array[N_group_vars] matrix[max_N_groups, max_N_groups] B;

    for(i in 1:N_group_vars){
     B[i,1:N_groups_per_var[i], 1:N_groups_per_var[i]] = to_matrix(block_effects[(block_indexes[i]+1):(block_indexes[i+1])], N_groups_per_var[i], N_groups_per_var[i]);
    }
            
    for(i in 1:N_id){
     vector[2] sr_terms;

     sr_terms[1] = dot_product(focal_effects,  to_vector(focal_predictors[i]));
     sr_terms[2] = dot_product(target_effects,  to_vector(target_predictors[i]));  

     sr[i] = diag_pre_multiply(sr_sigma, sr_L) * sr_raw[i] + sr_terms;
     }

    for(i in 1:(N_id-1)){
    for(j in (i+1):N_id){
      vector[2] scrap;
      vector[N_group_vars] br1;
      vector[N_group_vars] br2;

      scrap[1] = dr_raw[i,j];
      scrap[2] = dr_raw[j,i];
      scrap = rep_vector(dr_sigma, 2) .* (dr_L*scrap);

       for(q in 1:N_group_vars){
        br1[q] = B[q,block_set[i,q], block_set[j,q]];
        br2[q] = B[q,block_set[j,q], block_set[i,q]];
         }

       dr[i,j] = scrap[1] + dot_product(dyad_effects,  to_vector(dyad_predictors[i, j, ])) + sum(br1);
       dr[j,i] = scrap[2] + dot_product(dyad_effects,  to_vector(dyad_predictors[j, i, ])) + sum(br2);
    }}

    for(i in 1:N_id){
      dr[i,i] = -99; //# ignore this :)
    }


    for ( i in 1:N_id ) {
        for ( j in 1:N_id ) {
            if ( i != j ) {
      // consider each possible state of true tie and compute prob of data
      if(outcome_mode==1){
        if(link_mode==1){
         p[i,j] = inv_logit( sr[i,1] + sr[j,2] + dr[i,j]);
        }
        if(link_mode==2){
         p[i,j] = Phi( sr[i,1] + sr[j,2] + dr[i,j]);
        }
       }

      if(outcome_mode==2){
        if(link_mode==1){
         p[i,j] = inv_logit( sr[i,1] + sr[j,2] + dr[i,j]);
        }
        if(link_mode==2){
         p[i,j] = Phi( sr[i,1] + sr[j,2] + dr[i,j]);
        }
       }

      if(outcome_mode==3){
       p[i,j] = exp(sr[i,1] + sr[j,2] + dr[i,j]);  
       }

      if(outcome_mode==4){
       p[i,j] = sr[i,1] + sr[j,2] + dr[i,j];  
       }
            }
        }//j
    }//i

  for ( i in 1:N_id ) {
   p[i,i] = 0; 
   }
 }
}
```
