## Appendix 1, 2, 3, and 4 for "BayesForge: A Bayesian Inference library for Python, R, and Julia": SIM.py

```
##############################
#### Tested on linux
##############################
# %%
import rpy2.robjects as ro
from rpy2.robjects import pandas2ri, numpy2ri
from rpy2.robjects.conversion import localconverter
from rpy2.robjects.vectors import (
    ListVector,
    FloatVector,
    FloatMatrix,
    IntVector,
    IntMatrix,
    StrVector,
)
from rpy2.rinterface import NULLType
import numpy as np
import jax.numpy as jnp
import matplotlib.pyplot as plt
from jax import vmap
from functools import partial
from model_effects import Neteffect
from BI import bi

m = bi("cpu")


# %% Run R simulation and NumPyro fit
with localconverter(ro.default_converter + numpy2ri.converter + pandas2ri.converter):
    ro.r(
        """
library(posterior)
library(igraph)
library(STRAND)
library(ggplot2)

set.seed(1)

# Make data
N_id = 50

# Covariates
Kinship = standardize_strand(rlkjcorr( 1 , N_id , eta=1.5 ))
Dominant = matrix(rbern(N_id*N_id), 0.2, nrow=N_id, ncol=N_id)
Mass = rbern(N_id, 0.4)
Age = rnorm(N_id, 0, 1)
Love = rbern(N_id, 0.4)
Fire = rbern(N_id, 0.6)

# Organize into list
dyadic_preds = array(NA,c(N_id,N_id,3))
dyadic_preds[,,1] = Kinship
dyadic_preds[,,2] = Dominant
dyadic_preds[,,3] = Kinship*Dominant

# Set effect sizes
sr_mu = c(0,0)  
sr_sigma = c(2.2, 1.7) 
sr_rho = -0.7
dr_mu = 0 
dr_sigma = 3.5
dr_rho= 0.8
sr_effects_1 = c(2, 1.5)
sr_effects_2 = c(-1.4, 1.0)
sr_effects_3 = c(1.4, -1.3)
sr_effects_4 = c(-0.9, 0)
dr_effects_1 = c(1.5, -1.9, 2.4)

# Block structure
group_probs_block_size = c(0.25, c(0.25, 0.25)*(1-0.25))

B_1 = matrix(-8,nrow=1,ncol=1)
B_2 = matrix(rnorm(9,0,3),nrow=3,ncol=3)

diag(B_2) = diag(B_2) + 3.5

B = list(B_1, B_2)

groups_1 = rep("Any",N_id) 
groups_2 = sample( c("Red","White","Blue") , size=N_id , replace=TRUE , prob=group_probs_block_size )

groups_f = data.frame(Intercept=factor(groups_1), Merica=factor(groups_2))
individual = data.frame(Mass = Mass, Age = Age, Love = Love, Fire = Fire)

G = simulate_sbm_plus_srm_network(N_id = N_id, 
                                  B = B, 
                                  V = 2,
                                  groups=data.frame(Intercept=as.numeric(factor(groups_1)), Merica=as.numeric(factor(groups_2))),                  
                                  sr_mu = sr_mu,  
                                  sr_sigma = sr_sigma, 
                                  sr_rho = sr_rho,
                                  dr_mu = dr_mu,  
                                  dr_sigma = dr_sigma, 
                                  dr_rho = dr_rho,
                                  error_sigma = 1.0,
                                  outcome_mode="bernoulli", 
                                  link_mode="logit",                 
                                  individual_predictors = individual,
                                  dyadic_predictors = dyadic_preds,
                                  individual_effects = cbind(sr_effects_1, sr_effects_2, sr_effects_3, sr_effects_4),
                                  dyadic_effects = dr_effects_1
)

name_vec = paste("Individual", 1:N_id)
rownames(G$network) = colnames(G$network) = name_vec
rownames(G$samps) = colnames(G$samps) = name_vec
rownames(Kinship) = colnames(Kinship) = name_vec
rownames(Dominant) = colnames(Dominant) = name_vec
rownames(groups_f) = name_vec
rownames(individual) = name_vec

model_dat = make_strand_data(outcome=list(Association = G$network),  
                             block_covariates=groups_f, 
                             individual_covariates=individual, 
                             dyadic_covariates=list(Kinship=Kinship, Dominant=Dominant),  
                             outcome_mode="bernoulli", 
                             link_mode="logit",
                             exposure=list(Association = G$samps)
)

fit_numpyro = fit_block_plus_social_relations_model(
  data=model_dat,
  block_regression = ~ Merica,
  focal_regression = ~ Mass + Age + Love + Fire,
  target_regression = ~ Mass + Age + Love + Fire,
  dyad_regression = ~ Kinship*Dominant,
  mode="numpyro",
  mcmc_parameters = list(seed = 1, chains = 1, iter_warmup = 1000, iter_sampling = 1000)
)

res_stan = summarize_strand_results(fit_numpyro)
# Merge SRM samples and Block samples
strand_posteriors = c(res_stan$samples$srm_model_samples, res_stan$samples$block_parameters)
save(fit_numpyro, model_dat, file = 'STRAND.Rdata')

# Regenerate exact arrays for Python loading
d    = fit_numpyro$data
N_id = d$N_id
locs = make_dyadic_edgelist(N_id)

long_focal_set  = focal_set_to_long(as.matrix(d$focal_set))
long_target_set = target_set_to_long(as.matrix(d$target_set))
long_dyad_set   = dyadic_set_to_long(d$dyad_set)

# For blocks
block_dat = block_set_to_dyadic_block_set(d$block_set, priors=d$priors)
long_block_set = dyadic_set_to_long(block_dat$Y)
block_mu       = block_dat$Mu
block_sigma    = block_dat$Sigma

# For outcomes and exposure
out_mat = if(is.list(d$outcomes)) d$outcomes[[1]] else d$outcomes
exp_mat = if(is.list(d$exposure)) d$exposure[[1]] else d$exposure
if(length(dim(out_mat)) == 3) out_mat = out_mat[,,1]
if(length(dim(exp_mat)) == 3) exp_mat = exp_mat[,,1]

long_outcome = array(NA, c(nrow(locs), 2, 1))
long_outcome[,1,1] = out_mat[as.matrix(locs[,c("Var2","Var1")])]
long_outcome[,2,1] = out_mat[as.matrix(locs[,c("Var1","Var2")])]

long_exposure = array(NA, c(nrow(locs), 2, 1))
long_exposure[,1,1] = exp_mat[as.matrix(locs[,c("Var2","Var1")])]
long_exposure[,2,1] = exp_mat[as.matrix(locs[,c("Var1","Var2")])]

# --- Wide-format arrays ---
wide_outcome  = out_mat
wide_exposure = exp_mat
wide_focal_mat  = as.matrix(d$focal_set)[, -1, drop=FALSE]
wide_target_mat = as.matrix(d$target_set)[, -1, drop=FALSE]
raw_dyad_list = d$dyad_set[-1]
if (length(raw_dyad_list) > 0) {
  wide_dyad_mat = simplify2array(lapply(raw_dyad_list, as.matrix))
} else {
  wide_dyad_mat = array(0, dim=c(N_id, N_id, 0))
}
Any    = as.integer(as.factor(groups_1)) - 1L
Merica = as.integer(as.factor(groups_2)) - 1L

save(long_focal_set, long_target_set, long_dyad_set, long_block_set, 
     block_mu, block_sigma, long_outcome, long_exposure, locs, 
     wide_outcome, wide_exposure, wide_focal_mat, wide_target_mat, wide_dyad_mat,
     Any, Merica,
     file = 'strand_arrays.RData')
"""
    )


# %% Python side: Load and prepare data
ro.r["load"]("strand_arrays.RData")
with localconverter(ro.default_converter + numpy2ri.converter):
    long_focal_np = np.array(ro.globalenv["long_focal_set"])
    long_target_np = np.array(ro.globalenv["long_target_set"])
    long_dyad_np = np.array(ro.globalenv["long_dyad_set"])
    long_block_np = np.array(ro.globalenv["long_block_set"])
    block_mu_np = np.array(ro.globalenv["block_mu"])
    block_sigma_np = np.array(ro.globalenv["block_sigma"])
    long_outcome_np = np.array(ro.globalenv["long_outcome"])
    long_exposure_np = np.array(ro.globalenv["long_exposure"])
    locs_raw = np.array(ro.globalenv["locs"])
    wide_focal_np = np.array(ro.globalenv["wide_focal_mat"])
    wide_target_np = np.array(ro.globalenv["wide_target_mat"])
    wide_dyad_np = np.array(ro.globalenv["wide_dyad_mat"])
    Any_np = np.array(ro.globalenv["Any"]).astype(int).ravel()
    Merica_np = np.array(ro.globalenv["Merica"]).astype(int).ravel()

# Determine expected number of dyads to handle rpy2's unpredictable array transpositions
N_id_r = int(np.array(ro.globalenv["N_id"])[0])
N_dyads_expected = N_id_r * (N_id_r - 1) // 2

if long_focal_np.shape[0] != N_dyads_expected:
    long_focal_np = long_focal_np.transpose(2, 1, 0)
if long_block_np.shape[0] != N_dyads_expected:
    long_block_np = long_block_np.transpose(2, 1, 0)
if long_outcome_np.shape[0] != N_dyads_expected:
    long_outcome_np = long_outcome_np.transpose(2, 1, 0)
if long_exposure_np.shape[0] != N_dyads_expected:
    long_exposure_np = long_exposure_np.transpose(2, 1, 0)


if locs_raw.dtype.names is not None:
    locs_2d = np.stack([locs_raw[name] for name in ["Var2", "Var1"]], axis=1)
else:
    locs_2d = locs_raw.T if locs_raw.shape[0] == 2 else locs_raw
long_ids_int = jnp.array(locs_2d) - 1

long_focal_set = jnp.array(long_focal_np[:, :, 1:])
long_target_set = jnp.array(long_target_np[:, :, 1:])
long_dyad_set = jnp.array(long_dyad_np[:, :, 1:])
long_block_set = jnp.array(long_block_np)
network = jnp.array(long_outcome_np[:, :, 0])
exposure = jnp.array(long_exposure_np[:, :, 0])
N_nodes = int(jnp.max(long_ids_int) + 1)
N_dyads = int(long_ids_int.shape[0])

wide_network_edgl = jnp.array(long_outcome_np[:, :, 0])
wide_dyad_edgl = long_dyad_set
wide_sender_preds = jnp.array(wide_focal_np)
wide_receiver_preds = jnp.array(wide_target_np)

Any = jnp.array(Any_np, dtype=jnp.int32)
Merica = jnp.array(Merica_np, dtype=jnp.int32)

_, N_per_grp_Any = jnp.unique(Any, return_counts=True)
_, N_per_grp_Merica = jnp.unique(Merica, return_counts=True)
N_grp_Any = int(N_per_grp_Any.shape[0])
N_grp_Merica = int(N_per_grp_Merica.shape[0])

# --- Block Priors: use strand_block_prior() to auto-slice STRAND's flat vectors ---
# This replaces all manual reshaping and ensures identical priors across N_id values.
# (Removed manual injection as block_model now programmatically aligns with STRAND)


# %% Model definitions
def model_srm_long_block(
    network,
    long_dyad_set,
    long_focal_set,
    long_target_set,
    long_block_set,
    long_ids_int,
    exposure,
    sample=False,
):
    m = bi()
    N_var_focal, N_var_target, N_var_dyad, N_var_block = (
        long_focal_set.shape[2],
        long_target_set.shape[2],
        long_dyad_set.shape[2],
        long_block_set.shape[2],
    )
    block_effects = m.dist.normal(
        loc=block_mu_np,
        scale=block_sigma_np,
        shape=(N_var_block,),
        sample=sample,
        name="block_effects",
    )
    focal_effects = m.dist.normal(
        loc=jnp.zeros(N_var_focal),
        scale=2.5,
        shape=(N_var_focal,),
        sample=sample,
        name="focal_effects",
    )
    target_effects = m.dist.normal(
        loc=jnp.zeros(N_var_target),
        scale=2.5,
        shape=(N_var_target,),
        sample=sample,
        name="target_effects",
    )
    dyad_effects = m.dist.normal(
        loc=jnp.zeros(N_var_dyad),
        scale=2.5,
        shape=(N_var_dyad,),
        sample=sample,
        name="dyad_effects",
    )
    sr_L = m.dist.lkj_cholesky(2, 2.5, name="sr_L", sample=sample)
    sr_sigma = m.dist.truncated_normal(
        loc=0.0, scale=2.5, low=0.0, shape=(2,), sample=sample, name="sr_sigma"
    )
    sr_raw = m.dist.normal(
        loc=jnp.zeros((2, N_nodes)),
        scale=1.0,
        shape=(2, N_nodes),
        sample=sample,
        name="sr_raw",
    )
    sr = jnp.transpose(jnp.matmul(jnp.diag(sr_sigma) @ sr_L, sr_raw))
    dr_L = m.dist.lkj_cholesky(2, 2.5, name="dr_L", sample=sample)
    dr_sigma = m.dist.truncated_normal(
        loc=0.0, scale=2.5, low=0.0, shape=(), sample=sample, name="dr_sigma"
    )
    dr_raw = m.dist.normal(
        loc=jnp.zeros((2, N_dyads)),
        scale=1.0,
        shape=(2, N_dyads),
        sample=sample,
        name="dr_raw",
    )
    dr = jnp.transpose(
        jnp.matmul(jnp.expand_dims(jnp.repeat(dr_sigma, 2), 1) * dr_L, dr_raw)
    )
    mu = (
        jnp.tensordot(long_block_set, block_effects, axes=1)
        + jnp.tensordot(long_focal_set, focal_effects, axes=1)
        + jnp.tensordot(long_target_set, target_effects, axes=1)
        + jnp.tensordot(long_dyad_set, dyad_effects, axes=1)
    )
    S_i, R_j, S_j, R_i = (
        sr[long_ids_int[:, 0], 0],
        sr[long_ids_int[:, 1], 1],
        sr[long_ids_int[:, 1], 0],
        sr[long_ids_int[:, 0], 1],
    )
    gr_long = jnp.stack([S_i + R_j, S_j + R_i], axis=1)
    m.dist.binomial(
        jnp.ones_like(network), logits=mu + dr + gr_long, obs=network, name="network"
    )


def model_srm_wide(
    network_edgl, dyadic_predictors, sender_predictors, receiver_predictors, Any, Merica
):
    """Wide-format SRM using the programmatically aligned block_model API."""
    m2_inner = bi()
    # Now calls the updated internal logic which matches STRAND exactly
    B_any = Neteffect.block_model(Any, N_grp_Any, N_per_grp_Any, name="intercept")
    B_Merica = Neteffect.block_model(
        Merica, N_grp_Merica, N_per_grp_Merica, name="Merica"
    )
    sr = m2_inner.net.sender_receiver(sender_predictors, receiver_predictors)
    dr = m2_inner.net.dyadic_effect(dyadic_predictors)
    m2_inner.dist.bernoulli(
        logits=B_any + B_Merica + sr + dr, obs=network_edgl, name="network_edgl"
    )


# %% Fit models
m.data_on_model = dict(
    long_block_set=long_block_set,
    long_dyad_set=long_dyad_set,
    long_focal_set=long_focal_set,
    long_target_set=long_target_set,
    long_ids_int=long_ids_int,
    network=network,
    exposure=exposure,
)
m.fit(model_srm_long_block, num_samples=1000, num_warmup=1000, num_chains=1)
m2 = bi("cpu")
m2.data_on_model = dict(
    network_edgl=wide_network_edgl,
    dyadic_predictors=wide_dyad_edgl,
    sender_predictors=wide_sender_preds,
    receiver_predictors=wide_receiver_preds,
    Any=Any,
    Merica=Merica,
)
m2.fit(model_srm_wide, num_samples=1000, num_warmup=1000, num_chains=1)


# %% Robust Comparison
def r_to_py(obj):
    if isinstance(obj, ListVector):
        if isinstance(obj.names, (NULLType, type(None))):
            return [r_to_py(el) for el in obj]
        names = [str(n) for n in obj.names]
        return {n: r_to_py(obj.rx2(n)) for n in names}
    if hasattr(obj, "dim") and not isinstance(obj.dim, (NULLType, type(None))):
        return np.array(obj).reshape(list(obj.dim), order="F")
    if isinstance(obj, (FloatVector, IntVector, StrVector)):
        return np.array(obj)
    return obj


def compare_results(
    bi_long_posteriors,
    strand_posteriors,
    bi_wide_posteriors=None,
    outpath="forest_plot.png",
):
    def get_stats(data):
        m = np.mean(data)
        hpd = np.percentile(data, [2.5, 97.5])
        return m, [m - hpd[0], hpd[1] - m]

    summaries = []
    # ── 1. Blocks ──
    arr_l_blk = (
        np.array(bi_long_posteriors["block_effects"]).reshape(
            np.array(bi_long_posteriors["block_effects"]).shape[0], -1
        )
        if "block_effects" in bi_long_posteriors
        else None
    )
    long_bi_idx = 0
    wide_b_keys = (
        sorted(
            [
                k
                for k in (bi_wide_posteriors.keys() if bi_wide_posteriors else [])
                if k.startswith("b_")
            ],
            key=str.lower,
        )
        if bi_wide_posteriors is not None
        else []
    )
    raw_block_list = strand_posteriors.get("block_parameters", [])
    if hasattr(raw_block_list, "__len__") and not isinstance(
        raw_block_list, np.ndarray
    ):
        block_list = list(raw_block_list)
    elif isinstance(raw_block_list, np.ndarray) and raw_block_list.dtype == object:
        block_list = [raw_block_list[i] for i in range(len(raw_block_list))]
    else:
        block_list = [raw_block_list] if raw_block_list is not None else []

    def _flatten_block(arr):
        arr = np.array(arr)
        if arr.ndim == 3:
            S, M, N = arr.shape
            return arr.transpose(0, 2, 1).reshape(S, M * N)
        return arr.reshape(arr.shape[0], -1)

    for b_idx, raw_s in enumerate(block_list):
        arr_s = _flatten_block(raw_s)
        n_params = arr_s.shape[1]
        wide_arr = (
            _flatten_block(bi_wide_posteriors[wide_b_keys[b_idx]])
            if (bi_wide_posteriors and b_idx < len(wide_b_keys))
            else None
        )
        for j in range(n_params):
            label = f"block{b_idx}[{j}]" if n_params > 1 else f"block{b_idx}"
            sm, se = get_stats(arr_s[:, j])
            mm, me = (
                get_stats(arr_l_blk[:, long_bi_idx])
                if (arr_l_blk is not None and long_bi_idx < arr_l_blk.shape[1])
                else (None, None)
            )
            long_bi_idx += 1
            wm, we = (
                get_stats(wide_arr[:, j])
                if (wide_arr is not None and j < wide_arr.shape[1])
                else (None, None)
            )
            summaries.append(
                {
                    "param": label,
                    "strand_m": sm,
                    "strand_err": se,
                    "long_m": mm,
                    "long_err": me,
                    "wide_m": wm,
                    "wide_err": we,
                }
            )

    # ── 2. SRM ──
    mapping = [
        (("focal_effects", "sender_effects"), "focal_coeffs"),
        (("target_effects", "receiver_effects"), "target_coeffs"),
        (("dyad_effects", "dyadic_effects"), "dyadic_coeffs"),
        (("sr_sigma",), "focal_target_sd"),
        (("sr_L",), "focal_target_L"),
        (("dr_sigma",), "dyadic_sd"),
        (("dr_L",), "dyadic_L"),
    ]
    for bi_keys, s_key in mapping:
        if s_key not in strand_posteriors:
            continue
        arr_s = np.array(strand_posteriors[s_key]).reshape(
            np.array(strand_posteriors[s_key]).shape[0], -1
        )

        def _find_bi(post_dict, keys):
            if post_dict is None:
                return None
            for k in keys:
                if k in post_dict:
                    return np.array(post_dict[k]).reshape(
                        np.array(post_dict[k]).shape[0], -1
                    )
            return None

        arr_l, arr_w = _find_bi(bi_long_posteriors, bi_keys), _find_bi(
            bi_wide_posteriors, bi_keys
        )
        for j in range(arr_s.shape[1]):
            sm, se = get_stats(arr_s[:, j])
            mm, me = (
                get_stats(arr_l[:, j])
                if (arr_l is not None and j < arr_l.shape[1])
                else (None, None)
            )
            wm, we = (
                get_stats(arr_w[:, j])
                if (arr_w is not None and j < arr_w.shape[1])
                else (None, None)
            )
            label = f"{bi_keys[0]}[{j}]" if arr_s.shape[1] > 1 else bi_keys[0]
            summaries.append(
                {
                    "param": label,
                    "strand_m": sm,
                    "strand_err": se,
                    "long_m": mm,
                    "long_err": me,
                    "wide_m": wm,
                    "wide_err": we,
                }
            )

    # ── 3. Forest Plot ──
    params = [d["param"] for d in summaries]
    n = len(params)
    y = np.arange(n)
    colors = {
        "strand": "#ff7f0e",
        "long": "#1f77b4",
        "wide": "#2ca02c",
        "unique": "#d62728",
    }
    plt.figure(figsize=(10, n * 0.45 + 2))
    plt.axvline(0, color="k", ls="--", alpha=0.3)
    plt.errorbar(
        [d["strand_m"] for d in summaries],
        y + 0.15,
        xerr=np.array([d["strand_err"] for d in summaries]).T,
        fmt="o",
        label="STRAND",
        color=colors["strand"],
        capsize=3,
    )
    plt.errorbar(
        [d["long_m"] if d["long_m"] is not None else np.nan for d in summaries],
        y,
        xerr=np.array(
            [
                d["long_err"] if d["long_err"] is not None else [[0], [0]]
                for d in summaries
            ]
        ).T,
        fmt="s",
        label="BI long",
        color=colors["long"],
        capsize=3,
    )
    if bi_wide_posteriors:
        plt.errorbar(
            [d["wide_m"] if d["wide_m"] is not None else np.nan for d in summaries],
            y - 0.15,
            xerr=np.array(
                [
                    d["wide_err"] if d["wide_err"] is not None else [[0], [0]]
                    for d in summaries
                ]
            ).T,
            fmt="^",
            label="BI wide",
            color=colors["wide"],
            capsize=3,
        )
    plt.yticks(y, params, fontsize=8)
    plt.legend()
    plt.title("SRM Posterior Comparison")
    plt.tight_layout()
    plt.savefig(outpath, dpi=150)
    plt.show()
    plt.close()

    # ── 4. Scatters ──
    def _panel(ax, x, y, xl, yl, col, m):
        if x is None or y is None or len(x) == 0 or len(y) == 0:
            return
        ax.scatter(x, y, color=col, alpha=0.6, s=25, marker=m)
        all_v = np.concatenate([x, y])
        lo, hi = all_v.min(), all_v.max()
        pad = (hi - lo) * 0.1
        ax.plot([lo - pad, hi + pad], [lo - pad, hi + pad], "k--", alpha=0.3)
        r = np.corrcoef(x, y)[0, 1]
        ax.set_xlabel(xl, fontsize=7)
        ax.set_ylabel(yl, fontsize=7)
        ax.set_title(f"r={r:.3f}", fontsize=8)

    # Nodal
    # Nodal
    if "focal_target_random_effects" in strand_posteriors:
        st_sr = np.array(strand_posteriors["focal_target_random_effects"])
        print(f"[DEBUG] STRAND nodal RE shape: {st_sr.shape}")
        if st_sr.ndim >= 2:
            if st_sr.ndim == 3:
                st_sr = st_sr.mean(axis=0)  # (N, 2)
            li_sr = (
                np.array(bi_long_posteriors["sr_raw"]).mean(axis=0).T
                if "sr_raw" in bi_long_posteriors
                else None
            )
            wi_sr = (
                np.array(bi_wide_posteriors["sr_raw"]).mean(axis=0).T
                if bi_wide_posteriors and "sr_raw" in bi_wide_posteriors
                else None
            )
            fig, axes = plt.subplots(2, 3, figsize=(14, 8), squeeze=False)
            for i, role in enumerate(["Sender", "Receiver"]):
                _panel(
                    axes[i, 0],
                    st_sr[:, i],
                    li_sr[:, i] if li_sr is not None else None,
                    "STRAND",
                    f"BI long {role}",
                    colors["long"],
                    "s",
                )
                _panel(
                    axes[i, 1],
                    st_sr[:, i],
                    wi_sr[:, i] if wi_sr is not None else None,
                    "STRAND",
                    f"BI wide {role}",
                    colors["wide"],
                    "^",
                )
                _panel(
                    axes[i, 2],
                    li_sr[:, i] if li_sr is not None else None,
                    wi_sr[:, i] if wi_sr is not None else None,
                    "BI long",
                    f"BI wide {role}",
                    colors["unique"],
                    "o",
                )
            plt.tight_layout()
            plt.savefig(outpath.replace(".png", "_nodal_re.png"), dpi=150)
            plt.show()
            plt.close()

    # Dyadic
    if "dyadic_random_effects" in strand_posteriors:
        st_dr = np.array(strand_posteriors["dyadic_random_effects"])
        print(f"[DEBUG] STRAND dyadic RE shape: {st_dr.shape}")
        if st_dr.ndim >= 2:
            if st_dr.ndim == 3:
                st_dr = st_dr.mean(axis=0)  # (N_dyads, 2)
            li_dr = (
                np.array(bi_long_posteriors["dr_raw"]).mean(axis=0).T
                if "dr_raw" in bi_long_posteriors
                else None
            )
            wi_dr = (
                np.array(bi_wide_posteriors["dr_raw"]).mean(axis=0).T
                if bi_wide_posteriors and "dr_raw" in bi_wide_posteriors
                else None
            )
            fig, axes = plt.subplots(2, 3, figsize=(14, 8), squeeze=False)
            for i, dlabel in enumerate(["i->j", "j->i"]):
                _panel(
                    axes[i, 0],
                    st_dr[:, i],
                    li_dr[:, i] if li_dr is not None else None,
                    "STRAND",
                    f"BI long {dlabel}",
                    colors["long"],
                    "s",
                )
                _panel(
                    axes[i, 1],
                    st_dr[:, i],
                    wi_dr[:, i] if wi_dr is not None else None,
                    "STRAND",
                    f"BI wide {dlabel}",
                    colors["wide"],
                    "^",
                )
                _panel(
                    axes[i, 2],
                    li_dr[:, i] if li_dr is not None else None,
                    wi_dr[:, i] if wi_dr is not None else None,
                    "BI long",
                    f"BI wide {dlabel}",
                    colors["unique"],
                    "o",
                )
            plt.tight_layout()
            plt.savefig(outpath.replace(".png", "_dyadic_re.png"), dpi=150)
            plt.show()
            plt.close()
        else:
            print(f"[WARNING] Skipping dyadic plot: unexpected shape {st_dr.shape}")


def compare_results_srm(
    bi_long_posteriors,
    strand_posteriors,
    bi_wide_posteriors=None,
    outpath="forest_plot.png",
):
    return compare_results(
        bi_long_posteriors, strand_posteriors, bi_wide_posteriors, outpath
    )


# %% Run
strand_post = r_to_py(ro.globalenv["strand_posteriors"])
compare_results(m.posteriors, strand_post, m2.posteriors)
# %%
```
