## Appendix 1, 2, 3, and 4 for "BayesForge: A Bayesian Inference library for Python, R, and Julia": test_strand.R

```
library(STRAND)
set.seed(1)
Mass_f = rnorm(50)
Age_f = rnorm(50)
individual = data.frame(Mass_f=Mass_f, Age_f=Age_f)
name_vec = paste0('Ind_', 1:50)
rownames(individual) = name_vec
outcome_mat = matrix(rbinom(2500, 1, 0.3), 50, 50)
rownames(outcome_mat) = colnames(outcome_mat) = name_vec
groups_f = data.frame(Merica=as.factor(sample(1:3, 50, replace=TRUE)))
rownames(groups_f) = name_vec
model_dat = make_strand_data(
  outcome=list(Association = outcome_mat),  
  block_covariates=groups_f, 
  individual_covariates=individual, 
  outcome_mode='bernoulli',
  link_mode='logit'
)
print(dim(model_dat$focal_individual_predictors))
print(head(model_dat$focal_individual_predictors))
```
