## Appendix 1, 2, 3, and 4 for "BayesForge: A Bayesian Inference library for Python, R, and Julia": brms_model.stan

```
// generated with brms 2.23.0
functions {
}
data {
  int<lower=1> N;  // total number of observations
  vector[N] Y;  // response variable
  int<lower=1> K;  // number of population-level effects
  matrix[N, K] X;  // population-level design matrix
  int<lower=1> Kc;  // number of population-level effects after centering
  // data for group-level effects of ID 1
  int<lower=1> N_1;  // number of grouping levels
  int<lower=1> M_1;  // number of coefficients per level
  array[N] int<lower=1> J_1;  // grouping indicator per observation
  matrix[N_1, N_1] Lcov_1;  // cholesky factor of known covariance matrix
  // group-level predictor values
  vector[N] Z_1_1;
  int prior_only;  // should the likelihood be ignored?
}
transformed data {
  matrix[N, Kc] Xc;  // centered version of X without an intercept
  vector[Kc] means_X;  // column means of X before centering
  for (i in 2:K) {
    means_X[i - 1] = mean(X[, i]);
    Xc[, i - 1] = X[, i] - means_X[i - 1];
  }
}
parameters {
  vector[Kc] b;  // regression coefficients
  real Intercept;  // temporary intercept for centered predictors
  real<lower=0> sigma;  // dispersion parameter
  vector<lower=0>[M_1] sd_1;  // group-level standard deviations
  array[M_1] vector[N_1] z_1;  // standardized group-level effects
}
transformed parameters {
  vector[N_1] r_1_1;  // actual group-level effects
  // prior contributions to the log posterior
  real lprior = 0;
  r_1_1 = (sd_1[1] * (Lcov_1 * z_1[1]));
  lprior += normal_lpdf(b | 0, 10);
  lprior += normal_lpdf(Intercept | 0, 50);
  lprior += student_t_lpdf(sigma | 3, 0, 20)
    - 1 * student_t_lccdf(0 | 3, 0, 20);
  lprior += student_t_lpdf(sd_1 | 3, 0, 20)
    - 1 * student_t_lccdf(0 | 3, 0, 20);
}
model {
  // likelihood including constants
  if (!prior_only) {
    // initialize linear predictor term
    vector[N] mu = rep_vector(0.0, N);
    mu += Intercept;
    for (n in 1:N) {
      // add more terms to the linear predictor
      mu[n] += r_1_1[J_1[n]] * Z_1_1[n];
    }
    target += normal_id_glm_lpdf(Y | Xc, mu, b, sigma);
  }
  // priors including constants
  target += lprior;
  target += std_normal_lpdf(z_1[1]);
}
generated quantities {
  // actual population-level intercept
  real b_Intercept = Intercept - dot_product(means_X, b);
}
```
