## Appendix 1, 2, 3, and 4 for "BayesForge: A Bayesian Inference library for Python, R, and Julia": README.md

# Phylo Example — PGLMM: BayesForge vs brms (Box 2)

Phylogenetic generalized linear mixed model (PGLMM) over 100 mammalian
posterior trees. Regresses `log(BodyLength_mm) ~ log(BodyMass_g)` with a
phylogenetic random effect whose covariance is the raw VCV matrix from the
tree (matches `ape::vcv.phylo` default).

Priors: Intercept ~ Normal(0, 50), b ~ Normal(0, 10),
sd & sigma ~ half-student_t(3, 0, 20). Non-centered parameterization in
BayesForge.

## Install

Python side:
```
pip install -r requirements.txt
```

R side (not pip-installable): R with `brms` and `ape` installed, and
`Rscript` on `PATH` — `scripts/run_comparison.py` shells out to
`scripts/brms_baseline.R` via `subprocess.run(["Rscript", ...])` and hard
`sys.exit`s if that fails.

**Not independently re-verified end-to-end for this README update** — no R
install was available in the environment these examples were last tested
from, so `run_comparison.py`'s brms step and `validate_single_tree.py`
couldn't be executed here. `scripts/validate_single_tree.py`'s import was
updated from the old `from BI import bi` to `from BayesForge import bf` to
match `run_comparison.py` (already on BayesForge), but that specific change
is untested.

## Repository layout

```
Phylo/
├── data/                      # Input data (tracked)
│   ├── pruned_traits.csv      # 50-species trait table
│   └── mammal_100_trees.nexus # 100 pruned posterior trees
├── scripts/                   # All runnable scripts
│   ├── run_comparison.py      # Master pipeline (steps 1–4)
│   ├── brms_baseline.R        # Called internally by run_comparison.py
│   ├── validate_single_tree.py# Single-tree BI vs brms sanity check
│   ├── acquire_data.py        # Download traits → pruned_traits.csv
│   └── extract_mammal_trees.py# Stream + prune 100 trees from Upham zip
└── output/                    # Generated files (not tracked)
    ├── bi_posteriors.csv
    ├── brms_posteriors.csv
    ├── comparison_table.csv
    ├── comparison_results.txt
    ├── posterior_densities.png
    ├── validation_single_tree.csv
    └── validation_single_tree.png
```

## Data sources

1. **Traits** — TetrapodTraits v1.0.0 (Zenodo `10.5281/zenodo.10530618`).
   Filtered to Mammalia, dropped rows missing `BodyMass_g`, `BodyLength_mm`,
   `Scientific.Name`. First 50 species → `data/pruned_traits.csv`.
   Reproduced by `scripts/acquire_data.py`.

2. **Trees** — Upham et al. (2019) mammal credible tree set (Dryad
   `Data_S7_Mammalia_credibleTreeSets_tipDR.zip`,
   `MamPhy_fullPosterior_BDvr_Completed_5911sp`). 100 trees streamed and
   pruned to the 50 target species → `data/mammal_100_trees.nexus`.
   Produced by `scripts/extract_mammal_trees.py` (RAM-safe streaming pruner;
   requires the Dryad zip at
   `Mamal tree/Data_S7_Mammalia_credibleTreeSets_tipDR.zip`).

## Usage

```bash
# One-off data acquisition (only needed once):
python scripts/acquire_data.py

# Full benchmark — BayesForge + brms + comparison table + density plot:
python scripts/run_comparison.py
# Outputs written to output/

# Single-tree validation — confirms BayesForge and brms recover identical posteriors:
python scripts/validate_single_tree.py
# Outputs written to output/
```

`scripts/run_comparison.py` calls `Rscript scripts/brms_baseline.R` internally.
Requires: brms, ape (R); BayesForge, jax, dendropy, pandas (Python — see
`requirements.txt`).

## VCV note

`dendropy`'s `distance_from_root()` includes the root edge length, while
`ape::vcv.phylo` ignores it. The Python code subtracts
`tree.seed_node.edge.length` from all distances so both frameworks use an
identical VCV matrix (max absolute diff < 1e-13).
