## Appendix 1, 2, 3, and 4 for "BayesForge: A Bayesian Inference library for Python, R, and Julia": requirements.txt

# Python side only. R side (brms, ape) is separate -- see README.md, not
# pip-installable. NOT independently verified end-to-end in this repo (no R
# available in the environment these examples were last tested from) -- pins
# below match Voice/SRM's verified BayesForge stack; report back if anything
# here needs adjusting once actually run against brms_baseline.R.

BayesForge>=0.0.56
jax>=0.10.0
jaxlib>=0.10.0
numpyro>=0.21.0
arviz>=1.2.0
numpy>=2.2.0
pandas
matplotlib
seaborn
scipy
dendropy
requests
