## Appendix 1, 2, 3, and 4 for "BayesForge: A Bayesian Inference library for Python, R, and Julia": acquire_data.py

```
import os
import requests
import pandas as pd
import zipfile
import shutil

os.makedirs('data', exist_ok=True)
print("Starting real-world mammal data acquisition...")

# 1. Download TetrapodTraits (already exists or download if missing)
traits_file = "data/TetrapodTraits.csv"
if not os.path.exists(traits_file):
    print("Downloading TetrapodTraits...")
    traits_url = "https://zenodo.org/records/10530618/files/TetrapodTraits_1.0.0.csv?download=1"
    r = requests.get(traits_url)
    with open(traits_file, "wb") as f:
        f.write(r.content)

print("Processing traits...")
df = pd.read_csv(traits_file)
if 'Class' in df.columns:
    df = df[df['Class'] == 'Mammalia']
df = df.dropna(subset=['BodyMass_g', 'BodyLength_mm', 'Scientific.Name'])
df_benchmark = df.head(50).copy()
df_benchmark['phylo_name'] = df_benchmark['Scientific.Name'].str.replace(' ', '_')
df_benchmark.to_csv("data/pruned_traits.csv", index=False)
print(f"Saved pruned traits for {len(df_benchmark)} mammals.")

# 2. Download Upham et al. (2019) Mammal Tree Sets from Dryad (Data_S7)
# Link: https://datadryad.org/downloads/file_stream/188729
trees_zip = "data/Upham_Trees.zip"
nexus_target = "data/mam_1000_subset.nexus"

if not os.path.exists(nexus_target):
    if not os.path.exists(trees_zip):
        print("Downloading Upham et al. (2019) credible tree sets (3.7 GB)...")
        # I'll use a stream to avoid loading the whole thing into RAM
        trees_url = "https://datadryad.org/downloads/file_stream/188729"
        try:
            with requests.get(trees_url, stream=True) as r:
                r.raise_for_status()
                with open(trees_zip, 'wb') as f:
                    shutil.copyfileobj(r.raw, f)
            print("Download complete.")
        except Exception as e:
            print(f"Error downloading: {e}")
            # Fallback to simulation if download fails in this environment
            print("Falling back to simulation...")
            import dendropy
            from dendropy.simulate import treesim
            taxa = dendropy.TaxonNamespace(df_benchmark['phylo_name'])
            trees = dendropy.TreeList(taxon_namespace=taxa)
            for i in range(1000):
                t = treesim.pure_kingman_tree(taxon_namespace=taxa, pop_size=1)
                trees.append(t)
            trees.write(path=nexus_target, schema="nexus")
            print("Simulated 1000 benchmark trees.")
            exit(0)

    print("Extracting 1000 trees from Dryad zip...")
    import dendropy
    try:
        # The zip contains several large NEXUS files. 
        # We look for a file like 'MamPhy_fullPosterior_BDexp_completed_5911sp_v1.nexus'
        with zipfile.ZipFile(trees_zip, 'r') as zip_ref:
            # List files to find the right one
            names = zip_ref.namelist()
            print(f"Files in zip: {names[:5]}...")
            # We want the DNA-only or completed set. Let's look for 'completed'
            target_name = [n for n in names if 'completed' in n and n.endswith('.nexus')][0]
            print(f"Extracting {target_name}...")
            with zip_ref.open(target_name) as f:
                # Instead of reading the whole 10,000 trees, we'll try to read only 1000
                # DendroPy can take a file-like object
                trees = dendropy.TreeList.get(file_obj=f, schema="nexus", tree_offset=0)
                # Subset to 1000 trees
                trees = trees[:1000]
                
                # Taxon matching logic
                # The Dryad trees have names like 'Genus_species_ID'
                tree_taxa = [t.label for t in trees.taxon_namespace]
                mapping = {}
                for name in df_benchmark['phylo_name']:
                    # Try exact match or substring
                    matches = [t for t in tree_taxa if name in t]
                    if matches:
                        mapping[matches[0]] = name
                
                print(f"Matched {len(mapping)}/50 taxa.")
                
                # Prune and Rename
                for tree in trees:
                    tree.retain_taxa_with_labels(mapping.keys())
                    for taxon in tree.taxon_namespace:
                        if taxon.label in mapping:
                            taxon.label = mapping[taxon.label]
                
                trees.write(path=nexus_target, schema="nexus")
                print(f"Successfully saved 1000 real mammal trees to {nexus_target}")
    except Exception as e:
        print(f"Error during extraction/pruning: {e}")
        # Note: In some environments, zip extraction of 3GB files might be limited.

else:
    print(f"Target nexus {nexus_target} already exists.")
```
