## Appendix 1, 2, 3, and 4 for "BayesForge: A Bayesian Inference library for Python, R, and Julia": brms_baseline.R

```
# brms_baseline.R — PGLMM over 100 posterior trees
# Mirrors BI/Test/Phylogenic analysis/Model_1_Simple:
#   - raw vcv.phylo (covariance, NOT corr=TRUE)
#   - priors: Intercept ~ Normal(0,50), b ~ Normal(0,10),
#             sd & sigma ~ student_t(3, 0, 20)
#   - raw traits (no standardisation)
library(brms)
library(ape)

cat("Starting brms baseline execution...\n")

traits_file <- "data/pruned_traits.csv"
trees_file <- "data/mammal_100_trees.nexus"
out_file  <- "output/brms_posteriors.csv"
diag_file <- "output/brms_diagnostics.csv"
dir.create("output", showWarnings = FALSE)

if (!file.exists(traits_file) || !file.exists(trees_file)) {
    stop("Data files not found.")
}

dat <- read.csv(traits_file, stringsAsFactors = FALSE)
dat$phylo <- gsub(" ", "_", as.character(dat$Scientific.Name))
# log-transform: raw scale spans 3+ orders of magnitude (24 g – 64 kg).
dat$y <- log(as.numeric(dat$BodyLength_mm))
dat$x <- log(as.numeric(dat$BodyMass_g))
trees <- read.nexus(trees_file)
num_trees <- min(length(trees), 100L)

# ── compile once on tree 1 ────────────────────────────────────────────────────
cat("Compiling model on tree 1 (one-time cost)...\n")

prep_tree <- function(tr, dat) {
    tr$tip.label <- gsub("'", "", tr$tip.label)
    A <- ape::vcv.phylo(tr) # RAW VCV (no corr=TRUE)
    keep <- intersect(dat$phylo, rownames(A))
    list(A = A[keep, keep], dat = dat[dat$phylo %in% keep, ])
}

p1 <- prep_tree(trees[[1]], dat)
fit_base <- brm(
    y ~ x + (1 | gr(phylo, cov = A)),
    data = p1$dat,
    data2 = list(A = p1$A),
    family = gaussian(),
    prior = c(
        prior(normal(0, 10), class = "b"),
        prior(normal(0, 50), class = "Intercept"),
        prior(student_t(3, 0, 20), class = "sd"),
        prior(student_t(3, 0, 20), class = "sigma")
    ),
    chains = 2, cores = 1, iter = 1100, warmup = 1000,
    seed = 42, refresh = 0, silent = 2
)
rm(p1)
gc()

# ── save raw Stan code so priors can be audited ───────────────────────────────
writeLines(stancode(fit_base), "output/brms_model.stan")
cat("Stan code written -> output/brms_model.stan\n")

# ── stream results to CSV (no accumulation in RAM) ───────────────────────────
COLS <- c("b_Intercept", "b_x", "sigma", "sd_phylo__Intercept")
DIAG_COLS <- c("parameter", "Bulk_ESS", "Tail_ESS", "Rhat", "tree_index", "model")

if (file.exists(out_file)) file.remove(out_file)
if (file.exists(diag_file)) file.remove(diag_file)
con <- file(out_file, open = "w")
con_diag <- file(diag_file, open = "w")
writeLines(paste(c(COLS, "tree_index"), collapse = ","), con)
writeLines(paste(DIAG_COLS, collapse = ","), con_diag)

cat("Iterating over trees...\n")

for (i in seq_len(num_trees)) {
    cat(sprintf("  Tree %d/%d\n", i, num_trees))

    pi <- prep_tree(trees[[i]], dat)

    fit_i <- if (i == 1L) {
        fit_base
    } else {
        update(fit_base,
            newdata   = pi$dat,
            data2     = list(A = pi$A),
            recompile = FALSE,
            refresh   = 0,
            silent    = 2
        )
    }
    rm(pi)
    gc()

    post <- as_draws_df(fit_i)[, COLS]
    post$tree_index <- i
    write.table(post, con,
        sep = ",", row.names = FALSE, col.names = FALSE,
        quote = FALSE, append = TRUE
    )

    # ESS + Rhat from brms summary
    smry <- summary(fit_i)
    fx <- smry$fixed[, c("Bulk_ESS", "Tail_ESS", "Rhat"), drop = FALSE]
    fx$parameter <- rownames(fx)
    fx$tree_index <- i
    fx$model <- "brms"
    rx <- smry$random$phylo[, c("Bulk_ESS", "Tail_ESS", "Rhat"), drop = FALSE]
    rx$parameter <- paste0("sd_phylo__", rownames(rx))
    rx$tree_index <- i
    rx$model <- "brms"
    diag_i <- rbind(fx[, DIAG_COLS], rx[, DIAG_COLS])
    write.table(diag_i, con_diag,
        sep = ",", row.names = FALSE, col.names = FALSE,
        quote = FALSE, append = TRUE
    )

    rm(post, smry, fx, rx, diag_i)
    if (i > 1L) rm(fit_i)
    if (i %% 10L == 0L) gc()
}

close(con)
close(con_diag)
cat("Done. Results    ->", out_file, "\n")
cat("Done. Diagnostics ->", diag_file, "\n")
```
