## Appendix 1, 2, 3, and 4 for "BayesForge: A Bayesian Inference library for Python, R, and Julia": extract_mammal_trees.py

```
#!/usr/bin/env python3.13
"""
extract_mammal_trees.py
=======================
Stream N trees from the Upham Mammalia zip, prune to trait species, write nexus.

Strategy (RAM-limited, fast):
  - Never hold more than BATCH_SIZE pruned strings in RAM before flushing.
  - Uses a zero-dependency recursive newick pruner (~15 ms/tree).
  - Peak RAM ≈ 1 raw tree line (~100 KB) + translate dict (~2 MB).

Source file: MamPhy_fullPosterior_BDvr_Completed_5911sp (all 50 target species present).

Usage:
    python3.13 scripts/extract_mammal_trees.py [--n 100] [--batch 10]
"""

import argparse
import io
import os
import re
import zipfile
import gc
import time

import pandas as pd

SCRIPT_DIR    = os.path.dirname(os.path.abspath(__file__))
BASE_DIR      = os.path.dirname(SCRIPT_DIR)
ZIP_PATH      = os.path.join(BASE_DIR, "Mamal tree",
                             "Data_S7_Mammalia_credibleTreeSets_tipDR.zip")
INTERNAL_FILE = ("Data_S7_Mammalia_credibleTreeSets_tipDR/"
                 "MamPhy_fullPosterior_BDvr_Completed_5911sp_topoCons_NDexp_"
                 "all10k_v2_nexus.trees")
TRAITS_PATH   = os.path.join(BASE_DIR, "data", "pruned_traits.csv")
OUTPUT_PATH   = os.path.join(BASE_DIR, "data", "mammal_100_trees.nexus")

# ── fast newick pruner ────────────────────────────────────────────────────────

_BEAST_ANNOT = re.compile(r"^\s*\[&[A-Za-z]+\]\s*")
_LEAF_PAT    = re.compile(r"\b(\d+):")


def prune_newick(s: str, keep: set) -> str | None:
    """
    Prune newick string to only leaves whose labels are in `keep`.
    Handles: BEAST [&R] annotations, unifurcations (merges branch lengths).
    Returns pruned newick with trailing ';', or None if no target species.
    """
    s = _BEAST_ANNOT.sub("", s).rstrip().rstrip(";")
    pos = [0]
    n = len(s)

    def skip_comment():
        while pos[0] < n and s[pos[0]] == "[":
            while pos[0] < n and s[pos[0]] != "]":
                pos[0] += 1
            pos[0] += 1

    def read_bl():
        skip_comment()
        if pos[0] < n and s[pos[0]] == ":":
            pos[0] += 1
            start = pos[0]
            while pos[0] < n and s[pos[0]] not in ",);[":
                pos[0] += 1
            try:
                return float(s[start : pos[0]])
            except ValueError:
                return None
        return None

    def node():
        """Returns (newick_str_without_own_bl, bl_to_parent) or (None, bl)."""
        if pos[0] >= n:
            return None, None

        if s[pos[0]] == "(":
            pos[0] += 1
            children = []
            while True:
                cnwk, cbl = node()
                if cnwk is not None:
                    children.append((cnwk, cbl))
                while pos[0] < n and s[pos[0]] == " ":
                    pos[0] += 1
                if pos[0] < n and s[pos[0]] == ",":
                    pos[0] += 1
                elif pos[0] < n and s[pos[0]] == ")":
                    pos[0] += 1
                    break
                else:
                    break
            while pos[0] < n and s[pos[0]] not in ":,);[":
                pos[0] += 1
            bl = read_bl()

            if not children:
                return None, bl

            if len(children) == 1:
                cnwk, cbl = children[0]
                new_bl = (
                    (cbl or 0.0) + (bl or 0.0)
                    if (cbl is not None or bl is not None)
                    else None
                )
                return cnwk, new_bl

            parts = [
                f'{c}:{b:.10g}' if b is not None else c
                for c, b in children
            ]
            return f'({",".join(parts)})', bl

        else:
            start = pos[0]
            while pos[0] < n and s[pos[0]] not in ":,);[":
                pos[0] += 1
            label = s[start : pos[0]]
            bl = read_bl()
            return (label, bl) if label in keep else (None, bl)

    nwk, bl = node()
    if nwk is None:
        return None
    bl_s = f":{bl:.10g}" if bl is not None else ""
    return f"{nwk}{bl_s};"


# ── helpers ───────────────────────────────────────────────────────────────────

def parse_translate_block(f) -> dict:
    """Read lines until closing ';'. Returns {number_str: full_taxon_name}."""
    translate = {}
    for raw in f:
        line = raw.decode("utf-8") if isinstance(raw, bytes) else raw
        stripped = line.strip().rstrip(",")
        if stripped == ";":
            break
        parts = stripped.split(None, 1)
        if len(parts) == 2:
            translate[parts[0]] = parts[1].strip("'")
    return translate


def build_num_to_target(translate: dict, target_set: set) -> dict:
    """Map translate numbers → clean target name (only for target species)."""
    num_to_target = {}
    for num, full_name in translate.items():
        for target in target_set:
            if full_name == target or full_name.startswith(target + "_"):
                num_to_target[num] = target
                break
    return num_to_target


# ── main ──────────────────────────────────────────────────────────────────────

def main(num_trees: int = 100, batch_size: int = 10) -> None:
    t0 = time.time()

    df          = pd.read_csv(TRAITS_PATH)
    target_set  = set(df["Scientific.Name"].str.replace(" ", "_").tolist())
    target_list = sorted(target_set)
    print(f"  Target species: {len(target_set)}")

    os.makedirs(os.path.dirname(OUTPUT_PATH), exist_ok=True)

    with zipfile.ZipFile(ZIP_PATH, "r") as z, \
         z.open(INTERNAL_FILE) as raw_bytes, \
         open(OUTPUT_PATH, "w") as fout:

        f = io.TextIOWrapper(raw_bytes, encoding="utf-8", errors="replace")

        # 1. Seek to TRANSLATE block
        for raw_line in f:
            if "TRANSLATE" in raw_line.upper():
                break

        translate     = parse_translate_block(f)
        num_to_target = build_num_to_target(translate, target_set)
        matched       = len(set(num_to_target.values()))
        print(f"  Translate: {len(translate)} taxa, matched {matched}/{len(target_set)} targets")

        if matched < len(target_set):
            missing = target_set - set(num_to_target.values())
            print(f"  WARNING: missing species: {sorted(missing)[:5]}...")

        # 2. Pre-compile rename substitution (target numbers only).
        #    Quote species names so NEXUS readers preserve underscores.
        def rename_targets(newick: str) -> str:
            def r(m):
                nn = m.group(1)
                if nn in num_to_target:
                    return f"'{num_to_target[nn]}':"
                return nn + ":"
            return _LEAF_PAT.sub(r, newick)

        # Pruner `keep` must match the quoted form used in the newick
        target_set_quoted = {f"'{sp}'" for sp in target_set}

        # 3. Write nexus header (quoted names → underscores preserved)
        fout.write("#NEXUS\n\nBEGIN TAXA;\n")
        fout.write(f"    DIMENSIONS NTAX={len(target_list)};\n")
        fout.write("    TAXLABELS\n")
        for sp in target_list:
            fout.write(f"        '{sp}'\n")
        fout.write("    ;\nEND;\n\nBEGIN TREES;\n")

        # 4. Stream trees
        trees_done = 0
        skipped    = 0

        for raw_line in f:
            if trees_done >= num_trees:
                break

            line = raw_line.strip()
            if not line or line.upper().startswith("END"):
                continue
            if not line.upper().startswith("TREE"):
                continue

            eq     = line.index("=")
            newick = line[eq + 1 :].strip()

            try:
                named  = rename_targets(newick)
                pruned = prune_newick(named, target_set_quoted)

                if pruned is None:
                    skipped += 1
                    continue

                # Quick sanity: count matched leaves
                n_leaves = sum(1 for sp in target_set_quoted if sp in pruned)
                if n_leaves < matched:
                    skipped += 1
                    continue

                fout.write(f"    TREE {trees_done + 1} = {pruned}\n")
                trees_done += 1

                if trees_done % batch_size == 0:
                    elapsed = time.time() - t0
                    rate    = trees_done / elapsed
                    eta     = (num_trees - trees_done) / rate if rate > 0 else 0
                    print(
                        f"  [{trees_done:>4}/{num_trees}] "
                        f"{elapsed:.1f}s elapsed, ETA {eta:.0f}s"
                    )
                    fout.flush()
                    gc.collect()

            except Exception as exc:
                skipped += 1
                if skipped <= 5:
                    print(f"  Warning tree {trees_done + skipped}: {exc}")

        fout.write("END;\n")

    elapsed = time.time() - t0
    print(f"\n  Done: {trees_done} trees → {OUTPUT_PATH}")
    print(
        f"  Skipped: {skipped}  |  Total: {elapsed:.1f}s  "
        f"({elapsed / trees_done:.2f}s/tree)"
    )


if __name__ == "__main__":
    p = argparse.ArgumentParser()
    p.add_argument("--n",     type=int, default=100, help="Trees to extract")
    p.add_argument("--batch", type=int, default=10,  help="Flush/GC every N trees")
    args = p.parse_args()
    main(num_trees=args.n, batch_size=args.batch)
```
