## Appendix 1, 2, 3, and 4 for "BayesForge: A Bayesian Inference library for Python, R, and Julia": plot_diagnostics.py

```
#!/usr/bin/env python3
"""
plot_diagnostics.py — MCMC Diagnostics: ESS, R-hat, Efficiency
===============================================================
Reads output/bi_posteriors.csv and output/brms_diagnostics.csv.
Computes BI ESS/R-hat via ArviZ (assumes chains are stacked:
  first N_SAMPLES rows per tree = chain 0, next N_SAMPLES = chain 1).
Produces:
  output/diagnostics_table.png   — ESS / R-hat heatmap (style of Image #6)
  output/efficiency_table.png    — ESS/second efficiency comparison

Requires timing info from output/comparison_results.txt or falls back to
BI_TIME_S / BRMS_TIME_S constants set below.
"""

import os, re, sys
import numpy as np
import pandas as pd
import matplotlib
matplotlib.use("Agg")
import matplotlib.pyplot as plt
import matplotlib.colors as mcolors
import arviz as az

# ── paths ─────────────────────────────────────────────────────────────────────
SCRIPT_DIR = os.path.dirname(os.path.abspath(__file__))
ROOT_DIR   = os.path.dirname(SCRIPT_DIR)
OUT_DIR    = os.path.join(ROOT_DIR, "output")

BI_POST_CSV  = os.path.join(OUT_DIR, "bi_posteriors.csv")
BRMS_DIAG_CSV= os.path.join(OUT_DIR, "brms_diagnostics.csv")
TIMING_TXT   = os.path.join(OUT_DIR, "comparison_results.txt")
OUT_DIAG_PNG = os.path.join(OUT_DIR, "diagnostics_table.png")
OUT_EFF_PNG  = os.path.join(OUT_DIR, "efficiency_table.png")

for f in [BI_POST_CSV, BRMS_DIAG_CSV]:
    if not os.path.exists(f):
        sys.exit(f"Missing: {f} — run scripts/run_comparison.py first.")

# ── MCMC settings (must match run_comparison.py) ──────────────────────────────
N_CHAINS  = 2
N_SAMPLES = 500   # post-warmup samples per chain in BI

# ── parse timing from comparison_results.txt ─────────────────────────────────
bi_time_s   = None
brms_time_s = None
if os.path.exists(TIMING_TXT):
    txt = open(TIMING_TXT).read()
    m = re.search(r"brms:\s*([\d.]+)\s*s", txt)
    if m: brms_time_s = float(m.group(1))
    m = re.search(r"BI:\s*([\d.]+)\s*s", txt)
    if m: bi_time_s   = float(m.group(1))
if bi_time_s is None or brms_time_s is None:
    sys.exit("Cannot parse timing from comparison_results.txt. Check file format.")
print(f"Timing — BI: {bi_time_s:.2f}s   brms: {brms_time_s:.2f}s")

# ── parameter label maps ──────────────────────────────────────────────────────
# brms parameter names → display label
BRMS_PARAM_MAP = {
    "Intercept":               "b_Intercept",
    "x":                       "b_x",
    "sd_phylo__sd(Intercept)": "sd_phylo__Intercept",
    "sigma":                   "sigma",
}
# BI column names in bi_posteriors.csv
BI_PARAMS = ["b_Intercept", "b_x", "sigma", "sd_phylo__Intercept"]

DISPLAY = {
    "b_Intercept":         "Intercept",
    "b_x":                 "Slope (β)",
    "sigma":               "Sigma (residual)",
    "sd_phylo__Intercept": "SD phylo",
}

# ── brms diagnostics ──────────────────────────────────────────────────────────
brms_raw = pd.read_csv(BRMS_DIAG_CSV)
brms_raw["param_key"] = brms_raw["parameter"].map(BRMS_PARAM_MAP)
brms_raw = brms_raw.dropna(subset=["param_key"])

brms_diag = (brms_raw.groupby("param_key")
             .agg(mean_ESS=("Bulk_ESS", "mean"),
                  max_Rhat=("Rhat",     "max"))
             .reset_index())
print("\nbrms diagnostics (mean ESS, max R-hat across 100 trees):")
print(brms_diag.to_string(index=False))

# ── BI ESS / R-hat via ArviZ ──────────────────────────────────────────────────
print("\nComputing BI ESS/R-hat from posterior samples ...")
bi_post = pd.read_csv(BI_POST_CSV)
trees   = sorted(bi_post["tree_index"].unique())

bi_ess_rows = []
for t in trees:
    sub = bi_post[bi_post["tree_index"] == t].reset_index(drop=True)
    # samples stacked: rows 0..N_SAMPLES-1 = chain 0, N_SAMPLES..2*N_SAMPLES-1 = chain 1
    n_total = len(sub)
    n_per_chain = n_total // N_CHAINS
    for p in BI_PARAMS:
        vals = sub[p].values  # (N_CHAINS * n_per_chain,)
        chains = vals.reshape(N_CHAINS, n_per_chain)  # (chains, draws)
        data_dict = {"x": chains}
        ess  = float(az.ess(data_dict)["x"].item())
        rhat = float(az.rhat(data_dict)["x"].item())
        bi_ess_rows.append({"param_key": p, "tree_index": t, "Bulk_ESS": ess, "Rhat": rhat})

bi_ess_df = pd.DataFrame(bi_ess_rows)
bi_diag = (bi_ess_df.groupby("param_key")
           .agg(mean_ESS=("Bulk_ESS", "mean"),
                max_Rhat=("Rhat",     "max"))
           .reset_index())
print("\nBI diagnostics (mean ESS, max R-hat across 100 trees):")
print(bi_diag.to_string(index=False))

# ── build combined table ──────────────────────────────────────────────────────
# rows = display params, cols = brms | BI
all_params = [p for p in BI_PARAMS if p in brms_diag["param_key"].values or p in bi_diag["param_key"].values]

ess_df  = pd.DataFrame(index=[DISPLAY[p] for p in all_params], columns=["brms", "BI hierarchical"])
rhat_df = pd.DataFrame(index=[DISPLAY[p] for p in all_params], columns=["brms", "BI hierarchical"])

for p in all_params:
    label = DISPLAY[p]
    brms_row = brms_diag[brms_diag["param_key"] == p]
    bi_row   = bi_diag[bi_diag["param_key"] == p]
    ess_df.loc[label, "brms"]              = round(brms_row["mean_ESS"].values[0], 0) if len(brms_row) else np.nan
    ess_df.loc[label, "BI hierarchical"]   = round(bi_row["mean_ESS"].values[0],   0) if len(bi_row)   else np.nan
    rhat_df.loc[label, "brms"]             = round(brms_row["max_Rhat"].values[0],  3) if len(brms_row) else np.nan
    rhat_df.loc[label, "BI hierarchical"]  = round(bi_row["max_Rhat"].values[0],    3) if len(bi_row)   else np.nan

ess_df  = ess_df.astype(float)
rhat_df = rhat_df.astype(float)

# ── efficiency metric: ESS / total wall-clock time ───────────────────────────
eff_df = pd.DataFrame(index=ess_df.index, columns=["brms", "BI hierarchical"])
eff_df["brms"]            = (ess_df["brms"]            / brms_time_s).round(2)
eff_df["BI hierarchical"] = (ess_df["BI hierarchical"] / bi_time_s  ).round(2)
eff_df = eff_df.astype(float)

print("\nEfficiency (ESS / second):")
print(eff_df.to_string())

# ══════════════════════════════════════════════════════════════════════════════
# PLOT 1 — ESS / R-hat heatmap (matches Image #6 style)
# ══════════════════════════════════════════════════════════════════════════════
def _heatmap(ax, data, fmt, cmap, vmin, vmax, title, cbar_label):
    n_rows, n_cols = data.shape
    arr = data.values.astype(float)

    norm = mcolors.Normalize(vmin=vmin, vmax=vmax)
    im   = ax.imshow(arr, cmap=cmap, norm=norm, aspect="auto")

    # cell text
    for r in range(n_rows):
        for c in range(n_cols):
            v = arr[r, c]
            if np.isnan(v):
                continue
            text = fmt.format(v)
            brightness = im.cmap(norm(v))[:3]
            lum = 0.299*brightness[0] + 0.587*brightness[1] + 0.114*brightness[2]
            fc  = "white" if lum < 0.55 else "black"
            ax.text(c, r, text, ha="center", va="center", fontsize=11,
                    color=fc, fontweight="bold")

    ax.set_xticks(range(n_cols))
    ax.set_xticklabels(data.columns, fontsize=11)
    ax.set_yticks(range(n_rows))
    ax.set_yticklabels(data.index, fontsize=10)
    ax.tick_params(length=0)

    # white grid lines
    for r in range(n_rows + 1):
        ax.axhline(r - 0.5, color="white", lw=1.5)
    for c in range(n_cols + 1):
        ax.axvline(c - 0.5, color="white", lw=1.5)

    ax.set_title(title, fontsize=13, fontweight="bold", pad=10)
    cbar = plt.colorbar(im, ax=ax, fraction=0.04, pad=0.02)
    cbar.set_label(cbar_label, fontsize=9)
    return im


fig, (ax1, ax2) = plt.subplots(1, 2, figsize=(14, 0.85 * len(all_params) + 3))
fig.patch.set_facecolor("white")

_heatmap(ax1, ess_df,  "{:.0f}", "Greens",
         vmin=0, vmax=max(ess_df.max().max(), 500),
         title="Mean ESS\n(averaged across 100 trees)",
         cbar_label="ESS")

_heatmap(ax2, rhat_df, "{:.3f}", "RdYlGn_r",
         vmin=1.00, vmax=1.10,
         title="Max R-hat\n(worst across 100 trees)",
         cbar_label="R-hat")

fig.suptitle(
    "MCMC Diagnostics — ESS and R-hat\n"
    f"brms · BI independent PGLMM  "
    f"({N_CHAINS} chains × {N_SAMPLES} samples | 100 trees)",
    fontsize=13, fontweight="bold", y=1.03,
)
plt.tight_layout()
plt.savefig(OUT_DIAG_PNG, dpi=200, bbox_inches="tight")
print(f"\nSaved: {OUT_DIAG_PNG}")

# ══════════════════════════════════════════════════════════════════════════════
# PLOT 2 — Efficiency (ESS / second) bar chart
# ══════════════════════════════════════════════════════════════════════════════
fig2, axes2 = plt.subplots(1, 2, figsize=(14, 5))

CLR = {"brms": "#E07B54", "BI hierarchical": "#4D8FAC"}
x   = np.arange(len(all_params))
labels = [DISPLAY[p] for p in all_params]
width = 0.35

for ax, metric_df, title, ylabel in [
    (axes2[0], eff_df,  "Efficiency: ESS / Wall-Clock Second", "ESS per second"),
    (axes2[1], ess_df,  "Raw ESS (mean across 100 trees)",      "Mean ESS"),
]:
    for i, (col, color) in enumerate(CLR.items()):
        vals = metric_df[col].values
        bars = ax.bar(x + (i - 0.5) * width, vals, width,
                      label=col, color=color, alpha=0.85, edgecolor="white")
        for bar, v in zip(bars, vals):
            if not np.isnan(v):
                ax.text(bar.get_x() + bar.get_width()/2, bar.get_height() + 0.5,
                        f"{v:.1f}", ha="center", va="bottom", fontsize=8)
    ax.set_xticks(x)
    ax.set_xticklabels(labels, rotation=20, ha="right", fontsize=10)
    ax.set_ylabel(ylabel, fontsize=11)
    ax.set_title(title, fontsize=12, fontweight="bold")
    ax.legend(fontsize=10)
    ax.spines[["top", "right"]].set_visible(False)

# annotate speedup
axes2[0].text(0.98, 0.97,
    f"brms total: {brms_time_s:.1f}s\nBI total:   {bi_time_s:.1f}s\n"
    f"BI speedup: {brms_time_s/bi_time_s:.0f}×",
    transform=axes2[0].transAxes, ha="right", va="top", fontsize=9,
    bbox=dict(boxstyle="round,pad=0.4", facecolor="white", edgecolor="#ccc", alpha=0.9))

fig2.suptitle(
    "MCMC Efficiency: ESS / Total Wall-Clock Time\n"
    "Higher = more posterior information per second",
    fontsize=13, fontweight="bold",
)
plt.tight_layout()
plt.savefig(OUT_EFF_PNG, dpi=200, bbox_inches="tight")
print(f"Saved: {OUT_EFF_PNG}")
```
