## Appendix 1, 2, 3, and 4 for "BayesForge: A Bayesian Inference library for Python, R, and Julia": run_comparison.py

```
#!/usr/bin/env python3
"""
run_comparison.py  —  Phylogenetic PGLMM benchmark: BI vs brms (Box 2)
=======================================================================
Steps
  1. BI joint inference on 100 posterior trees (non-centered PGLMM)
  2. brms sequential loop over the same 100 trees
  3. Comparative table (CSV + terminal) with performance benchmark
  4. Density-overlay plot for all 4 parameters

Input  : data/pruned_traits.csv, data/mammal_100_trees.nexus
Output : output/  (bi_posteriors.csv, brms_posteriors.csv,
                   comparison_table.csv, comparison_results.txt,
                   posterior_densities.png)

Single-tree validation lives in validate_single_tree.py (same scripts/ folder).
"""

import os
import sys
import time
import subprocess

SCRIPT_DIR  = os.path.dirname(os.path.abspath(__file__))    # Phylo/scripts/
ROOT_DIR    = os.path.dirname(SCRIPT_DIR)                    # Phylo/
DATA_DIR    = os.path.join(ROOT_DIR, "data")
OUTPUT_DIR  = os.path.join(ROOT_DIR, "output")
BI_CSV      = os.path.join(OUTPUT_DIR, "bi_posteriors.csv")
BRMS_CSV    = os.path.join(OUTPUT_DIR, "brms_posteriors.csv")
TABLE_CSV   = os.path.join(OUTPUT_DIR, "comparison_table.csv")
TABLE_TXT   = os.path.join(OUTPUT_DIR, "comparison_results.txt")
PLOT_FILE   = os.path.join(OUTPUT_DIR, "posterior_densities.png")
NUM_TREES   = 100

os.makedirs(OUTPUT_DIR, exist_ok=True)
os.chdir(ROOT_DIR)

TREES_FILE = os.path.join(DATA_DIR, "mammal_100_trees.nexus")
EXTRACTOR  = os.path.join(SCRIPT_DIR, "extract_mammal_trees.py")


# ══════════════════════════════════════════════════════════════════════════════
# STEP 0 – Extract 100 trees from the Upham mammalia zip (streaming, RAM-safe)
# ══════════════════════════════════════════════════════════════════════════════
def _count_nexus_trees(path):
    count = 0
    with open(path) as fh:
        for ln in fh:
            if ln.strip().upper().startswith("TREE "):
                count += 1
    return count


need_extract = (
    not os.path.exists(TREES_FILE) or _count_nexus_trees(TREES_FILE) < NUM_TREES
)

if need_extract:
    print("\n" + "=" * 70)
    print(f"  STEP 0 | Extracting {NUM_TREES} trees from Mammal zip (streaming)")
    print("=" * 70)
    t0 = time.time()
    result = subprocess.run(
        [sys.executable, EXTRACTOR, "--n", str(NUM_TREES), "--batch", "10"],
        cwd=ROOT_DIR,
        capture_output=False,
        text=True,
    )
    if result.returncode != 0:
        sys.exit("Tree extraction failed.")
    print(f"  Extraction done in {time.time()-t0:.1f}s")
else:
    n = _count_nexus_trees(TREES_FILE)
    print(f"\n  STEP 0 | Using existing {n}-tree file ({TREES_FILE})")


# ══════════════════════════════════════════════════════════════════════════════
# STEP 1 – BI (JAX / NumPyro)
# ══════════════════════════════════════════════════════════════════════════════
print("\n" + "=" * 70)
print(f"  STEP 1 | Running BI inference on {NUM_TREES} trees (Box 2 MVN)")
print("=" * 70)

import time, pandas as pd, numpy as np
import jax.numpy as jnp
from BayesForge import bf

m = bf(platform="cpu")


def load_data(num_trees_to_load=NUM_TREES):
    import dendropy

    traits_file = os.path.join(DATA_DIR, "pruned_traits.csv")
    trees_file  = os.path.join(DATA_DIR, "mammal_100_trees.nexus")

    if not os.path.exists(traits_file) or not os.path.exists(trees_file):
        raise FileNotFoundError("Missing data — run scripts/acquire_data.py first.")

    df = pd.read_csv(traits_file)
    df["phylo_name"] = df["Scientific.Name"].str.replace(" ", "_")
    df = df.sort_values("phylo_name").reset_index(drop=True)
    df["y"] = np.log(df["BodyLength_mm"].astype(float))
    df["x"] = np.log(df["BodyMass_g"].astype(float))

    trees = dendropy.TreeList.get(path=trees_file, schema="nexus")
    trees = trees[:num_trees_to_load]

    n_sp = len(df)
    L_tensor = np.zeros((num_trees_to_load, n_sp, n_sp))

    print(f"  Computing {num_trees_to_load} raw VCV Cholesky factors ...")
    for i, tree in enumerate(trees):
        for t in tree.taxon_namespace:
            if t.label and t.label.startswith("'") and t.label.endswith("'"):
                t.label = t.label[1:-1]

        # dendropy distance_from_root() includes the root edge; ape::vcv.phylo ignores it.
        root_offset = tree.seed_node.edge.length or 0.0

        pdm  = tree.phylogenetic_distance_matrix()
        taxa = [t for t in tree.taxon_namespace if t.label in df["phylo_name"].values]
        taxa = sorted(taxa, key=lambda t: t.label)
        taxa_nodes = {
            t: tree.find_node(filter_fn=lambda n, t=t: n.taxon == t) for t in taxa
        }

        A_raw = np.zeros((n_sp, n_sp))
        for idx1, t1 in enumerate(taxa):
            node1 = taxa_nodes[t1]
            for idx2, t2 in enumerate(taxa):
                if idx1 == idx2:
                    A_raw[idx1, idx2] = node1.distance_from_root() - root_offset
                elif idx1 < idx2:
                    mrca = pdm.mrca(t1, t2)
                    dist = (
                        mrca.distance_from_root() - root_offset
                        if hasattr(mrca, "distance_from_root")
                        else 0.0
                    )
                    A_raw[idx1, idx2] = dist
                    A_raw[idx2, idx1] = dist

        L_tensor[i] = np.linalg.cholesky(A_raw + 1e-8 * np.eye(n_sp))

    df_sorted = df.sort_values("phylo_name").reset_index(drop=True)
    return (
        df_sorted["x"].values,
        df_sorted["y"].values,
        L_tensor,
        df_sorted["phylo_name"].tolist(),
    )


print(f"  Loading traits + pre-computing {NUM_TREES} trees ...")
X_np, Y_np, L_np, sp_names = load_data(NUM_TREES)
N_SP = len(sp_names)

mean_x   = float(X_np.mean())
X_centred = X_np - mean_x

X = jnp.array(X_centred, dtype=jnp.float64)
Y = jnp.array(Y_np,       dtype=jnp.float64)
L = jnp.array(L_np,       dtype=jnp.float64)
phylo_idx = jnp.arange(N_SP, dtype=jnp.int32)

m.data_on_model = dict(X=X, Y=Y, L=L, phylo_idx=phylo_idx)


def model(X, Y, L, phylo_idx):
    T = L.shape[0]
    N = X.shape[0]

    intercept  = m.dist.normal(0, 50, name="Intercept",  shape=(T,))
    b_cofactor = m.dist.normal(0, 10, name="b_cofactor", shape=(T,))
    sd_phylo   = m.dist.left_truncated_distribution(
        m.dist.student_t(3, 0, 20, create_obj=True), low=0.0, name="sd_phylo", shape=(T,)
    )
    sigma = m.dist.left_truncated_distribution(
        m.dist.student_t(3, 0, 20, create_obj=True), low=0.0, name="sigma", shape=(T,)
    )

    u_phylo = m.effects.phylo_effect(L, sd_phylo, name="z_phylo")

    mu = intercept.reshape(T, 1) + jnp.outer(b_cofactor, X) + u_phylo[:, phylo_idx]
    m.dist.normal(mu, sigma.reshape(T, 1), name="obs",
                  obs=jnp.broadcast_to(Y.reshape(1, N), (T, N)))


print("  Fitting BI model (non-centered, matched priors) ...")
t_start_bi = time.time()
m.fit(model, num_warmup=500, num_samples=500, num_chains=2)
bi_time = time.time() - t_start_bi
print(f"  BI done in {bi_time:.2f} s ({bi_time/NUM_TREES:.3f} s/tree)")

samples = m.posteriors
b_intercept_samp = (
    np.asarray(samples["Intercept"]) - mean_x * np.asarray(samples["b_cofactor"])
)
posteriors = []
for t in range(NUM_TREES):
    posteriors.append(pd.DataFrame({
        "b_Intercept":        b_intercept_samp[:, t],
        "b_x":                np.asarray(samples["b_cofactor"])[:, t],
        "sigma":              np.asarray(samples["sigma"])[:, t],
        "sd_phylo__Intercept": np.asarray(samples["sd_phylo"])[:, t],
        "tree_index":         t + 1,
    }))

df_bi = pd.concat(posteriors)
df_bi.to_csv(BI_CSV, index=False)
print(f"  Saved BI posteriors -> {BI_CSV}")


# ══════════════════════════════════════════════════════════════════════════════
# STEP 2 – brms (via Rscript)
# ══════════════════════════════════════════════════════════════════════════════
print("\n" + "=" * 70)
print(f"  STEP 2 | Running brms baseline on {NUM_TREES} trees (Rscript)")
print("=" * 70)

r_script = os.path.join(SCRIPT_DIR, "brms_baseline.R")
t_start_brms = time.time()
result = subprocess.run(
    ["Rscript", r_script],
    cwd=ROOT_DIR,
    capture_output=True,
    text=True,
)
brms_time = time.time() - t_start_brms
print(result.stdout[-3000:] if len(result.stdout) > 3000 else result.stdout)
if result.returncode != 0:
    print("  [STDERR]", result.stderr[-2000:])
    sys.exit(f"Rscript failed with return code {result.returncode}")
print(f"  brms done in {brms_time:.2f} s ({brms_time/NUM_TREES:.3f} s/tree)")


# ══════════════════════════════════════════════════════════════════════════════
# STEP 3 – Comparative table & Benchmark
# ══════════════════════════════════════════════════════════════════════════════
print("\n" + "=" * 70)
print("  STEP 3 | Comparative table & Benchmark Results")
print("=" * 70)

df_bi   = pd.read_csv(BI_CSV)
df_brms = pd.read_csv(BRMS_CSV)

PARAMS = ["b_Intercept", "b_x", "sigma", "sd_phylo__Intercept"]
PARAM_LABELS = {
    "b_Intercept":          "Intercept (alpha)",
    "b_x":                  "Slope (beta)",
    "sigma":                "Residual SD (sigma)",
    "sd_phylo__Intercept":  "Phylo. SD (sigma_phylo)",
}

rows = []
for p in PARAMS:
    bm   = df_brms[p].mean();  bsd  = df_brms[p].std()
    bim  = df_bi[p].mean();    bisd = df_bi[p].std()
    diff = abs(bm - bim)
    pct  = 100 * diff / (abs(bm) + 1e-12)
    rows.append({
        "Parameter":       PARAM_LABELS[p],
        "brms Mean":       round(bm,   4),
        "brms SD":         round(bsd,  4),
        "BI Mean":         round(bim,  4),
        "BI SD":           round(bisd, 4),
        "Abs. Difference": round(diff, 4),
        "% Difference":    round(pct,  2),
    })

df_table = pd.DataFrame(rows)
df_table.to_csv(TABLE_CSV, index=False)

speedup = brms_time / bi_time if bi_time > 0 else 0

sep = "-" * 90
print(f"\n  {'Parameter':<26} {'brms Mean':>10} {'brms SD':>8} {'BI Mean':>10} {'BI SD':>8}  {'|diff|':>8}  {'%diff':>6}")
print(f"  {sep}")
for _, r in df_table.iterrows():
    print(f"  {r['Parameter']:<26} {r['brms Mean']:>10.4f} {r['brms SD']:>8.4f} "
          f"{r['BI Mean']:>10.4f} {r['BI SD']:>8.4f}  {r['Abs. Difference']:>8.4f}  "
          f"{r['% Difference']:>5.2f}%")
print(f"  {sep}")

print(f"\n  BENCHMARK ({NUM_TREES} Trees)")
print(f"  {sep}")
print(f"  {'Framework':<26} {'Total Time (s)':>15} {'Time/Tree (s)':>15}")
print(f"  {'brms (Stan)':<26} {brms_time:>15.2f} {brms_time/NUM_TREES:>15.3f}")
print(f"  {'BI (NumPyro)':<26} {bi_time:>15.2f} {bi_time/NUM_TREES:>15.3f}")
print(f"  {sep}")
print(f"  BI Relative Speedup: {speedup:.2f}x\n")

with open(TABLE_TXT, "w") as f:
    f.write(f"Phylogenetic Benchmark: BI vs brms\n")
    f.write(f"{NUM_TREES} Trees | 500 warmup | 500 samples | 2 chains\n\n")
    f.write(df_table.to_string(index=False))
    f.write("\n\n--- Performance ---\n")
    f.write(f"brms: {brms_time:.2f} s ({brms_time/NUM_TREES:.3f} s/tree)\n")
    f.write(f"BI:   {bi_time:.2f} s ({bi_time/NUM_TREES:.3f} s/tree)\n")
    f.write(f"BI speedup: {speedup:.2f}x\n")

print(f"  Saved -> {TABLE_CSV}")
print(f"  Saved -> {TABLE_TXT}")


# ══════════════════════════════════════════════════════════════════════════════
# STEP 4 – Density-overlay plot
# ══════════════════════════════════════════════════════════════════════════════
print("\n" + "=" * 70)
print("  STEP 4 | Generating density-overlay plots")
print("=" * 70)

import matplotlib
matplotlib.use("Agg")
import matplotlib.pyplot as plt
from matplotlib.lines import Line2D
import seaborn as sns

CLR_BRMS = "#E07B54"
CLR_BI   = "#4D8FAC"

sns.set_theme(style="white", font_scale=1.15)
fig, axes = plt.subplots(2, 2, figsize=(14, 10))
axes = axes.flatten()

for ax, p in zip(axes, PARAMS):
    brms_vals = df_brms[p].dropna().values
    bi_vals   = df_bi[p].dropna().values
    sns.kdeplot(brms_vals, ax=ax, color=CLR_BRMS, fill=True, alpha=0.35, linewidth=2.5)
    sns.kdeplot(bi_vals,   ax=ax, color=CLR_BI,   fill=True, alpha=0.35, linewidth=2.5)
    ax.axvline(brms_vals.mean(), color=CLR_BRMS, lw=1.8, ls="--", alpha=0.9)
    ax.axvline(bi_vals.mean(),   color=CLR_BI,   lw=1.8, ls="--", alpha=0.9)
    ax.set_title(PARAM_LABELS[p], fontsize=13, fontweight="bold", pad=10)
    ax.set_xlabel("Posterior value", fontsize=11)
    ax.set_ylabel("Density", fontsize=11)
    diff = abs(brms_vals.mean() - bi_vals.mean())
    ax.text(0.97, 0.95,
            f"brms: {brms_vals.mean():.4f}\n BI:  {bi_vals.mean():.4f}\n diff: {diff:.4f}",
            transform=ax.transAxes, ha="right", va="top", fontsize=9,
            bbox=dict(boxstyle="round,pad=0.4", facecolor="white",
                      edgecolor="#cccccc", alpha=0.9))
    ax.spines[["top", "right"]].set_visible(False)

legend_elements = [
    Line2D([0], [0], color=CLR_BRMS, lw=2.5, label="brms"),
    Line2D([0], [0], color=CLR_BI,   lw=2.5, label="BI"),
]
fig.legend(handles=legend_elements, loc="lower center", ncol=2,
           fontsize=13, frameon=False, bbox_to_anchor=(0.5, -0.01))
fig.suptitle(
    f"100-Tree Phylogenetic Benchmark (Non-centered, Matched Priors)\n"
    f"(BI vs brms | BI Speedup: {speedup:.1f}x)",
    fontsize=14, fontweight="bold", y=1.01,
)
plt.tight_layout()
plt.savefig(PLOT_FILE, dpi=300, bbox_inches="tight")
print(f"  Saved -> {PLOT_FILE}")

print("\n" + "=" * 70)
print("  Pipeline complete. Outputs in output/")
print("=" * 70 + "\n")
```
