## Appendix 1, 2, 3, and 4 for "BayesForge: A Bayesian Inference library for Python, R, and Julia": validate_single_tree.py

```
#!/usr/bin/env python3
"""
validate_single_tree.py — Single-tree BI vs brms validation (Box 2)
====================================================================
Fits the identical PGLMM to tree 1 independently in both BI and brms,
then overlays the posterior densities.  Demonstrates that the two
frameworks recover the same posterior when the model and data are identical.

Input  : ../data/pruned_traits.csv, ../data/mammal_100_trees.nexus
Output : ../output/validation_single_tree.png
         ../output/validation_single_tree.csv

Run from any directory:
    python scripts/validate_single_tree.py
"""

import os
import sys
import subprocess

import matplotlib
matplotlib.use("Agg")
import matplotlib.pyplot as plt
from matplotlib.lines import Line2D
from scipy.stats import gaussian_kde

import numpy as np
import pandas as pd
import jax.numpy as jnp
import dendropy
from BayesForge import bf

# ── paths ─────────────────────────────────────────────────────────────────────
SCRIPT_DIR = os.path.dirname(os.path.abspath(__file__))
ROOT_DIR   = os.path.dirname(SCRIPT_DIR)
DATA_DIR   = os.path.join(ROOT_DIR, "data")
OUTPUT_DIR = os.path.join(ROOT_DIR, "output")
os.makedirs(OUTPUT_DIR, exist_ok=True)
os.chdir(ROOT_DIR)

TRAITS_FILE = os.path.join(DATA_DIR, "pruned_traits.csv")
TREES_FILE  = os.path.join(DATA_DIR, "mammal_100_trees.nexus")
OUT_PNG     = os.path.join(OUTPUT_DIR, "validation_single_tree.png")
OUT_CSV     = os.path.join(OUTPUT_DIR, "validation_single_tree.csv")

for f in [TRAITS_FILE, TREES_FILE]:
    if not os.path.exists(f):
        sys.exit(f"Missing: {f}  — run scripts/acquire_data.py first.")

# ── data ──────────────────────────────────────────────────────────────────────
df = pd.read_csv(TRAITS_FILE)
df["phylo_name"] = df["Scientific.Name"].str.replace(" ", "_")
df = df.dropna(subset=["BodyMass_g", "BodyLength_mm"])
for col in ["BodyMass_g", "BodyLength_mm"]:
    df = df[df[col].astype(str).str.strip().str.lower() != "na"]
df["x_raw"] = np.log(df["BodyMass_g"].astype(float))
df["y_raw"] = np.log(df["BodyLength_mm"].astype(float))
df = df.sort_values("phylo_name").reset_index(drop=True)
n_sp = len(df)

# ── VCV for tree 1 (matches ape::vcv.phylo — root edge excluded) ──────────────
print("Building VCV for tree 1 ...")
trees = dendropy.TreeList.get(path=TREES_FILE, schema="nexus")
tree  = trees[0]
for t in tree.taxon_namespace:
    if t.label and t.label.startswith("'") and t.label.endswith("'"):
        t.label = t.label[1:-1]

root_offset = tree.seed_node.edge.length or 0.0
pdm   = tree.phylogenetic_distance_matrix()
taxa  = [t for t in tree.taxon_namespace if t.label in df["phylo_name"].values]
taxa  = sorted(taxa, key=lambda t: t.label)
nodes = {t: tree.find_node(filter_fn=lambda n, t=t: n.taxon == t) for t in taxa}

A = np.zeros((n_sp, n_sp))
for i, t1 in enumerate(taxa):
    A[i, i] = nodes[t1].distance_from_root() - root_offset
    for j, t2 in enumerate(taxa):
        if i < j:
            mrca = pdm.mrca(t1, t2)
            d = (mrca.distance_from_root() - root_offset
                 if hasattr(mrca, "distance_from_root") else 0.0)
            A[i, j] = d
            A[j, i] = d

L = np.linalg.cholesky(A + 1e-8 * np.eye(n_sp))

X_np   = df["x_raw"].values
Y_np   = df["y_raw"].values
mean_x = float(X_np.mean())
X_c    = X_np - mean_x

# ── BI ────────────────────────────────────────────────────────────────────────
print("Fitting BI on tree 1 ...")
m = bf("cpu")
m.data_on_model = dict(
    phen       = jnp.array(Y_np, jnp.float64),
    cofactor   = jnp.array(X_c,  jnp.float64),
    phylo_idx  = jnp.arange(n_sp, dtype=jnp.int32),
    A_cholesky = jnp.array(L,    jnp.float64),
)


def model(phen, cofactor, phylo_idx, A_cholesky):
    intercept  = m.dist.normal(0, 50, name="Intercept")
    b_cofactor = m.dist.normal(0, 10, name="b_cofactor")
    sd_phylo   = m.dist.left_truncated_distribution(
        m.dist.student_t(3, 0, 20, create_obj=True), low=0.0, name="sd_phylo"
    )
    sigma = m.dist.left_truncated_distribution(
        m.dist.student_t(3, 0, 20, create_obj=True), low=0.0, name="sigma"
    )
    u     = m.effects.phylo_effect(A_cholesky, sd_phylo, name="z_phylo")
    mu    = intercept + b_cofactor * cofactor + u[phylo_idx]
    m.dist.normal(mu, sigma, name="obs", obs=phen)


m.fit(model, num_warmup=1000, num_samples=1000, num_chains=2)
sv     = m.posteriors
b_int  = np.asarray(sv["Intercept"]).flatten() - mean_x * np.asarray(sv["b_cofactor"]).flatten()
df_bi  = pd.DataFrame({
    "b_Intercept":         b_int,
    "b_x":                 np.asarray(sv["b_cofactor"]).flatten(),
    "sigma":               np.asarray(sv["sigma"]).flatten(),
    "sd_phylo__Intercept": np.asarray(sv["sd_phylo"]).flatten(),
    "framework":           "BI",
})

# ── brms ──────────────────────────────────────────────────────────────────────
print("Fitting brms on tree 1 ...")
r_code = r"""
suppressPackageStartupMessages({library(brms); library(ape)})
dat <- read.csv("data/pruned_traits.csv", stringsAsFactors=FALSE)
dat$phylo <- gsub(" ", "_", dat$Scientific.Name)
dat$y <- log(as.numeric(dat$BodyLength_mm))
dat$x <- log(as.numeric(dat$BodyMass_g))
dat <- dat[is.finite(dat$y) & is.finite(dat$x), ]
tr <- read.nexus("data/mammal_100_trees.nexus")[[1]]
tr$tip.label <- gsub("'", "", tr$tip.label)
A <- vcv.phylo(tr)
keep <- intersect(dat$phylo, rownames(A))
dat <- dat[dat$phylo %in% keep, ]
A   <- A[keep, keep]
fit <- brm(
    y ~ x + (1 | gr(phylo, cov = A)),
    data = dat, data2 = list(A = A), family = gaussian(),
    prior = c(
        prior(normal(0, 10), class = "b"),
        prior(normal(0, 50), class = "Intercept"),
        prior(student_t(3, 0, 20), class = "sd"),
        prior(student_t(3, 0, 20), class = "sigma")
    ),
    chains = 2, cores = 1, iter = 2000, warmup = 1000,
    seed = 42, refresh = 0, silent = 2
)
post <- as_draws_df(fit)[, c("b_Intercept", "b_x", "sigma", "sd_phylo__Intercept")]
write.csv(post, "output/brms_tree1_tmp.csv", row.names = FALSE)
"""
tmp_r = os.path.join(OUTPUT_DIR, "_val_tmp.R")
with open(tmp_r, "w") as f:
    f.write(r_code)
result = subprocess.run(["Rscript", tmp_r], cwd=ROOT_DIR, capture_output=True, text=True)
os.remove(tmp_r)
if result.returncode != 0:
    sys.exit(f"brms failed:\n{result.stderr[-2000:]}")

df_brms = pd.read_csv(os.path.join(OUTPUT_DIR, "brms_tree1_tmp.csv"))
os.remove(os.path.join(OUTPUT_DIR, "brms_tree1_tmp.csv"))
df_brms["framework"] = "brms"

# ── save combined CSV ─────────────────────────────────────────────────────────
PARAMS = ["b_Intercept", "b_x", "sigma", "sd_phylo__Intercept"]
df_val = pd.concat([df_bi[PARAMS + ["framework"]], df_brms[PARAMS + ["framework"]]],
                   ignore_index=True)
df_val.to_csv(OUT_CSV, index=False)

# ── comparison table ──────────────────────────────────────────────────────────
PARAM_LABELS = {
    "b_Intercept":          "Intercept (alpha)",
    "b_x":                  "Slope (beta)",
    "sigma":                "Residual SD (sigma)",
    "sd_phylo__Intercept":  "Phylo. SD (sigma_phylo)",
}
print(f"\n  {'Parameter':<26} {'brms Mean':>10} {'brms SD':>9} {'BI Mean':>10} {'BI SD':>9}  {'%diff mean':>10}")
for p in PARAMS:
    bm  = df_brms[p].mean(); bsd = df_brms[p].std()
    bim = df_bi[p].mean();   bisd = df_bi[p].std()
    pct = 100 * abs(bm - bim) / (abs(bm) + 1e-12)
    print(f"  {PARAM_LABELS[p]:<24} {bm:>10.4f} {bsd:>9.4f} {bim:>10.4f} {bisd:>9.4f}  {pct:>9.2f}%")

# ── density plot ──────────────────────────────────────────────────────────────
CLR_BRMS = "#E08080"
CLR_BI   = "#5A9AC5"

fig, axes = plt.subplots(2, 2, figsize=(12, 8))
axes = axes.flatten()
titles = [PARAM_LABELS[p] for p in PARAMS]

for ax, p, title in zip(axes, PARAMS, titles):
    vb  = df_brms[p].values
    vbi = df_bi[p].values
    lo = min(vb.min(), vbi.min()); hi = max(vb.max(), vbi.max()); rng = hi - lo
    xs = np.linspace(lo - 0.1 * rng, hi + 0.1 * rng, 400)
    kb  = gaussian_kde(vb)
    kbi = gaussian_kde(vbi)
    ax.fill_between(xs, kb(xs),  alpha=0.3, color=CLR_BRMS)
    ax.fill_between(xs, kbi(xs), alpha=0.3, color=CLR_BI)
    ax.plot(xs, kb(xs),  color=CLR_BRMS, lw=2)
    ax.plot(xs, kbi(xs), color=CLR_BI,   lw=2)
    ax.axvline(vb.mean(),  color=CLR_BRMS, ls="--", lw=1.2)
    ax.axvline(vbi.mean(), color=CLR_BI,   ls="--", lw=1.2)
    ax.set_title(title, fontsize=12, fontweight="bold")
    ax.set_xlabel("Posterior value", fontsize=9)
    ax.set_ylabel("Density", fontsize=9)
    info = (f"brms: {vb.mean():.4f}\n"
            f"BI:   {vbi.mean():.4f}\n"
            f"diff: {abs(vb.mean()-vbi.mean()):.4f}")
    ax.text(0.97, 0.97, info, transform=ax.transAxes, ha="right", va="top",
            fontsize=8, bbox=dict(boxstyle="round,pad=0.4", facecolor="white",
                                  edgecolor="#ccc", alpha=0.9))
    ax.spines[["top", "right"]].set_visible(False)

fig.legend(
    handles=[Line2D([0], [0], color=CLR_BRMS, lw=2.5, label="brms"),
             Line2D([0], [0], color=CLR_BI,   lw=2.5, label="BI")],
    loc="lower center", ncol=2, fontsize=13, frameon=False,
    bbox_to_anchor=(0.5, -0.01),
)
fig.suptitle(
    "Single-Tree Validation (Tree 1): BI vs brms\n"
    "Both frameworks use identical model, priors, and VCV matrix",
    fontsize=12, fontweight="bold", y=1.01,
)
plt.tight_layout()
plt.savefig(OUT_PNG, dpi=300, bbox_inches="tight")
print(f"\nSaved: {OUT_PNG}")
print(f"Saved: {OUT_CSV}")
```
