## Appendix 1, 2, 3, and 4 for "BayesForge: A Bayesian Inference library for Python, R, and Julia": README.md

# SRM Example — Social Relations Model

Generative + inference demo for a count-valued directed network. Simulates
a network from the Social Relations Model, then recovers parameters with
BayesForge.

## Install

```
pip install -r requirements.txt
```

Single environment — no Stan, no R.

## Model

Dyadic count `Y_ij ~ Binomial(E_ij, logit_p)` for each ordered pair, where
`E_ij = E_i + E_j` and, per direction:
```
logit_p(i→j) = alpha + beta_i*x_i + beta_j*x_j + beta_D*d_ij + lambda_i + pi_j + delta_ij
logit_p(j→i) = alpha + beta_i*x_j + beta_j*x_i + beta_D*d_ij + lambda_j + pi_i + delta_ji
```
i.e. each direction combines the *sender's* own random effect (`lambda`) with
the *receiver's* own random effect (`pi`) — matching the standard SRM
decomposition (`S_i + R_j`, `S_j + R_i`) that `m.net.sender_receiver()`
implements internally.

- `alpha` — intercept
- `beta_i, beta_j, beta_D` — fixed slopes (sender/receiver nodal + dyadic covariate)
- `lambda_i, pi_j` — correlated sender/receiver random effects per node (LKJ Cholesky)
- `delta_ij, delta_ji` — correlated dyadic random effects (LKJ Cholesky)
- Exposure `E_i ~ Poisson(exp(0.5 + 1.2 * x_i))`

## Data origin

No external dataset. Data is generated inside `srm_example.py` using
`m.dist.*` sampling (seed 42, N = 100 nodes, N*(N-1)/2 dyads). Ground-truth
parameter values are hard-coded and used both to simulate and to overlay on
the posterior forest plot.

## Reproducibility note

`m = bf(platform="cpu", rand_seed=False)` only seeds `m.dist` deterministically
by itself — `BayesForge>=0.0.57` (or this repo's `BF` source past commit
`84dbfcd0`) additionally propagates that seed into `m.effects`/`m.net`'s
internal random-effect generators, which otherwise silently ignore
`rand_seed` and reseed from wall-clock time on every run. On an older/
unpatched BayesForge, `lambda_i`/`pi_j`/`delta_ij`/`delta_ji` — and therefore
the whole simulated dataset — will differ on every run despite the fixed
seed.

## Files

| File | Purpose |
|------|---------|
| `srm_example.py` | Self-contained script: simulate data → fit BayesForge → forest plot + summary |
| `srm_forest_plot.png` | Posterior forest plot with ground-truth overlay (output) |
| `srm_summary.txt` | Posterior mean / SD / HDI table (output) |

## Run

```
python srm_example.py
```
