## Appendix 1, 2, 3, and 4 for "BayesForge: A Bayesian Inference library for Python, R, and Julia": srm_example.py

```
# %%
"""
SRM (Social Relations Model) — Generative + Inference Example
=============================================================
Simulates a count-valued directed network (Binomial with exposure) using the
Social Relations Model (no block effects), then recovers parameters via
Bayesian inference in the BI framework.

Generative structure:
  - Intercept (alpha)
  - Sender/receiver fixed slopes on nodal covariate x_i    (beta_i, beta_j)
  - Dyadic covariate fixed slope                            (beta_D)
  - Correlated sender/receiver random effects (lambda_i, pi_j) via LKJ Cholesky
    — same as m.net.sender_receiver() internally
  - Correlated dyadic random effects (delta_ij, delta_ji) via LKJ Cholesky
    — same as m.net.dyadic_effect() internally
  - Exposure E_i ~ Poisson(exp(0.5 + 1.2 * x_i))

The inference model mirrors this structure exactly so posteriors can be
compared to the known generative parameter values.
"""

import os
import jax.numpy as jnp
import numpy as np
import pandas as pd
import matplotlib

matplotlib.use("Agg")  # headless / non-interactive backend
import matplotlib.pyplot as plt
from BayesForge import bf

# %%
# ── Reproducibility ───────────────────────────────────────────────────────────
SEED = 42
rng = np.random.default_rng(SEED)
m = bf(platform="cpu", rand_seed=False)

# %%
# ════════════════════════════════════════════════════════════════════════════
# 1.  Generative model
# ════════════════════════════════════════════════════════════════════════════
N = 100
N_dyads = N * (N - 1) // 2  # upper-triangle pairs

# ── 1a. Nodal covariate ───────────────────────────────────────────────────
id_cov = m.dist.normal(0, 1, shape=(N_dyads,), sample=True)
x = m.dist.normal(0, 1, shape=(N,), sample=True)  # (N,)

# ── 1b. Exposure: E_i ~ Poisson(exp(0.5 + 1.2*x_i)) ─────────────────────
E = m.dist.poisson(m.link.exp(0.5 + 1.2 * x), sample=True)  # (N,)

# ── 1c. Dyadic covariate (symmetric, no self-loops) ───────────────────────
d_ij = m.net.mat_to_edgl(m.net.dyad_vec_to_mat(id_cov, symmetric=True))

# %%
# ── 1d. Sender/receiver random effects via varying_effects (zero-centered) ──
lambda_i, pi_j = m.effects.varying_effects(
    N_vars=1,
    N_group=N,
    group_id=jnp.arange(N),
    centered=True,
    alpha_bar=jnp.zeros(1),
    beta_bar=jnp.zeros(1),
    sd_intercept=jnp.array([0.2]),
    sd_beta=jnp.array([0.2]),
    sample=True,
)
# lambda_i : (N,)    — sender random effects, zero-centered
# pi_j     : (N,)    — receiver random effects, zero-centered

# ── 1e. Dyadic random effects via varying_effects (zero-centered) ────────────
delta_ij, delta_ji_raw = m.effects.varying_effects(
    N_vars=1,
    N_group=N_dyads,
    group_id=jnp.arange(N_dyads),
    group_name="dyadic",
    centered=False,
    alpha_bar=jnp.zeros(1),
    beta_bar=jnp.zeros(1),
    sd_intercept=jnp.array([0.2]),
    sd_beta=jnp.array([0.2]),
    sample=True,
)
delta_ji = delta_ji_raw  # (N_dyads,) — already squeezed for N_vars=1

# ── 1f. Fixed effect parameters ───────────────────────────────────────────
alpha, beta_i, beta_j, beta_D = 0, 0.5, -0.3, 0.4

# ── 1g. Edgelist format for covariates and node IDs ──────────────────────
x_edgl = m.net.vec_to_edgl(x)
edgl_ID = m.net.vec_to_edgl(jnp.arange(N))

# ── 1h. Linear predictors on logit scale ─────────────────────────────────
#   direction i→j: sender=i (lambda_i[i]), receiver=j (pi_j[j]) — matches
#   Neteffect.node_effects_to_dyadic_format's S_i + R_j.
logit_ij = (
    alpha
    + beta_i * x_edgl[:, 0]  # sender fixed slope
    + beta_j * x_edgl[:, 1]  # receiver fixed slope
    + beta_D * d_ij[:, 0]  # dyadic covariate
    + lambda_i[edgl_ID[:, 0]]  # sender i's random λ (S_i)
    + pi_j[edgl_ID[:, 1]]  # receiver j's random π (R_j)
    + delta_ij  # dyadic random i→j (one per dyad)
)

#   direction j→i: sender=j (lambda_i[j]), receiver=i (pi_j[i]) — matches
#   Neteffect.node_effects_to_dyadic_format's S_j + R_i.
logit_ji = (
    alpha
    + beta_i * x_edgl[:, 1]
    + beta_j * x_edgl[:, 0]
    + beta_D * d_ij[:, 1]
    + lambda_i[edgl_ID[:, 1]]  # sender j's random λ (S_j)
    + pi_j[edgl_ID[:, 0]]  # receiver i's random π (R_i)
    + delta_ji  # dyadic random j→i (one per dyad)
)

logits_dyadic = jnp.stack([logit_ij, logit_ji], axis=1)

# ── 1i. Binomial draws ────────────────────────────────────────────────────
exposure = m.net.vec_to_edgl(E.astype(int))  # (N_dyads, 2): col0=E_i, col1=E_j
Y_ij = m.dist.binomial(
    total_count=exposure,
    logits=logits_dyadic,
    sample=True,
)


# %%
# ════════════════════════════════════════════════════════════════════════════
# 2.  Inference model
# ════════════════════════════════════════════════════════════════════════════
m.data_on_model = dict(
    network=Y_ij,
    dyadic_predictors=d_ij,
    focal_individual_predictors=x.reshape(N, 1),
    target_individual_predictors=x.reshape(N, 1),
    exposure=exposure,
)


def model(
    network,
    dyadic_predictors,
    focal_individual_predictors,
    target_individual_predictors,
    exposure,
):
    # Intercept
    alpha = m.dist.normal(0, 1, name="alpha")

    # Sender–receiver: fixed slopes + correlated LKJ random effects
    sr = m.net.sender_receiver(
        focal_individual_predictors,
        target_individual_predictors,
    )  # shape (N_dyads, 2)

    # Dyadic: fixed slope on d_ij + correlated LKJ reciprocity
    dr = m.net.dyadic_effect(dyadic_predictors)  # shape (N_dyads, 2)

    # Binomial likelihood (handles count data with exposure)
    m.dist.binomial(total_count=exposure, logits=alpha + sr + dr, obs=network)


print("\n" + "=" * 60)
print("Fitting SRM inference model ...")
print("=" * 60)
m.fit(model, num_samples=1000, num_warmup=1000, num_chains=1, thinning=1)

# %%
print(f"True alpha is {alpha}")
print(f"True sender_effect is {beta_i}")
print(f"True receiver_effect is {beta_j}")
print(f"True dyadic_effect is {beta_D}")
m.summary(
    exclude_vars=[
        "dr_L",
        "sr_rf",
        "dr_raw",
        "sr_raw",
        "dr_rf",
        "sr_L",
        "dr_sigma",
        "sr_sigma",
    ]
)

# %%

# ════════════════════════════════════════════════════════════════════════════
# 3.  Posterior summary — table + forest plot
# ════════════════════════════════════════════════════════════════════════════
SCRIPT_DIR = os.path.dirname(os.path.abspath(__file__))
TABLE_PATH = os.path.join(SCRIPT_DIR, "srm_summary.txt")
PLOT_PATH = os.path.join(SCRIPT_DIR, "srm_forest_plot.png")

post = m.posteriors
print(f"\nPosterior keys: {list(post.keys())}\n")

FOCUS_PARAMS = {"alpha", "dyad_effects", "receiver_effects", "sender_effects"}

_ref_fixed = {
    "sender_effects": beta_i,
    "receiver_effects": beta_j,
    "dyad_effects": beta_D,
}
true_vals = {"alpha": alpha}
true_fixed = {
    "alpha": alpha,
    "beta_sender": beta_i,
    "beta_receiver": beta_j,
    "beta_dyad": beta_D,
}


# ── Helpers ───────────────────────────────────────────────────────────────
def expand_param(pname, arr):
    """Yield (label, flat_samples) for each scalar element of a posterior array."""
    arr = np.array(arr)
    if arr.ndim == 1:
        yield pname, arr
    else:
        for idx in np.ndindex(arr.shape[1:]):
            label = pname + "[" + ",".join(map(str, idx)) + "]"
            samples = arr[(slice(None),) + idx].flatten()
            yield label, samples


# ── 3a. Build summary dataframe (focused params only) ────────────────────
rows = []
for pname, arr in post.items():
    if pname not in FOCUS_PARAMS:
        continue
    arr_np = np.array(arr)
    for label, samples in expand_param(pname, arr_np):
        rows.append(
            {
                "parameter": label,
                "mean": float(samples.mean()),
                "sd": float(samples.std()),
                "2.5%": float(np.percentile(samples, 2.5)),
                "50%": float(np.median(samples)),
                "97.5%": float(np.percentile(samples, 97.5)),
            }
        )

summary_df = pd.DataFrame(rows).set_index("parameter").round(3)


# ── 3b. Print and save table ──────────────────────────────────────────────
print(summary_df.to_string())
with open(TABLE_PATH, "w") as fh:
    fh.write("SRM Posterior Summary\n")
    fh.write("=" * 72 + "\n")
    fh.write(summary_df.to_string())
    fh.write("\n\nTrue generative values (fixed parameters):\n")
    for k, v in true_fixed.items():
        fh.write(f"  {k:30s} = {v:+.4f}\n")
    fh.write(
        "\nNote: sender/receiver/dyad_effects are random effects (z-scores * sigma*L);\n"
    )
    fh.write(
        "      per-node true values are not directly comparable to posterior scale.\n"
    )
print(f"\n[Output] Summary table -> {TABLE_PATH}")

# ── 3c. Forest plot ───────────────────────────────────────────────────────
ORDERED_VARS = [
    "dyad_effects",
    "receiver_effects",
    "sender_effects",
    "alpha",
]

# Populate true_vals for random effects using fixed slopes as reference
for pname, ref_val in _ref_fixed.items():
    if pname in post:
        for label, _ in expand_param(pname, np.array(post[pname])):
            true_vals[label] = ref_val

plot_entries = []  # (label, mean, lo_err, hi_err)
for pname in ORDERED_VARS:
    if pname not in post:
        continue
    for label, samples in expand_param(pname, np.array(post[pname])):
        m_val = float(samples.mean())
        lo_err = m_val - float(np.percentile(samples, 2.5))
        hi_err = float(np.percentile(samples, 97.5)) - m_val
        plot_entries.append((label, m_val, lo_err, hi_err))

n_params = len(plot_entries)
fig_h = max(6, n_params * 0.45 + 2)
fig, ax = plt.subplots(figsize=(9, fig_h))

labels = [e[0] for e in plot_entries]
means = np.array([e[1] for e in plot_entries])
lo_errs = np.array([e[2] for e in plot_entries])
hi_errs = np.array([e[3] for e in plot_entries])
y_pos = np.arange(n_params)

ax.errorbar(
    means,
    y_pos,
    xerr=[lo_errs, hi_errs],
    fmt="s",
    color="#1f77b4",
    capsize=3,
    markersize=5,
    linewidth=1.2,
    label="BI posterior (mean ± 95% CI)",
)

# Overlay true generative values as red diamonds
first_true = True
for i, lbl in enumerate(labels):
    if lbl in true_vals:
        ax.plot(
            true_vals[lbl],
            y_pos[i],
            "D",
            color="#d62728",
            markersize=7,
            zorder=5,
            label="True (generative)" if first_true else "_nolegend_",
        )
        first_true = False

ax.axvline(0, color="k", ls="--", alpha=0.35, lw=1)
ax.set_yticks(y_pos)
ax.set_yticklabels(labels, fontsize=8)
ax.set_xlabel("Parameter value")
ax.set_title(
    f"SRM Inference — Posterior Summary  (N={N}, 1000 samples)\n"
    "Red diamonds = true generative values",
    fontsize=10,
)
ax.legend(fontsize=8, loc="lower right")
ax.invert_yaxis()  # top-to-bottom ordering (matches reference plot)
plt.tight_layout()
plt.savefig(PLOT_PATH, dpi=150)
plt.close()
print(f"[Output] Forest plot      -> {PLOT_PATH}")
print("\nDone.")

# %%
```
