## Appendix 1, 2, 3, and 4 for "BayesForge: A Bayesian Inference library for Python, R, and Julia": 01_model.stan

```
data{
    int N; // individuals
    int M; // tokens
    int J; // number of recordings
    vector[J] d; // durations of each recording
    array[J] int id; // individual
    array[J,M] int Y; // observed token counts
}
transformed data{
    // gather all recordings of an individual into one aggregated observation
    vector[N] dd_agg = rep_vector(0, N);
    array[N,M] int Y_agg = rep_array(0, N, M);
    for ( j in 1:J ) {
        dd_agg[id[j]] += d[j];
        for ( m in 1:M ) Y_agg[id[j],m] += Y[j,m];
    }//j
}
parameters {
    vector<lower=0>[M] L; // rates of each token, conditional on individual possessing it
    vector<lower=0,upper=1>[M] p; // population probabilities of each token in repertoire
}
model{
    L ~ exponential(1);
    p ~ beta(2,2);
    // loop over individuals using their aggregated recordings
    for ( i in 1:N ) {
        for ( m in 1:M ) {
            if ( Y_agg[i,m] >0 ) {
                // observed non-zero count, estimate rate
                Y_agg[i,m] ~ poisson( L[m] * dd_agg[i] );
                target += log(p[m]);
            } else {
                // observed zero so account for mixture
                target += log_sum_exp(
                    log(p[m]) + poisson_lpmf(0|L[m]*dd_agg[i]) , // has it, but produced zero
                    log1m(p[m]) // doesn't have it
                );
            }
        }//m
    }//i
}
generated quantities {
    // calculate posterior prob of each token for each individual in sample
    // this allows estimation of undercounting due to sampling and rate variation
    matrix[N,M] phi;
    vector[N] R; // repertiore size
    for ( i in 1:N ) {
        vector[2] pp;
        real Z;
        for ( m in 1:M ) {
            // Pr(p[m]==1|Y) = Pr(Y|p[m]==1) p[m] / Z
            // Z = Pr(Y|p[m]==1) p[m] + Pr(Y|p[m]==0) (1-p[m])
            if ( Y_agg[i,m]>0 )
                // individual has it for sure, so no calculation needed
                phi[i,m] = 1;
            else {
                // bayes it up
                pp[1] = log(p[m]) + poisson_lpmf(0|L[m]*dd_agg[i]);
                pp[2] = log1m(p[m]);
                Z = log_sum_exp( pp[1] , pp[2] );
                phi[i,m] = exp( pp[1] - Z );
            }
        }//m
        R[i] = sum( bernoulli_rng( phi[i,] ) );
    }//i
}
```
