## Appendix 1, 2, 3, and 4 for "BayesForge: A Bayesian Inference library for Python, R, and Julia": 02_run_stan.py

```
import json
import pandas as pd
from cmdstanpy import CmdStanModel


def main():
    with open('00_data.json', 'r') as f:
        data = json.load(f)

    print("Running Stan model...", flush=True)
    model = CmdStanModel(stan_file='01_model.stan')
    fit = model.sample(
        data='00_data.json',
        seed=42,
        iter_sampling=1000,
        iter_warmup=1000,
        chains=2,
    )

    draws = fit.draws_pd()
    draws.to_csv('stan_samples.csv', index=False)

    summary = fit.summary()
    L_means = [summary.loc[f'L[{i+1}]', 'Mean'] for i in range(data['M'])]
    p_means = [summary.loc[f'p[{i+1}]', 'Mean'] for i in range(data['M'])]
    pd.DataFrame({'L_mean': L_means, 'p_mean': p_means}).to_csv('stan_means.csv', index=False)

    print("Saved: stan_samples.csv, stan_means.csv")


if __name__ == "__main__":
    main()
```
