## Appendix 1, 2, 3, and 4 for "BayesForge: A Bayesian Inference library for Python, R, and Julia": 03_run_bi.py

```
import json
import numpy as np
import pandas as pd
import jax.numpy as jnp
import jax

from BayesForge import bf

def main():
    with open('00_data.json', 'r') as f:
        data = json.load(f)

    N = data['N']
    M = data['M']
    id_idx = jnp.array(data['id']) - 1
    d_arr = jnp.array(data['d'])
    Y_arr = jnp.array(data['Y'], dtype=jnp.int32)

    # Gather all recordings of an individual into one aggregated observation
    dd_agg = jax.ops.segment_sum(d_arr, id_idx, num_segments=N)  # shape (N,)
    Y_agg = jax.ops.segment_sum(Y_arr, id_idx, num_segments=N)   # shape (N, M)

    bi_data = {
        'N': N,
        'M': M,
        'dd_agg': dd_agg,
        'Y_agg': Y_agg,
    }

    m = bf('cpu', print_devices_found=False)

    def bi_model(N, M, dd_agg, Y_agg):
        with m.dist.plate('tokens', M):
            L = m.dist.exponential(1.0, name='L')
            p = m.dist.beta(2.0, 2.0, name='p')

            # ZIP evaluated once per individual, on their aggregated recordings
            with m.dist.plate('individuals', N):
                Z_ind = m.dist.bernoulli(p, name='Z_ind')
                rate = dd_agg[:, None] * L[None, :]
                gate = 1.0 - Z_ind
                m.dist.zero_inflated_poisson(gate=gate, rate=rate, obs=Y_agg, name='Y')

    print("Fitting BI model (matching Stan's individual-level aggregated ZIP likelihood)...")
    m.fit(bi_model, obs=bi_data, num_samples=1000, num_warmup=1000, num_chains=2, seed=42)

    # Recover Z_ind directly from the model's posterior via exact discrete-site
    # sampling (conditioned on Y_agg), instead of deriving phi by hand.
    print("Sampling Z_ind posterior (infer_discrete=True)...")
    draws = m.sample(data=bi_data, posterior=True, seed=42, infer_discrete=True)
    Z_ind = draws['Z_ind']              # shape (2000, N, M)
    R_posterior = Z_ind.sum(axis=-1)    # shape (2000, N)

    print("Z_ind posterior shape:", Z_ind.shape)
    print("R posterior shape:", R_posterior.shape)
    
    # Save the full posterior explicitly
    posteriors = dict(m.posteriors)
    # We rename Z_ind to 'Z' for compatibility with the plotting script
    posteriors['Z'] = Z_ind
    posteriors['R'] = R_posterior
    jnp.savez('bi_samples.npz', **posteriors)
    
    # Calculate and save posterior means of R
    R_means = np.array(R_posterior.mean(axis=0))
    np.savetxt("bi_R_means.csv", R_means, delimiter=",")
    print("Done! Repertoire sizes saved to bi_R_means.csv")

if __name__ == "__main__":
    main()
```
