## Appendix 1, 2, 3, and 4 for "BayesForge: A Bayesian Inference library for Python, R, and Julia": 04_density_kl.py

```
import json
import numpy as np
import pandas as pd
import matplotlib.pyplot as plt
from scipy.stats import gaussian_kde, entropy

EPS = 1e-10
N_BINS = 60


def kl_divergence(stan_samples, bi_samples, discrete=False):
    """KL(Stan || BF), estimated via shared-bin histograms."""
    lo = min(stan_samples.min(), bi_samples.min())
    hi = max(stan_samples.max(), bi_samples.max())
    if discrete:
        bins = np.arange(lo, hi + 2) - 0.5
    else:
        bins = np.linspace(lo, hi, N_BINS + 1)

    p_hist, _ = np.histogram(stan_samples, bins=bins, density=True)
    q_hist, _ = np.histogram(bi_samples, bins=bins, density=True)
    p_hist = p_hist + EPS
    q_hist = q_hist + EPS
    return entropy(p_hist, q_hist)


def plot_grid(param_name, stan_mat, bi_mat, n_items, kl_vals, discrete=False):
    ncols = 5
    nrows = int(np.ceil(n_items / ncols))
    fig, axes = plt.subplots(nrows, ncols, figsize=(ncols * 3.2, nrows * 2.4))
    axes = np.atleast_2d(axes)

    for i in range(nrows * ncols):
        r, c = divmod(i, ncols)
        ax = axes[r, c]
        if i >= n_items:
            ax.axis('off')
            continue

        s = stan_mat[:, i]
        b = bi_mat[:, i]

        if discrete:
            lo = min(s.min(), b.min())
            hi = max(s.max(), b.max())
            bins = np.arange(lo, hi + 2) - 0.5
            ax.hist(s, bins=bins, density=True, alpha=0.5, color='blue', label='Stan')
            ax.hist(b, bins=bins, density=True, alpha=0.5, color='orange', label='BF')
        else:
            grid = np.linspace(min(s.min(), b.min()), max(s.max(), b.max()), 200)
            ax.plot(grid, gaussian_kde(s)(grid), color='blue', label='Stan')
            ax.plot(grid, gaussian_kde(b)(grid), color='orange', label='BF')

        ax.set_title(f'{param_name}[{i}]  KL={kl_vals[i]:.4f}', fontsize=9)
        ax.tick_params(labelsize=7)

    handles, labels = axes[0, 0].get_legend_handles_labels()
    fig.legend(handles, labels, loc='upper center', bbox_to_anchor=(0.5, 1.02),
               ncol=2, frameon=False, fontsize=12)
    plt.tight_layout(rect=[0, 0, 1, 0.98])
    out = f'density_{param_name}.png'
    plt.savefig(out, dpi=150, bbox_inches='tight')
    plt.close(fig)
    print(f"Saved: {out}")


def main():
    with open('00_data.json', 'r') as f:
        data = json.load(f)
    M, N = data['M'], data['N']

    stan_samples = pd.read_csv('stan_samples.csv')
    bi_samples = np.load('bi_samples.npz')

    stan_L = np.column_stack([stan_samples[f'L[{i+1}]'].values for i in range(M)])
    stan_p = np.column_stack([stan_samples[f'p[{i+1}]'].values for i in range(M)])
    stan_R = np.column_stack([stan_samples[f'R[{i+1}]'].values for i in range(N)]).astype(int)

    bi_L = bi_samples['L']
    bi_p = bi_samples['p']
    bi_R = bi_samples['R'].astype(int)

    with open('log.txt', 'w') as f:
        f.write("Parameter | Stan Mean | BF Mean | Difference\n")
        f.write("-" * 50 + "\n")
        f.write("--- L (Rates) ---\n")
        for i in range(M):
            s, b = stan_L[:, i].mean(), bi_L[:, i].mean()
            f.write(f"L[{i}] | {s:.4f} | {b:.4f} | {s - b:.4f}\n")

        f.write("\n--- p (Probabilities) ---\n")
        for i in range(M):
            s, b = stan_p[:, i].mean(), bi_p[:, i].mean()
            f.write(f"p[{i}] | {s:.4f} | {b:.4f} | {s - b:.4f}\n")

        f.write("\n--- R (Repertoire Size per Individual) ---\n")
        for i in range(N):
            s, b = stan_R[:, i].mean(), bi_R[:, i].mean()
            f.write(f"R[{i}] | {s:.4f} | {b:.4f} | {s - b:.4f}\n")
    print("Saved: log.txt")

    results = []
    kl_L = [kl_divergence(stan_L[:, i], bi_L[:, i]) for i in range(M)]
    kl_p = [kl_divergence(stan_p[:, i], bi_p[:, i]) for i in range(M)]
    kl_R = [kl_divergence(stan_R[:, i], bi_R[:, i], discrete=True) for i in range(N)]

    for i in range(M):
        results.append(('L', i, kl_L[i]))
    for i in range(M):
        results.append(('p', i, kl_p[i]))
    for i in range(N):
        results.append(('R', i, kl_R[i]))

    df = pd.DataFrame(results, columns=['parameter', 'index', 'kl_stan_bf'])
    df.to_csv('kl_divergence.csv', index=False)
    print("Saved: kl_divergence.csv")
    print(f"Mean KL  L: {np.mean(kl_L):.4f}  p: {np.mean(kl_p):.4f}  R: {np.mean(kl_R):.4f}")

    plot_grid('L', stan_L, bi_L, M, kl_L, discrete=False)
    plot_grid('p', stan_p, bi_p, M, kl_p, discrete=False)
    plot_grid('R', stan_R, bi_R, N, kl_R, discrete=True)


if __name__ == "__main__":
    main()
```
