## Appendix 1, 2, 3, and 4 for "BayesForge: A Bayesian Inference library for Python, R, and Julia": README.md

# Voice Example — BayesForge vs Stan

Zero-inflated Poisson model for vocal repertoires. Estimates signal repertoire size at the individual level while accounting for undersampling (observation duration, recording count, vocalization rate, signal commonness). Counts of M tokens across J recordings from N individuals, durations `d`. Each token has rate `L[m]` (Exponential(1) prior) gated by population probability `p[m]` (Beta(2,2) prior). Recordings are aggregated to the individual level (Poisson additivity makes this exact, not an approximation) before fitting: `Y_agg[i,m] ~ ZIPoisson(gate = 1 - p[m], rate = L[m] * dd_agg[i])`.

## Source

Model designed by Richard McElreath. Stan code from `https://github.com/rmcelreath/cg_vocal_repertoires` (file `model0.stan`), adapted here to aggregate recordings per individual before fitting (`01_model.stan`). Used in published analysis:

> Sigmundson, R., Girard-Buttoz, C., Le Floch, A., Souha Azaiez, T., McElreath, R., Zuberbühler, K., Wittig, R. M., & Crockford, C. (2025). Vocal sequence diversity and length remain stable across ontogeny in a catarrhine monkey (Cercocebus atys). *Communications Biology*, 8:465. https://doi.org/10.1038/s42003-025-07922-2

## Data origin

`00_data.json` is **simulated** synthetic data matching the validation setup in the upstream repo (`https://github.com/rmcelreath/cg_vocal_repertoires/blob/main/simulation.r`): 50 individuals (`N`), 50 utterance types (`M`), ~10 recordings per individual (`J = 504`), ground-truth `lambda` and `p` drawn from the model priors. JSON fields match Stan input: `N, M, J, d[J], id[J], Y[J,M]`.

Ground-truth `true_L.csv`/`true_p.csv` referenced by older versions of this example are **not present** in this repo (gitignored, no generator script tracked) — `04_density_kl.py` does not depend on them.

## Environments

This example needs **two separate Python environments** — Stan (`cmdstanpy`) and BayesForge (JAX/NumPyro) have incompatible dependency trees (different numpyro/jax pins) and are not installable together.

```
python -m venv .venv-stan && .venv-stan/Scripts/pip install -r requirements-stan.txt
python -m venv .venv-bf   && .venv-bf/Scripts/pip install -r requirements-bf.txt
```

`requirements-stan.txt` also needs CmdStan itself (not a pip package):
```
.venv-stan/Scripts/python -m cmdstanpy.install_cmdstan
```
Verified against CmdStan 2.36.0.

`requirements-bf.txt` pins `funsor>=0.4.8` specifically — `03_run_bi.py`'s `infer_discrete=True` call crashes under funsor<=0.4.5 with `TypeError: clip() got an unexpected keyword argument 'a_max'` (funsor calls the numpy 1.x-style `np.clip` kwargs; jax>=0.10.0, which BayesForge requires, dropped those aliases from `jax.numpy.clip`). See `requirements-bf.txt` for the full note.

## Files

| File | Purpose |
|-----------------------------|-------------------------------------------|
| `00_data.json` | Simulated Stan input (counts + metadata) |
| `01_model.stan` | Stan model (individual-aggregated ZIP likelihood + phi/R generated quantities) |
| `02_run_stan.py` | Fit Stan (cmdstanpy) — `requirements-stan.txt` env. Writes `stan_samples.csv`, `stan_means.csv` |
| `03_run_bi.py` | Fit BayesForge (JAX/NumPyro) — `requirements-bf.txt` env. Writes `bi_samples.npz`, `bi_means.csv`, `bi_R_means.csv` |
| `04_density_kl.py` | Load both fits — either env (only needs pandas/numpy/matplotlib/scipy). Writes `log.txt` (mean/diff table), `kl_divergence.csv`, `density_L.png`, `density_p.png`, `density_R.png` |

## Run

```
.venv-stan/Scripts/python 02_run_stan.py
.venv-bf/Scripts/python   03_run_bi.py
.venv-stan/Scripts/python 04_density_kl.py   # or .venv-bf, either works
```

Seed 42. Stan: 2 chains, 1000 warmup + 1000 sampling. BayesForge: 2 chains, 1000 warmup + 1000 sampling, repertoire size `R` read directly from the posterior via `m.sample(infer_discrete=True)` (exact discrete-site sampling conditioned on the data, no manual marginal derivation needed) rather than a hand-derived Bayes'-rule formula.
