## Appendix 1, 2, 3, and 4 for "BayesForge: A Bayesian Inference library for Python, R, and Julia": requirements-bf.txt

# Python env for 03_run_bi.py and 04_density_kl.py (BayesForge / JAX side).
# Verified working combination as of 2026-08-14.
#
# funsor==0.4.8 is required, not just "latest": funsor<=0.4.5 calls
# np.clip(..., a_min=..., a_max=...), and jax>=0.10.0 (required by BayesForge)
# dropped those legacy aliases from jax.numpy.clip, so 03_run_bi.py's
# infer_discrete=True path crashes with
#   TypeError: clip() got an unexpected keyword argument 'a_max'
# under any funsor <= 0.4.5. 0.4.8 fixes it.

BayesForge>=0.0.56
jax>=0.10.0
jaxlib>=0.10.0
numpyro>=0.21.0
funsor>=0.4.8
arviz>=1.2.0
numpy>=2.2.0
pandas
scipy
