## Appendix 1, 2, 3, and 4 for "BayesForge: A Bayesian Inference library for Python, R, and Julia": test_aggregated.py

```
#%%
import os, sys, json, time
import numpy as np
import jax
import jax.numpy as jnp
from jax import random
from numpyro.infer import Predictive

sys.path.append(os.path.abspath(os.path.join(os.getcwd(), "..")))
from BI import bi
#%%
with open('00_data.json') as f:
    data = json.load(f)

N, M, J = data['N'], data['M'], data['J']
id_idx  = jnp.array(data['id']) - 1                       # (J,) 0-based
d_arr   = jnp.array(data['d'])                            # (J,)
Y_arr   = jnp.array(data['Y'], dtype=jnp.int32)           # (J, M)

# --- Pre-aggregate recordings per individual BEFORE model ---
# Sum Y and d across all recordings belonging to same individual.
# Result: one row per individual instead of one row per recording.
# Y_agg[i,m] = total count of token m for individual i across all recordings
# D[i]       = total recording duration for individual i
Y_agg = jax.ops.segment_sum(Y_arr, id_idx, num_segments=N)  # (N, M)
D     = jax.ops.segment_sum(d_arr, id_idx, num_segments=N)  # (N,)

bi_data = {'N': N, 'M': M, 'Y_agg': Y_agg, 'D': D}

m = bi('cpu', print_devices_found=False)

def bi_model(**d):
    N, M = d['N'], d['M']

    with m.dist.plate('tokens', M, dim=-1):
        L = m.dist.exponential(1.0, name='L')   # (M,)
        p = m.dist.beta(2.0, 2.0,  name='p')    # (M,)

        with m.dist.plate('individuals', N, dim=-2):
            # Z[i,m] = 1 if individual i possesses token m — directly individual-level.
            # Marginalized automatically by BI during NUTS.
            Z    = m.dist.bernoulli(p, name='Z')           # (N, M)
            # Aggregated rate: total expected count for individual i if they possess token m.
            # Sum of Poissons = Poisson with summed rates → rate = L[m] * D[i].
            rate = d['D'][:, None] * L[None, :]            # (N, M)
            gate = 1.0 - Z                                  # structural zero if Z=0
            m.dist.zero_inflated_poisson(
                gate=gate, rate=rate, obs=d['Y_agg'], name='Y')

print(f"N={N} M={M} — aggregated data (N×M), Z directly individual-level")
print("Fitting...", flush=True)
t0 = time.time()
m.fit(bi_model, obs=bi_data, num_samples=1000, num_warmup=1000, num_chains=2, seed=42)
t1 = time.time()
print(f"Fit time: {t1-t0:.1f}s")

# --- Recover Z via Predictive — no post-hoc formula needed ---
# Z[i,m] is already individual-level; rowsum gives repertoire size directly.
print("Recovering Z via Predictive...", flush=True)
t2 = time.time()
predictive = Predictive(bi_model, m.posteriors, infer_discrete=True)
samples    = predictive(random.PRNGKey(42), **bi_data)
t3 = time.time()
print(f"Predictive time: {t3-t2:.1f}s")

Z_post = np.array(samples['Z'])          # (S, N, M)
R      = Z_post.sum(axis=-1).astype(float)  # (S, N) — row sum = individual repertoire size

print(f"Z posterior shape: {Z_post.shape}")
print(f"R mean (first 5):  {R.mean(0)[:5].round(2)}")

ref = np.load('bi_samples.npz')
print(f"R mean ref recording-level (first 5): {ref['R'].mean(0)[:5].round(2)}")

import pandas as pd
stan = pd.read_csv('stan_samples.csv')
stan_R = np.array([stan[f'R[{i+1}]'].values for i in range(N)]).T  # (S, N)
print(f"R mean Stan         (first 5): {stan_R.mean(0)[:5].round(2)}")
```
