## Appendix 1, 2, 3, and 4 for "BayesForge: A Bayesian Inference library for Python, R, and Julia": test_bf_sample_Z.py

```
import os
import sys
import json
import time
import numpy as np
import pandas as pd
import jax.numpy as jnp

sys.path.append(os.path.abspath(os.path.join(os.getcwd(), "..")))
from BI import bi

with open('00_data.json') as f:
    data = json.load(f)

N, M = data['N'], data['M']
id_idx = jnp.array(data['id']) - 1
d_arr = jnp.array(data['d'])
Y_arr = jnp.array(data['Y'], dtype=jnp.int32)

import jax
dd_agg = jax.ops.segment_sum(d_arr, id_idx, num_segments=N)
Y_agg = jax.ops.segment_sum(Y_arr, id_idx, num_segments=N)

bi_data = {'N': N, 'M': M, 'dd_agg': dd_agg, 'Y_agg': Y_agg}

m = bi('cpu', print_devices_found=False)

def bi_model(N, M, dd_agg, Y_agg):
    with m.dist.plate('tokens', M):
        L = m.dist.exponential(1.0, name='L')
        p = m.dist.beta(2.0, 2.0, name='p')
        with m.dist.plate('individuals', N):
            Z_ind = m.dist.bernoulli(p, name='Z_ind')
            rate = dd_agg[:, None] * L[None, :]
            gate = 1.0 - Z_ind
            m.dist.zero_inflated_poisson(gate=gate, rate=rate, obs=Y_agg, name='Y')

print("Fitting...", flush=True)
t0 = time.time()
m.fit(bi_model, obs=bi_data, num_samples=1000, num_warmup=1000, num_chains=2, seed=42)
print(f"Fit time: {time.time()-t0:.1f}s")

print("m.sample(infer_discrete=True) — exact posterior draw of Z conditioned on Y_agg...", flush=True)
t1 = time.time()
draws = m.sample(data=bi_data, posterior=True, seed=42, infer_discrete=True)
print(f"Sample time: {time.time()-t1:.1f}s")

Z_pred = np.array(draws['Z_ind'])   # (S, N, M) or (1, S, N, M) depending on wrapping
print("Z_ind draw shape:", Z_pred.shape)
Z_pred = Z_pred.reshape(-1, N, M)
R_pred = Z_pred.sum(axis=-1)        # (S, N)
R_pred_mean = R_pred.mean(axis=0)

stan_means = pd.read_csv('stan_means.csv') if os.path.exists('stan_means.csv') else None
stan_samples = pd.read_csv('stan_samples.csv')
stan_R_mean = np.array([stan_samples[f'R[{i+1}]'].mean() for i in range(N)])

print("\nR mean, m.sample(infer_discrete=True) vs Stan (Bayes-corrected):")
for i in range(min(10, N)):
    print(f"R[{i}] | bf.sample={R_pred_mean[i]:.4f} | stan={stan_R_mean[i]:.4f} | diff={R_pred_mean[i]-stan_R_mean[i]:.4f}")

print(f"\nOverall mean: bf.sample={R_pred_mean.mean():.4f}  stan={stan_R_mean.mean():.4f}  diff={R_pred_mean.mean()-stan_R_mean.mean():.4f}")
print(f"Max abs diff: {np.abs(R_pred_mean - stan_R_mean).max():.4f}")
print(f"Correlation:  {np.corrcoef(R_pred_mean, stan_R_mean)[0,1]:.6f}")
```
