## Appendix 1, 2, 3, and 4 for "BayesForge: A Bayesian Inference library for Python, R, and Julia": test_individual_Z.py

```
import os, sys, json, time
import numpy as np
import jax.numpy as jnp
from jax import random
import numpyro
import numpyro.distributions as ndist
from numpyro.infer import Predictive

sys.path.append(os.path.abspath(os.path.join(os.getcwd(), "..")))
from BI import bi

with open('00_data.json') as f:
    data = json.load(f)

N, M, J   = data['N'], data['M'], data['J']
id_idx    = jnp.array(data['id']) - 1        # (J,) 0-based individual index per recording
d_arr     = jnp.array(data['d'])             # (J,)
Y_arr     = jnp.array(data['Y'], dtype=jnp.int32)  # (J, M)

bi_data = {'N': N, 'M': M, 'J': J, 'd': d_arr, 'Y': Y_arr, 'id_idx': id_idx}

m = bi('cpu', print_devices_found=False)

def bi_model(**d):
    N, M = d['N'], d['M']
    id_idx = d['id_idx']

    with m.dist.plate('tokens', M, dim=-1):
        L = m.dist.exponential(1.0, name='L')   # (M,)
        p = m.dist.beta(2.0, 2.0,  name='p')    # (M,)

        with m.dist.plate('individuals', N, dim=-2):
            Z = m.dist.bernoulli(p, name='Z')   # (N, M) — enumerated during NUTS

    # Map individual Z[i,m] → recording Z[j,m] via id_idx.
    # jnp.take(..., axis=-2) is enumeration-safe: works regardless of leading
    # enumeration dims prepended by numpyro (operates on the N dimension only).
    Z_rec = jnp.take(Z, id_idx, axis=-2)        # (..., J, M)

    rate = d['d'][:, None] * L[None, :]          # (J, M)
    gate = 1.0 - Z_rec                            # (..., J, M)

    # Manual log-likelihood via numpyro.factor — bypasses plate obs tracking,
    # which breaks on fancy-indexed tensors.
    log_prob = ndist.ZeroInflatedPoisson(gate=gate, rate=rate).log_prob(d['Y'])
    numpyro.factor('Y_obs', log_prob.sum(axis=(-1, -2)))

print(f"N={N} M={M} J={J} — individual Z[i,m] mapped to recordings via id_idx")
print("Fitting...", flush=True)
t0 = time.time()
m.fit(bi_model, obs=bi_data, num_samples=500, num_warmup=500, num_chains=1, seed=42)
t1 = time.time()
print(f"Fit time: {t1-t0:.1f}s")

print("Predictive (infer_discrete=True)...", flush=True)
t2 = time.time()
predictive = Predictive(bi_model, m.posteriors, infer_discrete=True)
samples    = predictive(random.PRNGKey(42), **bi_data)
t3 = time.time()
print(f"Predictive time: {t3-t2:.1f}s")

Z_post = np.array(samples['Z'])           # (S, N, M)
R      = Z_post.sum(axis=-1)              # (S, N) — no post-hoc formula needed

print(f"Z posterior shape: {Z_post.shape}")
print(f"R mean (first 5):  {R.mean(0)[:5].round(2)}")

ref = np.load('bi_samples.npz')
print(f"R mean ref (first 5, recording-level): {ref['R'].mean(0)[:5].round(2)}")
```
