## Appendix 1, 2, 3, and 4 for "BayesForge: A Bayesian Inference library for Python, R, and Julia": README.md

# Benchmark

Validation and performance benchmarking for the SRM (Social Relations Model)
implementation, distinct from `../Manuscript/Examples/SRM/` (the small,
pedagogical, self-contained example behind Box 1). This folder cross-checks
BayesForge's SRM fit against independent implementations (Stan, STRAND) and
measures scaling performance.

**Not independently re-verified in this pass** — this README documents the
existing file structure and purpose from reading the scripts; none of these
were rerun this session (STRAND/rpy2/R and additional Stan models aren't set
up in this environment). Treat descriptions of what each script *does* as
accurate; treat any implied "this currently runs cleanly" as unverified.

## Structure

```
Benchmark/
├── README.md                 (this file)
└── SRM/
    ├── SIM.py                 BF (long + wide param.) vs STRAND (R, via rpy2) — Linux only
    ├── fit_comparison.py      BF vs 2 Stan formulations (original_srm.stan, STAN2.stan)
    ├── original_srm.stan      Stan SRM, "original" formulation
    ├── original_srm           compiled CmdStan binary for the above
    ├── original_clean.py      empty (0 bytes) — stub/leftover, not used
    ├── test_strand.R          scratch: checks STRAND::make_strand_data() output shape
    ├── test_strand_2.R        scratch: same, sourced from test_strand.R
    ├── original_fit_comparison.txt   empty (0 bytes) — stub/leftover
    ├── log.txt                 output of SIM.py: posterior-mean comparison table
    │                            (STRAND vs BF-long vs BF-wide), one row per parameter
    ├── forest_plot.png         output of SIM.py
    ├── forest_plot_nodal_re.png          output of SIM.py
    ├── forest_plot_nodal_re_scatter.png  output of SIM.py
    ├── stan V BF.png            output of fit_comparison.py
    └── benchmark_suite/        scaling/performance benchmark (own README.md)
        ├── benchmark_stan2.py        CPU backends: STAN2 / Stan Original / BF
        ├── benchmark_gpu_only.py     GPU backend: BF (JAX)
        ├── plot_benchmark.py         → benchmark_plot_cpu.png (wall-clock time)
        ├── plot_efficiency.py        → benchmark_efficiency.png (Min/Max ESS/sec)
        ├── STAN2.stan                 Stan SRM, "STAN2" formulation (shared with fit_comparison.py)
        ├── original_srm.stan, original_srm   copy of the top-level original formulation
        └── benchmark_results_{cpu,gpu}.csv   raw results, N = 100/200/400/800
```

## What each comparison checks

- **`SIM.py`** — simulates block + individual + dyadic covariate data via R (`rpy2`),
  fits the same generative model three ways: BayesForge with parameters in "long"
  format, BayesForge with parameters in "wide" format, and R's `STRAND` package.
  Confirms all three recover consistent posteriors on identical data — the
  long/wide split is specifically a check that BayesForge's two equivalent
  parameterizations of the same model agree with each other, not just with STRAND.
  Header comment: `Tested on linux` — needs `rpy2` + a working R install with
  `STRAND` on `PATH`, not portable to a plain pip environment.

- **`fit_comparison.py`** — simulates data in pure Python/JAX, fits it with
  BayesForge and with two independently-written Stan models (`original_srm.stan`,
  `benchmark_suite/STAN2.stan`) via `cmdstanpy`. Produces `stan V BF.png`, a
  forest plot of all three posteriors overlaid per parameter.

- **`benchmark_suite/`** — not a correctness check, a performance one: wall-clock
  time and effective-sample-size-per-second across network sizes N = 100, 200,
  400, 800, for STAN2 / Stan Original / BayesForge (CPU and GPU). See its own
  `README.md` for how to reproduce the plots and regenerate the underlying CSVs
  (note: that README's reproduction commands reference a different machine's
  paths — `/home/sebastian_sosa/...` — adjust for your own environment).

## Dependencies

Python side (`fit_comparison.py`, `benchmark_suite/*.py`): `BayesForge`, `jax`,
`cmdstanpy` (+ CmdStan itself), `pandas`, `matplotlib`, `numpy` — same
BayesForge stack as `Examples/SRM/requirements.txt` and `Examples/Voice/requirements-stan.txt`
combined into one environment (both `bf` and `cmdstanpy` needed here, unlike
Voice which splits them).

`SIM.py` additionally needs R + `STRAND` + `rpy2`, and per its own comment is
Linux-only.

`test_strand.R`/`test_strand_2.R` need R + `STRAND`, no Python.
