## Supplementary figures and images for "BayesForge: A Bayesian Inference library for Python, R, and Julia"

### benchmark_efficiency.png

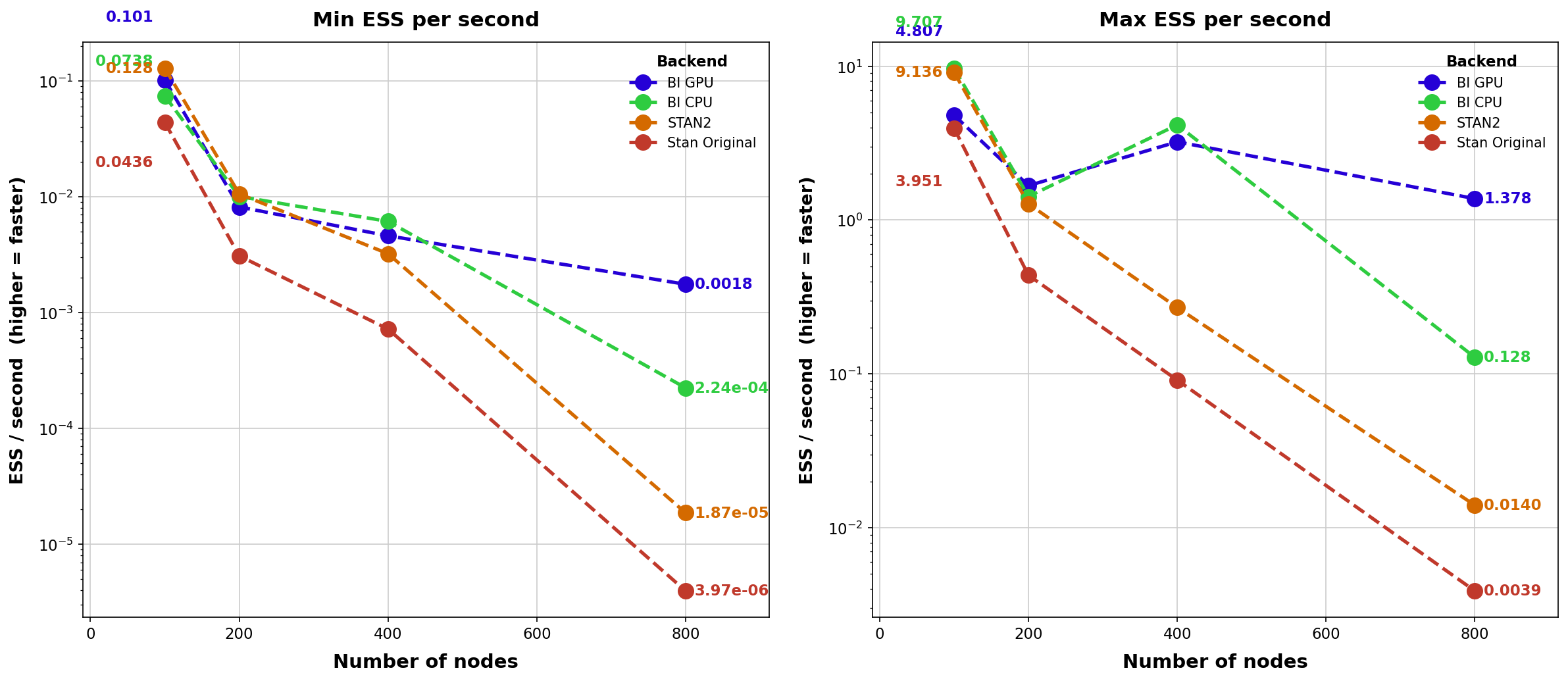

### benchmark_plot_cpu.png

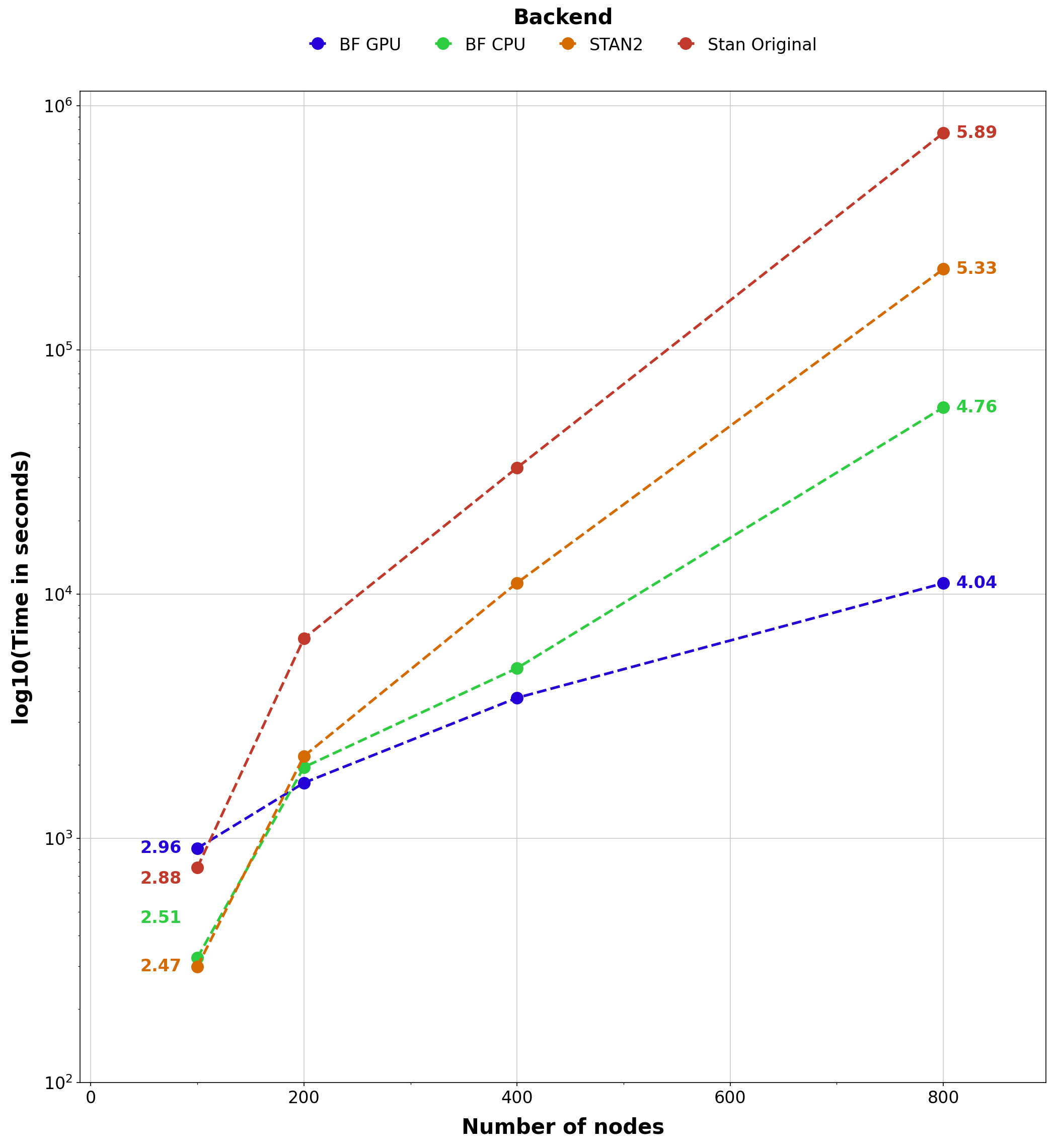

### density_L.png

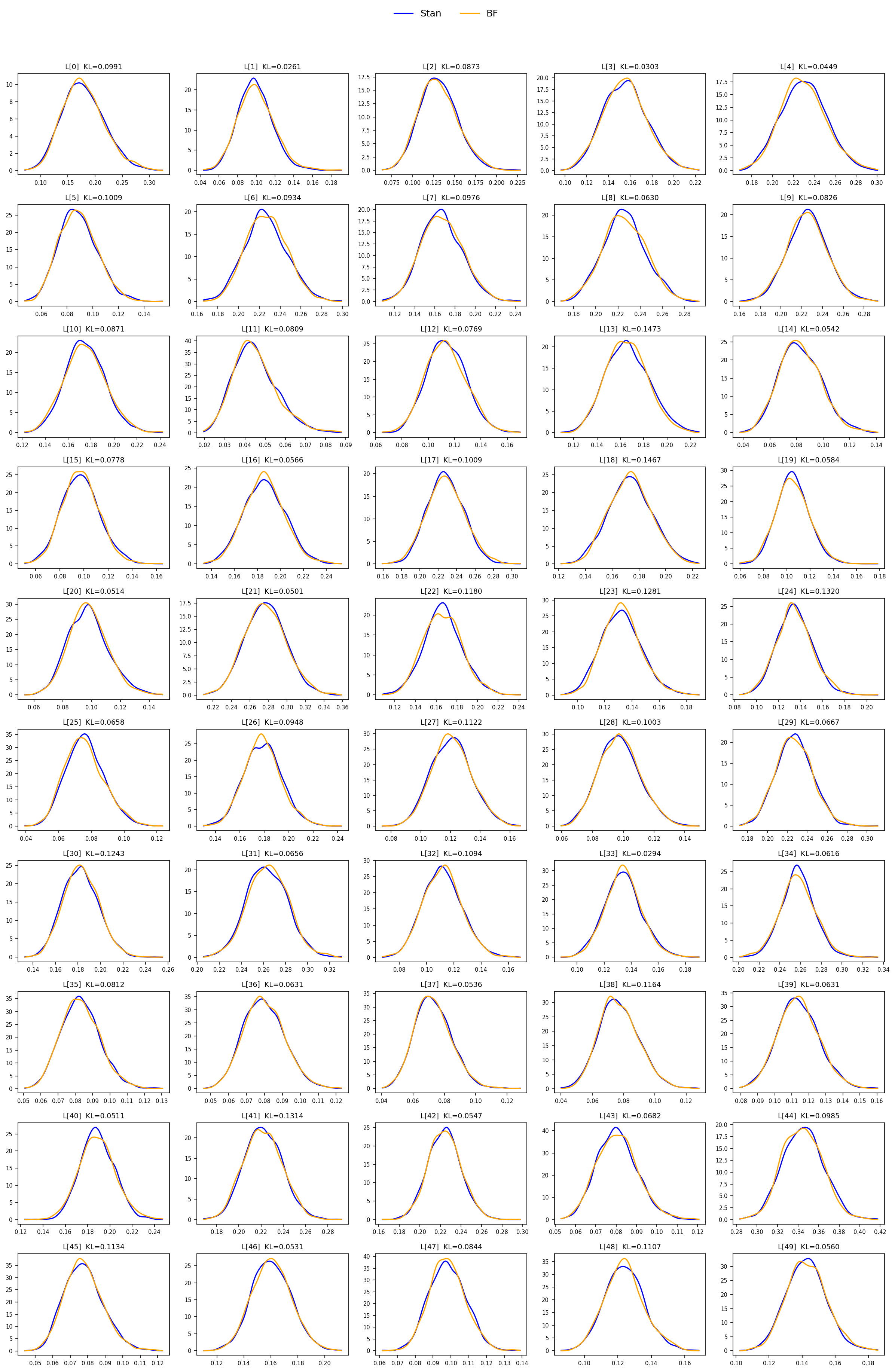

### density_p.png

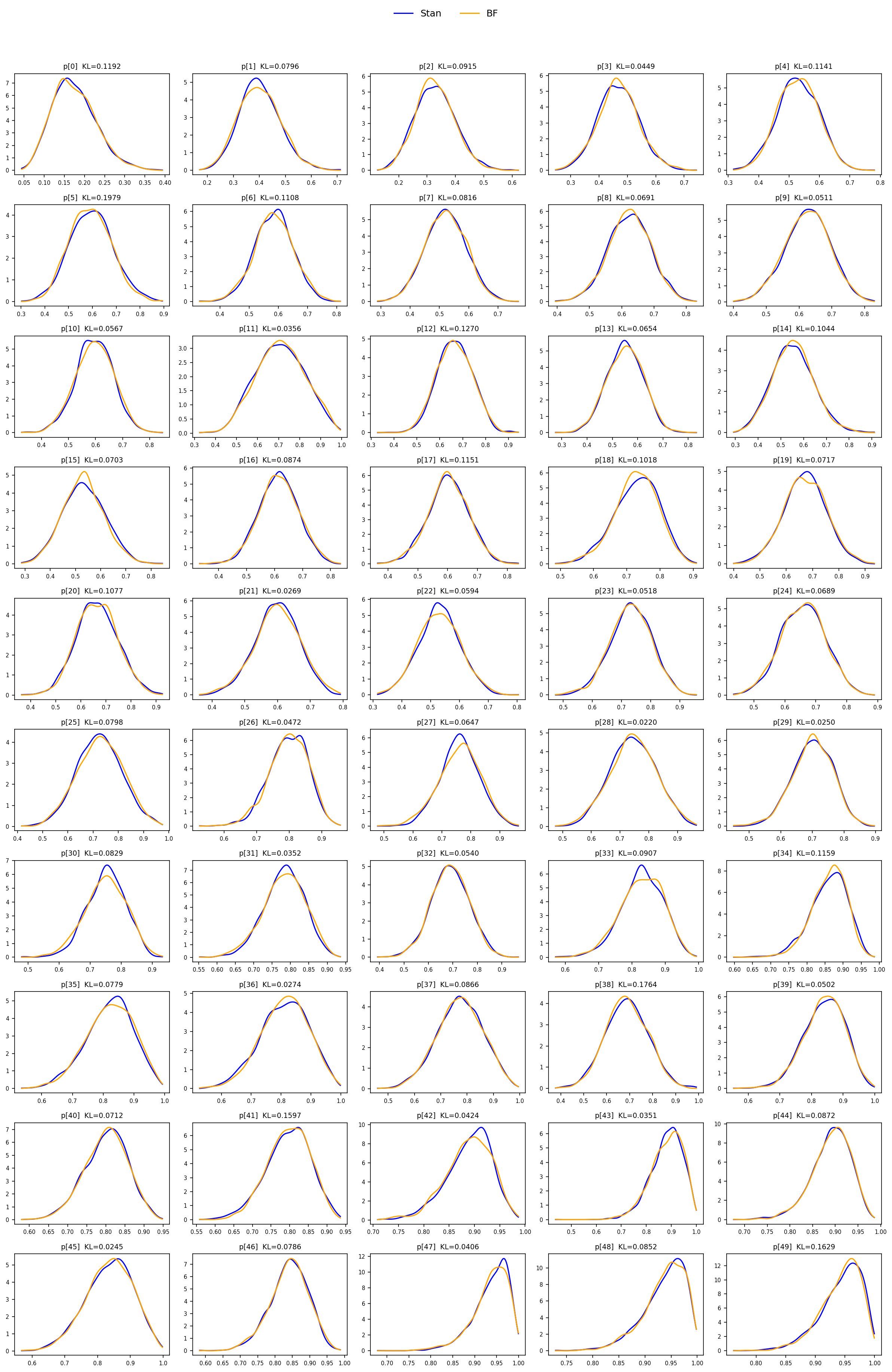

### density_R.png

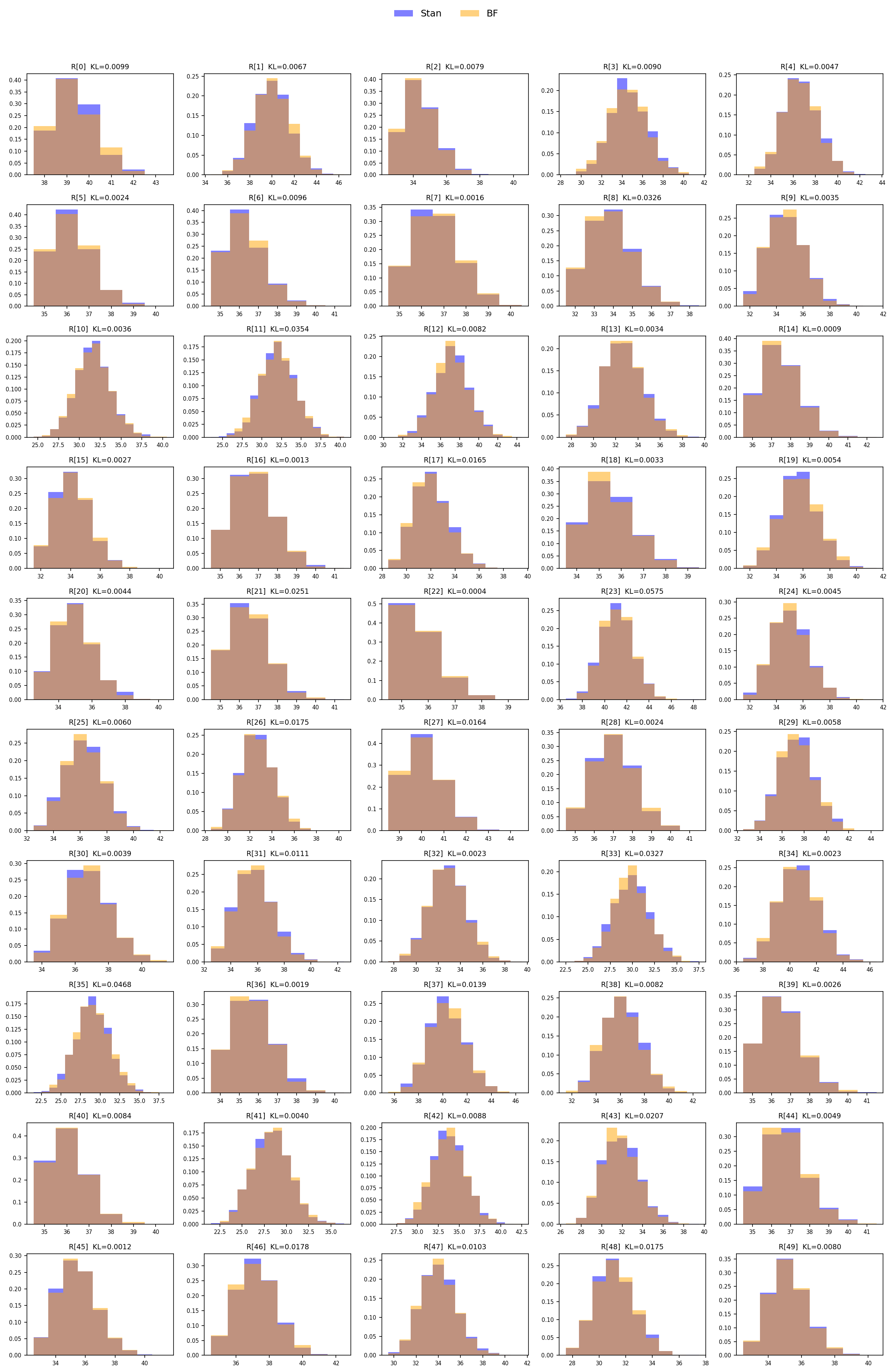

### diagnostics_table.png

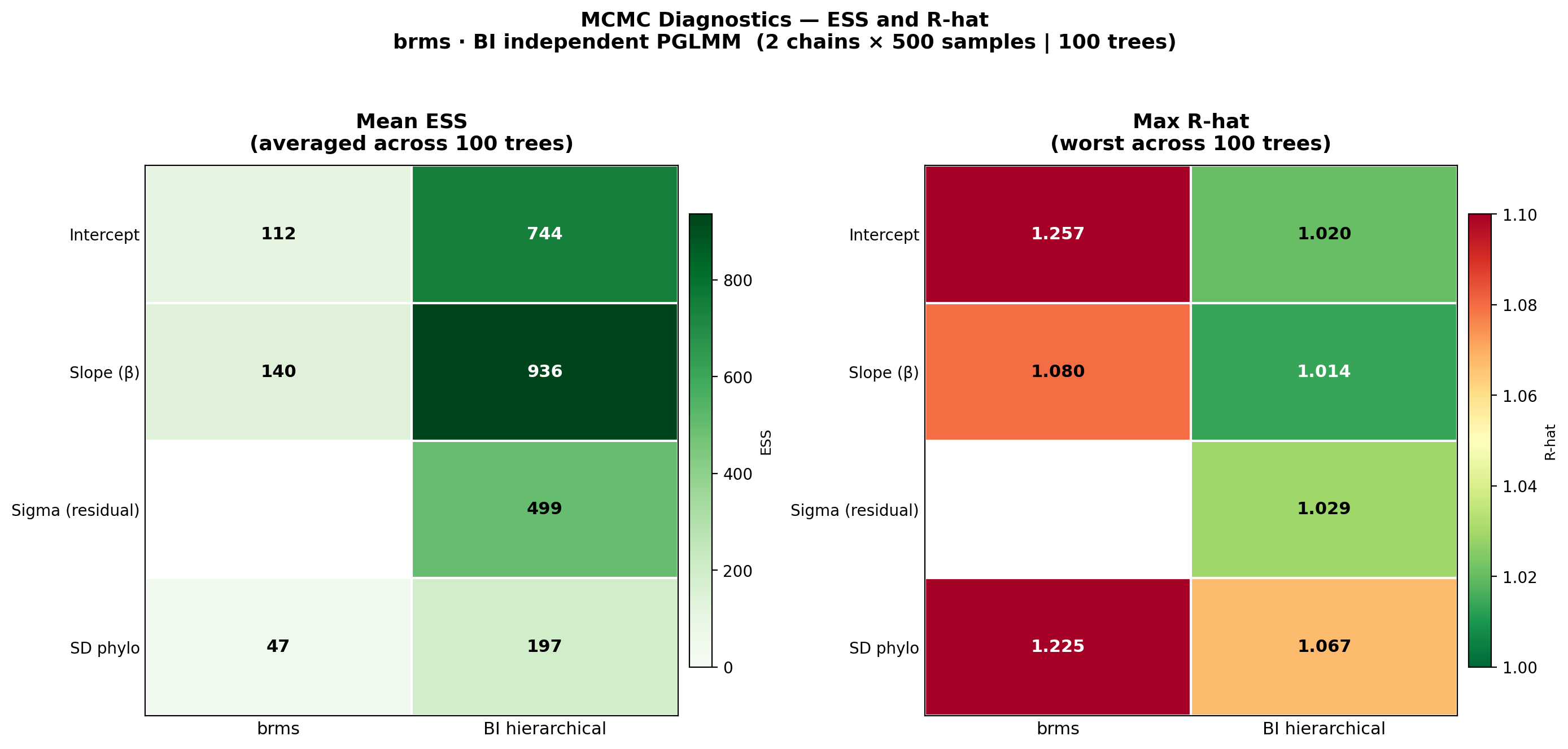

### efficiency_table.png

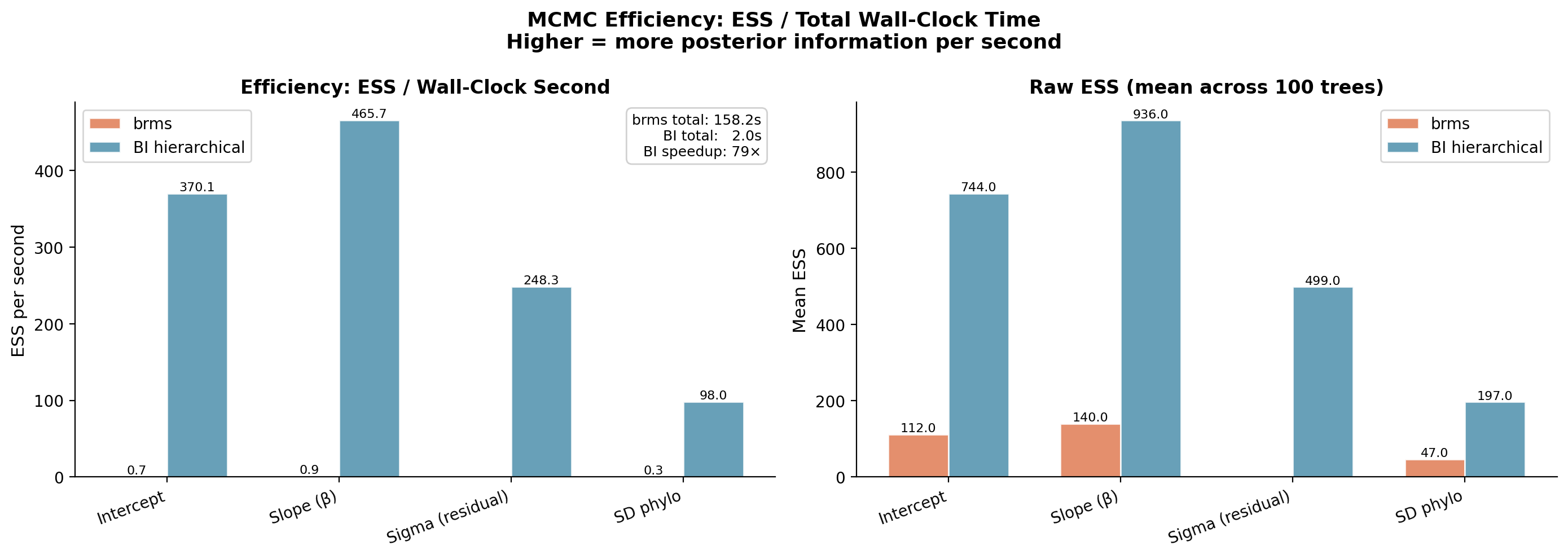

### forest_plot.png

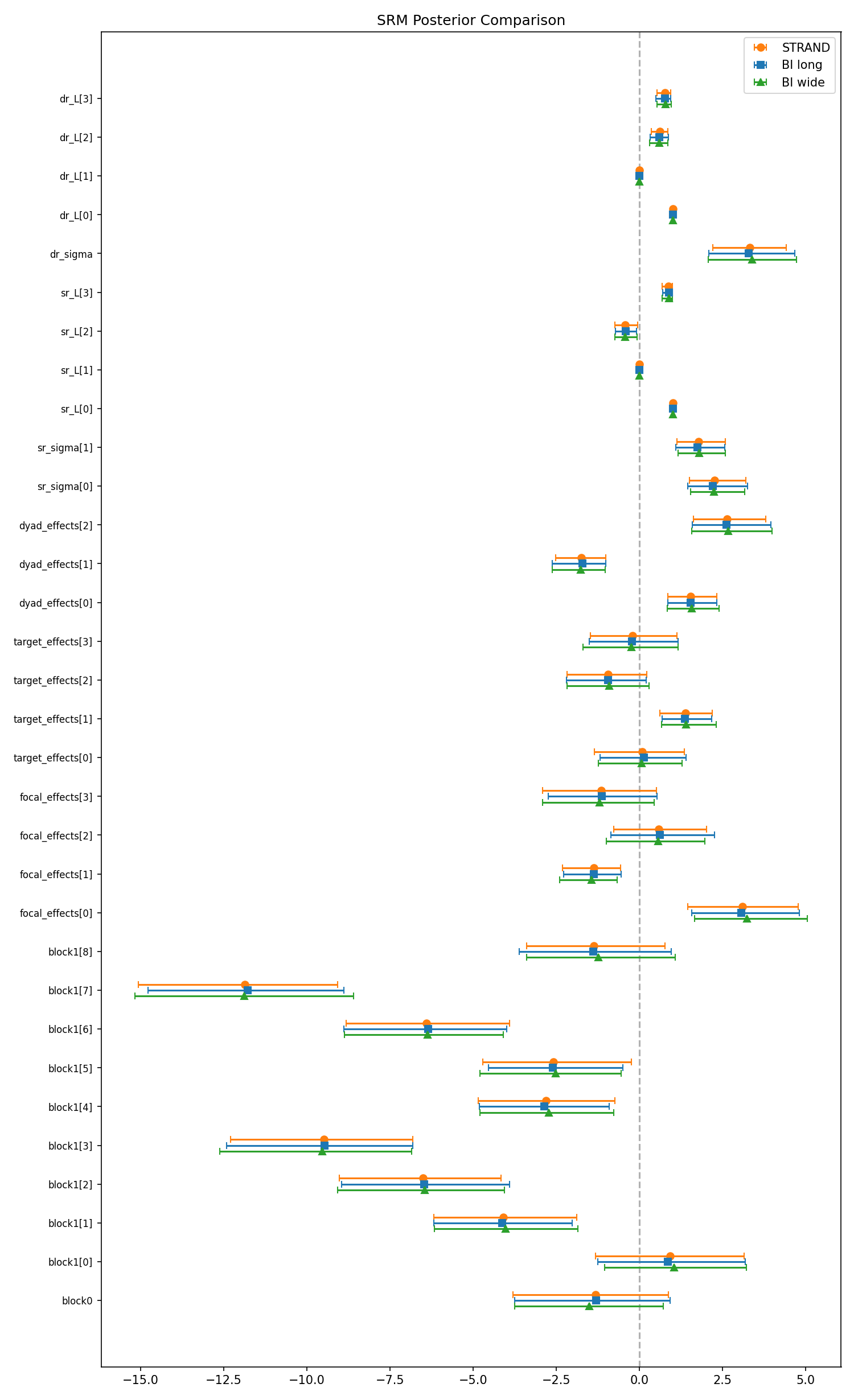

### forest_plot_nodal_re.png

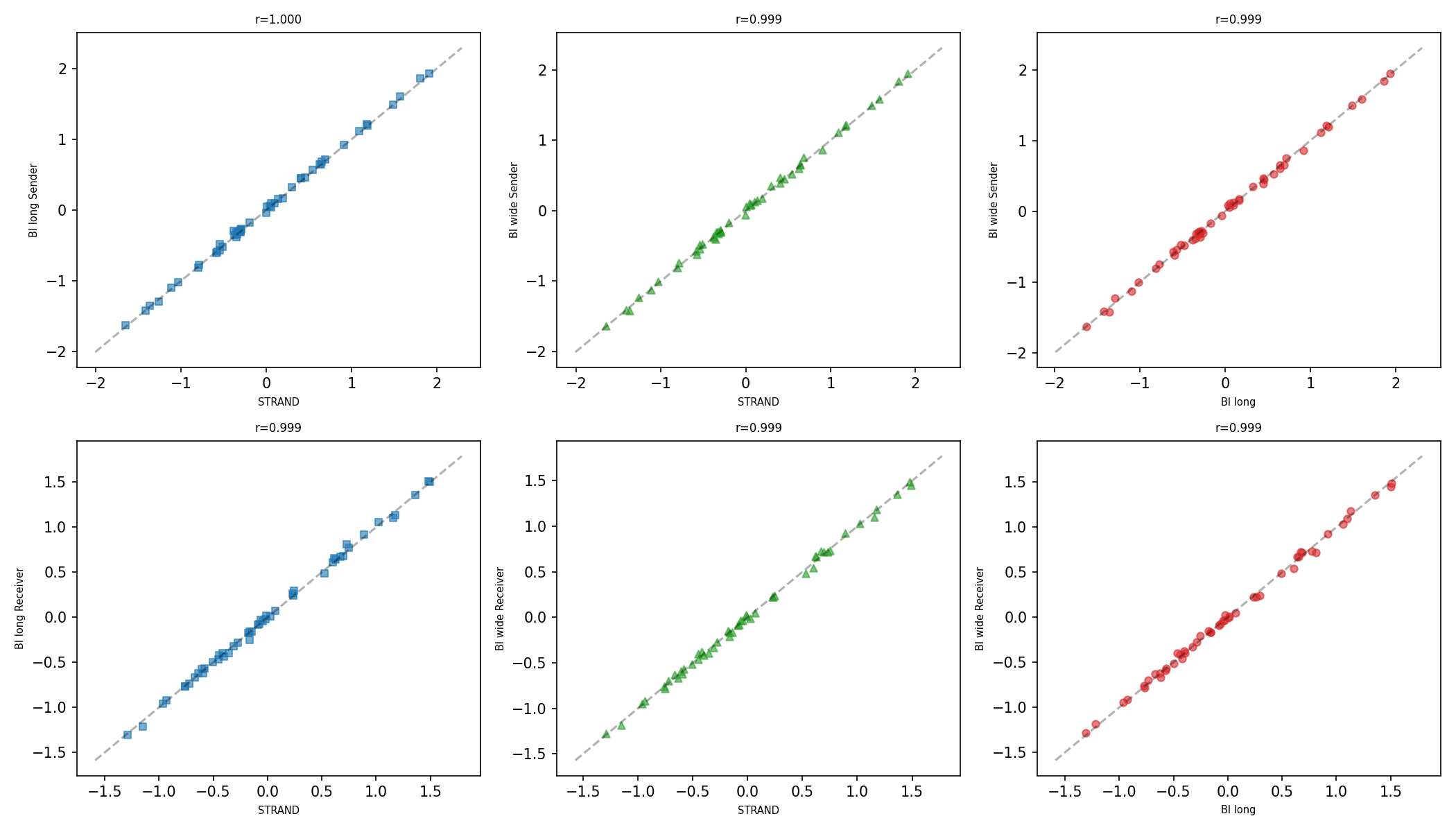

### forest_plot_nodal_re_scatter.png

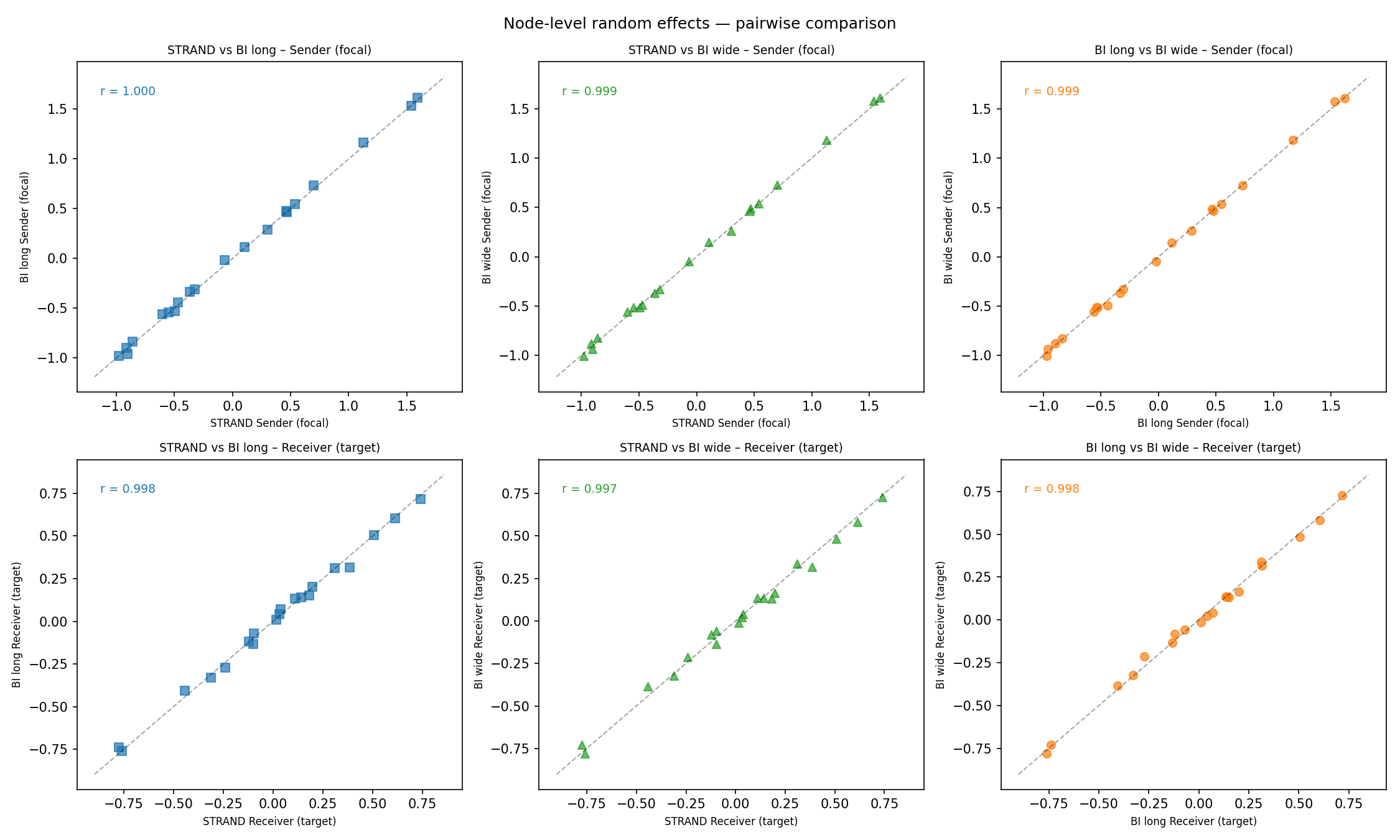

### posterior_densities.png

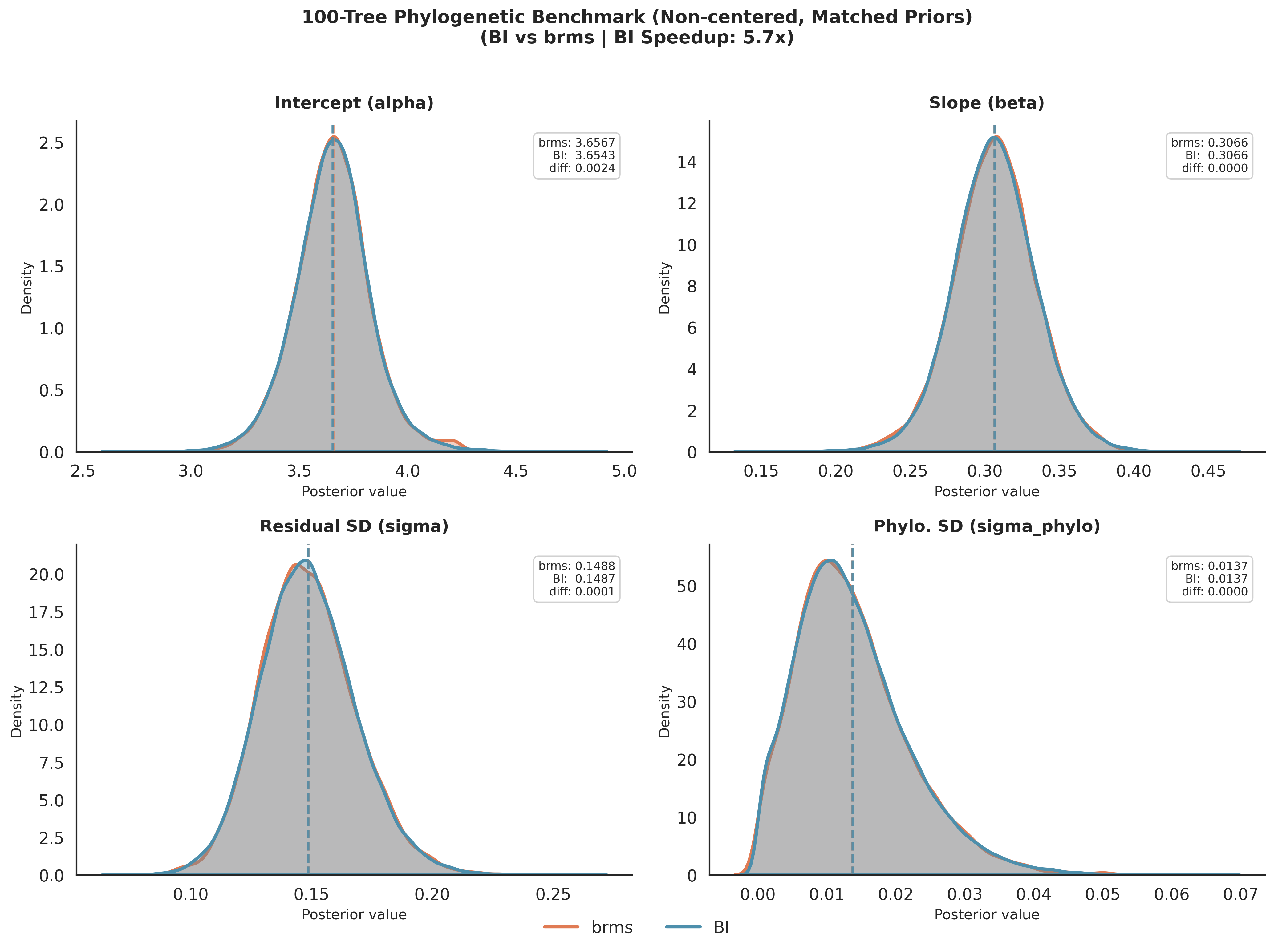

### srm_forest_plot.png

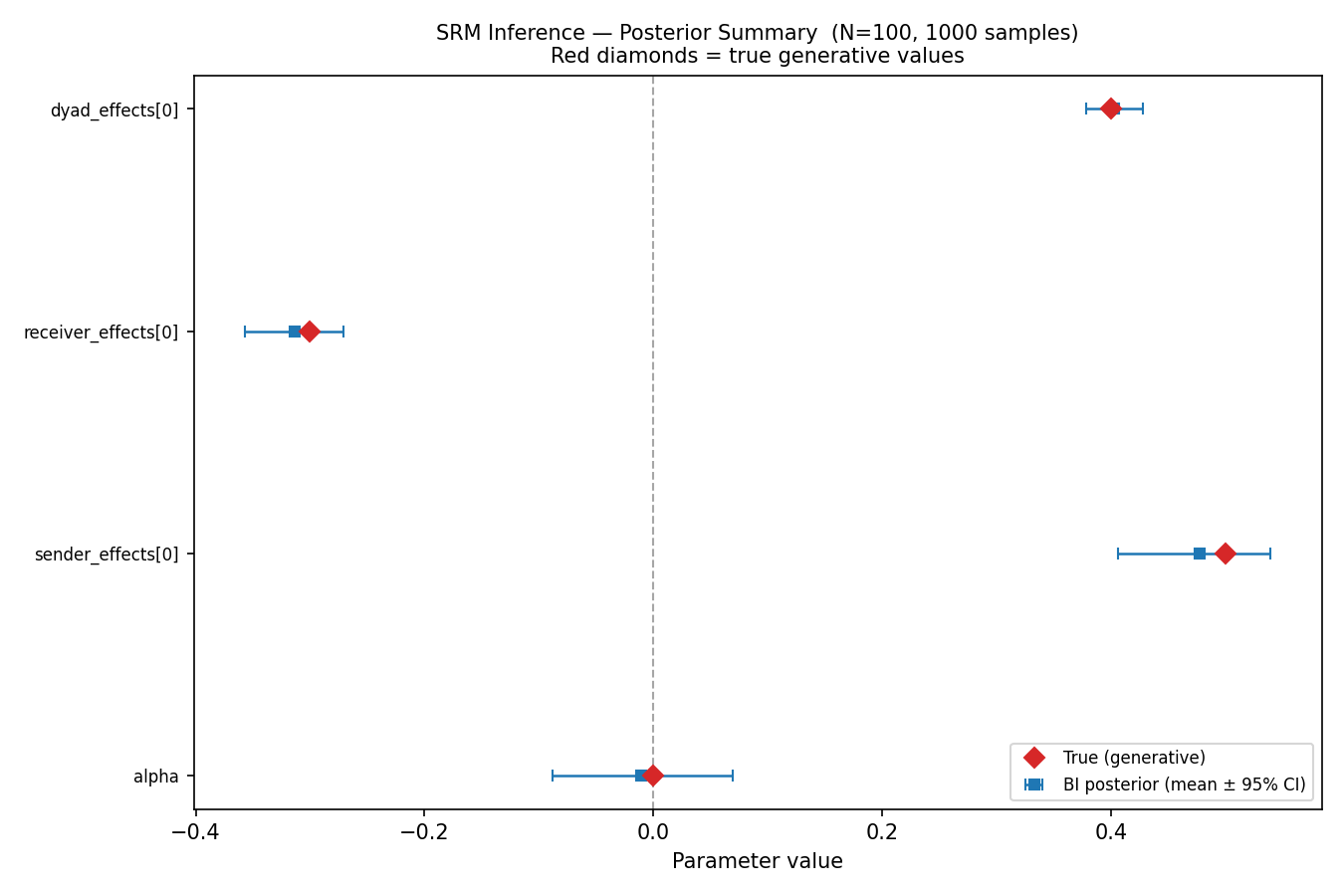

### srm_forest_plot_old.png

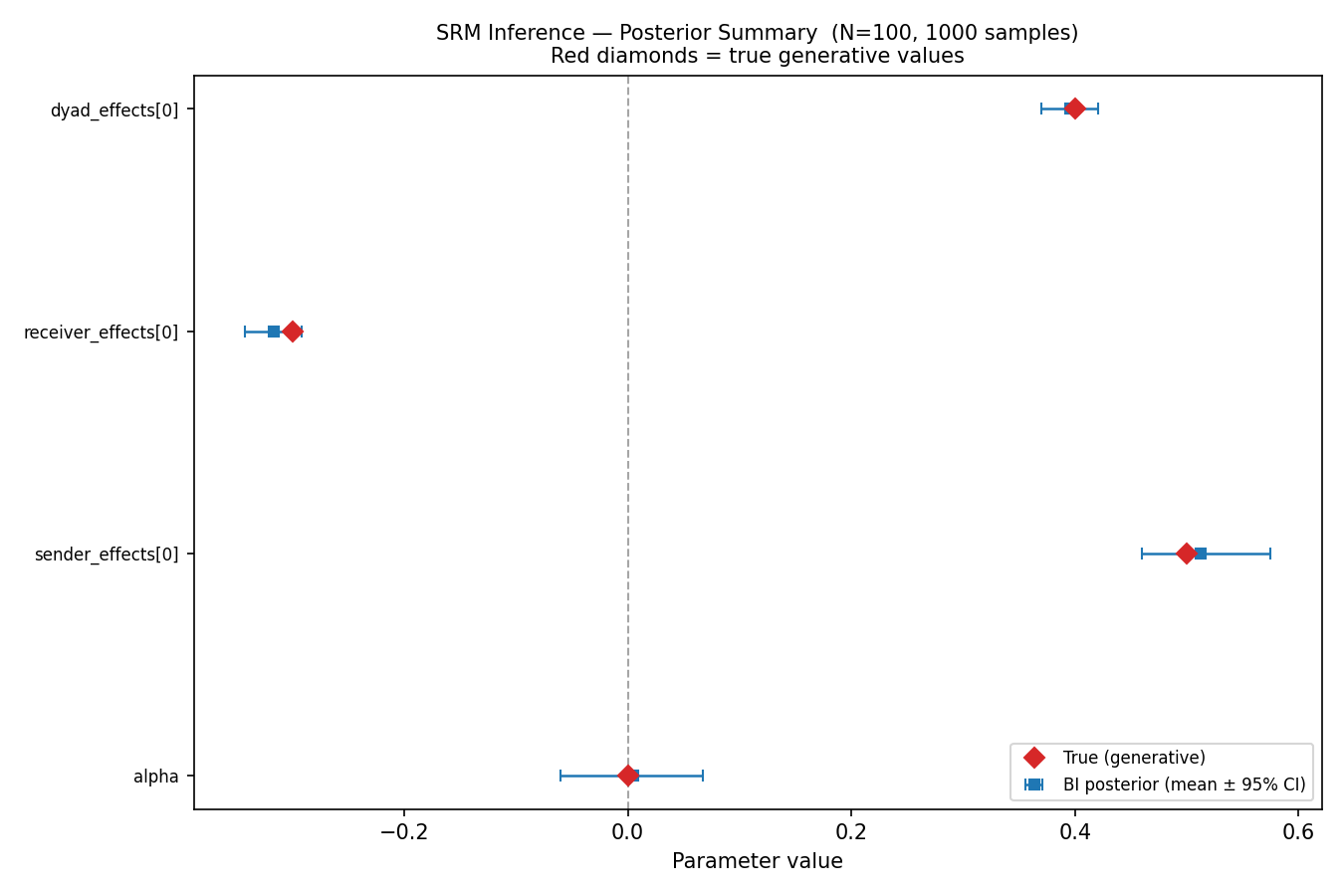

### stan V BF.png

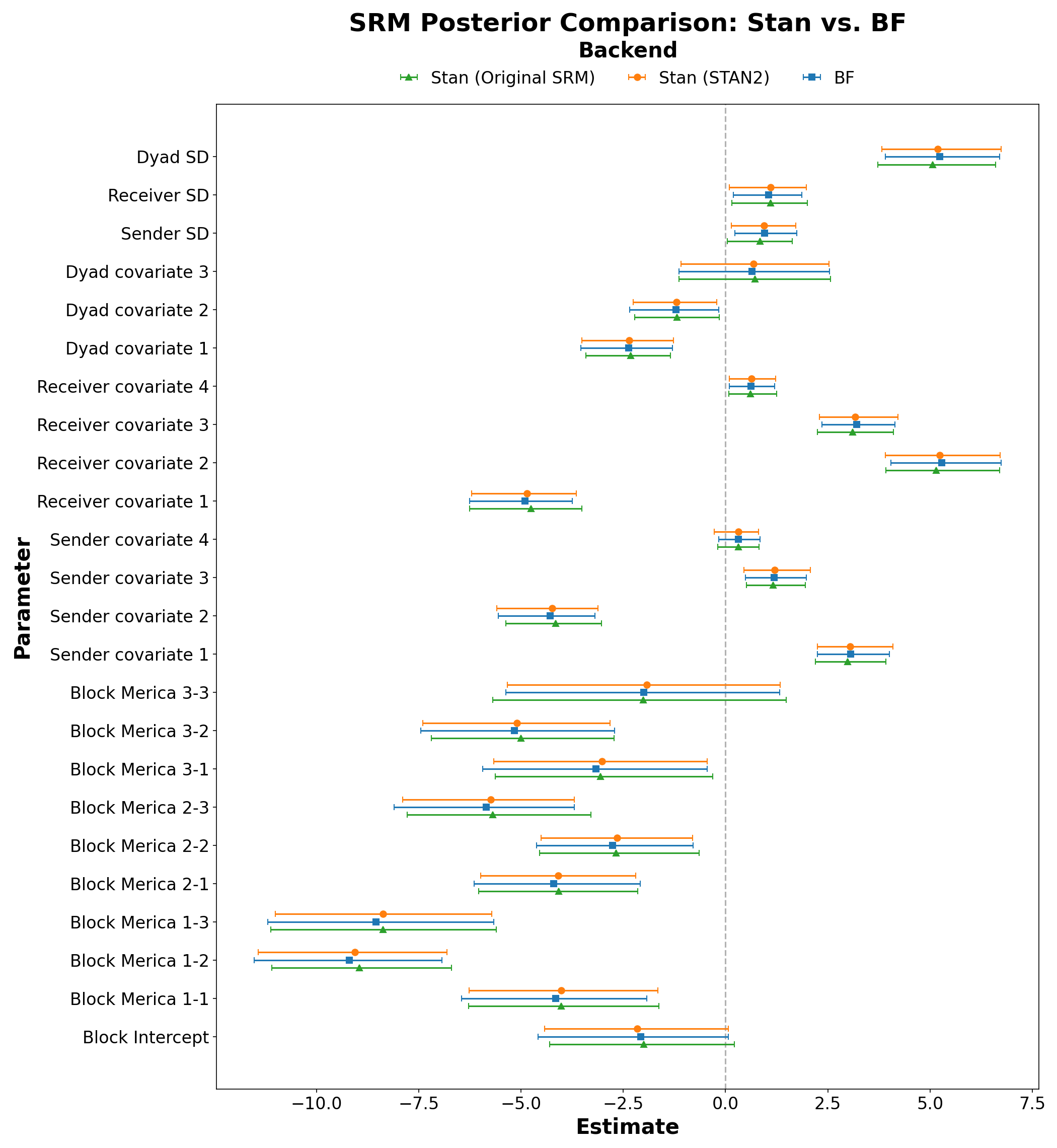
